# Heterogeneity Assessment and Protein Pathway Prediction via Spatial Lipidomic and Proteomic Correlation: Advancing Dry Proteomics concept for Human Glioblastoma Prognosis

**DOI:** 10.1101/2024.10.22.619687

**Authors:** Laurine Lagache, Yanis Zirem, Émilie Le Rhun, Isabelle Fournier, Michel Salzet

## Abstract

Prediction of proteins and associated biological pathways from lipid analyses via MALDI MSI is a pressing challenge. We introduced “dry proteomics,” using MALDI MSI to validate spatial localization of identified optimal clusters in lipid or protein imaging. Consistent cluster appearance across omics images suggests association with specific lipid and protein pathways, forming the basis of dry proteomics. The methodology was refined using rat brain tissue as a model, then applied to human glioblastoma, a highly heterogeneous cancer. Sequential tissue sections underwent omics MALDI MSI and unsupervised clustering. Differentiated lipid and protein clusters, with distinct spatial locations, were identified. Spatial omics analysis facilitated lipid and protein characterization, leading to a predictive model identifying clusters in any tissue based on unique lipid signatures and predicting associated protein pathways. Application to rat brain slices revealed diverse tissue subpopulations, including successfully predicted cerebellum areas. Similar analysis on 50 glioblastoma patients confirmed lipid-protein associations, correlating with patient prognosis.

**GRAPHICAL ABSTRACT:** 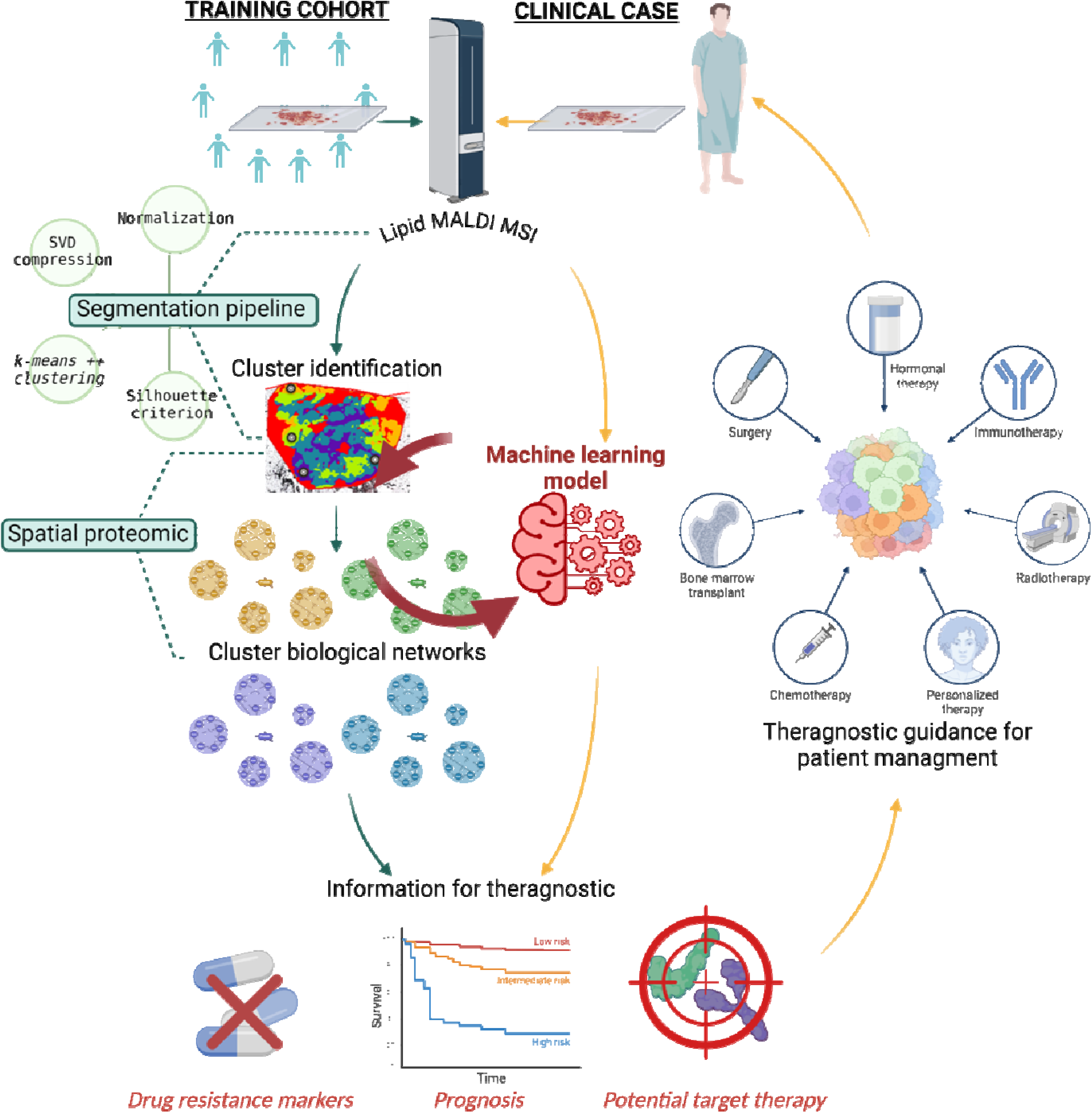

## INTRODUCTION

Since the gap between mass spectrometry imaging (MSI) and proteomics has been bridged by the development of spatially resolved proteomics guided by MALDI MSI^1–5^, the next challenge was to perform multi-omics analyses at the spatial level^6–11^. Nevertheless, there are still developments to be performed to correlate from lipid MSI data, proteins and lipid networks to retrieved functions. Multi-omics MSI is particularly valuable for the analysis of heterogeneous biological samples, such as brain or tumours, which consist of different cell types and regions with distinct molecular composition and function^3^. Indeed, tumour heterogeneity is a significant and growing area in cancer research. An overview on tumoral heterogeneous proteome is subsequently linked to therapeutic, allowing drug resistance analysis and optimized treatment guideline proposal, tending to personalize medicine strategy. However, the complex nature of protein annotation and the lack of standardized methodologies pose challenges to the effectiveness of MALDI-MSI data analysis, especially in multi-omics clinical research. The interpretation and integration of the vast amount of data generated by these technologies remains a significant limitation^12^. Extracting meaningful insights from complex datasets therefore requires sophisticated computational approaches and bioinformatic analysis^13^. MALDI MSI data analysis involves pre-processing and processing stages, preparing them for subsequent statistical analysis. Reduction techniques, like PCA (Principal Component Analysis)^14,15^, t-SNE (t-distributed Stochastic Neighbour Embedding)^16,17^, or NNMF (Non-negative matrix factorisation)^18,19^, are particularly useful for exploring the spatial distribution of molecular features in MALDI MSI data^20,21^. In addition, the combination of MSI and machine learning methods is widely used in the processing step to effectively extract the essential information contained in complex MSI data. The emergence of segmentation methods, such as bisecting *k*-means, hierarchical clustering and *k*-means clustering^22,23^, provides valuable insights from complex data like meaningful regions corresponding to biological features in heterogeneous sample. However, choosing the right number of *k*-clusters is not straightforward, limiting biological conclusions^22,23^. The common method involves performing *k*-means clustering for different k values (2<*k*<*k*_max_) and calculate the distances between clusters. The aim is to find the optimal *k* that minimizes intra-class distances while maximizing inter-class distances. Several statistical indices, called criterions, have been developed for this purpose^24,25^ .

Here, we introduce the concept of dry proteomics, an automated procedure capable of identifying heterogeneous clusters of biological samples according to their lipid signature, thought lipid MALDI MSI, and automatically providing their associated protein data without any proteomic experiments. The development of this machine learning method required overcoming several challenges (see, Graphical Abstract).The central hypothesis was that if a cluster appeared identical in both lipid and protein images, it should possess lipids and paired proteins related to a specific biological pathway, like a unique barcode that allows one cluster to be distinguished from others. Thus, the correlation between lipids and proteins in a biological network, within different clusters, forms the basis of dry proteomics. The data processing workflow was first developed on lipid, protein, and peptide MSI datasets performed on rat brain (RB) tissue. We succeeded in building a segmentation pipeline, consisting of Singular Value Decomposition (SVD) data compression pre-processing and *k*-means++ segmentation processing steps. The integration of the silhouette criterion allowed to optimize and automate the optimal number of clusters finding for MSI analysis, corresponding to the sample heterogeneity. The next step was to develop a prediction model that could blindly identify the different RB clusters from a lipid MS image according to their spectral fingerprint. The prediction model was complemented by discriminative lipid and protein identifications for each cluster, forming a dry proteomic reference dataset for RB tissue section.

Finally, the dry proteomics concept is a simple and rapid procedure, as the user only needs to perform lipid MALDI MSI to automatically identify the heterogeneous clusters present in a sample and obtain their specific proteome. The development of this tool is aimed at clinical application for patient therapeutic guidance. Indeed, the protein information provided by the dry proteomics process can be related to drug resistance, potential therapeutic target or patient survival, which could help the oncologist to propose a therapeutic guideline adapted to the patient’s tumour. In this way, the ultimate phase of presented research involved the application of this innovative concept to intricate and heterogeneous pathology samples, particularly human Glioblastoma ^26–28^. In addition, by applying the dry proteomics workflow, correlation between predicted protein and patient survival outcome information allowed to establish a robust model for glioblastoma patient survival prediction. This crucial validation step not only enhances confidence in the reliability of this approach but also holds significant promise for advancing personalized medicine strategies in the management of this challenging disease. Indeed, the assessment of heterogeneity, whether intra or interpatient, is pivotal in personalized medicine, as it allows for the identification of unique molecular profiles that can inform tailored treatment strategies for individual patients.

## EXPERIMENTAL PROCEDURES

### Experimental Design and Statistical Rationale

For MALDI imaging and spatial omics development studies nD=D3 male Wistar rats were sacrificed. All the experiments were performed in biological triplicate to ensure data reproducibility. For the proteomic statistical analysis of conditioned media, as a criterion of significance, we applied an ANOVA significance threshold of p-value ≤ 0.01, and heat maps were generated. Normalization was achieved using a Z-score with matrix access by rows. To assess the statistical significance of biomarkers for lipids MSI biomarkers, a non-parametric Kruskal-Wallis test was employed. Bonferroni corrections were applied to adjust p-values for multiple comparisons. Values are presented as medians and visualized through scatter boxplots.

A retrospective cohort of 50 fresh frozen (FF) glioblastoma tissues was obtained from the Pathology department of Lille Hospital, France. A prospective cohort of 50 FF glioblastoma tissues were also included in this study. 50 patients with newly diagnosed glioblastoma were prospectively enrolled between September 2014 and November 2018 at Lille University Hospital, France (NCT02473484). This research complies with all relevant ethical regulations. Approval of the study protocol was obtained from the Lille Hospital research ethics committee (ID-RCB 2014-A00185-42) before the initiation of the study. The study adhered to the principles of the Declaration of Helsinki and the Guidelines for Good Clinical Practice and is registered at NCT02473484. Informed consent was obtained from patients. Participants did not receive any compensation. According to the French Public Health Code and in application of the General Data Protection Regulations, all patients had been informed at the time of care that their standard clinical and biological data could be used for research purposes regarding the retrospective analysis of FF samples, and none had expressed his opposition. Regarding the prospective collection of samples, each patient’s informed consent for the collection and publication of clinical and biological data was obtained at the time of hospitalization prior to surgical intervention^27,28^. Tissue sections were subject to H&E coloration for histopathological analysis. The regions annotations were made by an anatomopathologist.

### Chemical products and Material

Water (H_2_O), ethanol (EtOH), acetic acid, acetonitrile (ACN) and methanol (MeOH) were obtained from Thermo Fisher Scientific (Courtaboeuf, France). 99% pure trifluoroacetic acid (TFA), α-cyano-4-hydroxycinnamic acid (HCCA), sinapinic acid (SA), 2,5-dihydroxybenzoic acid (2,5-DHB), aniline, formic acid (FA) and ammonium bicarbonate (NH_4_HCO_3_) were purchased from Sigma-Aldrich (Saint-Quentin Fallavier, France). The chloroform (CHCl_3_) was obtained from Carlo Erba Reagents (Val-de-Reuil, France). Porcine Trypsin Sequencing Grade was from Promega (Charbonnières, France).

Tissues were cut on a cryostat (Leica Microsystems, Nanterre, France). Indium Tin Oxide slides were purchased from LaserBio Labs (Valbonne, France), whereas the poly-lysine coated slides were from Epredia^TM^ (Braunschweig, Germany). The MALDI matrices and the trypsin were deposited on the tissue sections using the HTX M5-Sprayer™ (HTX Technologies, Carboro, NC, USA). Mass spectrometry imaging analyses were performed using the MALDI-TOF Rapiflex Tissuetyper (Bruker Daltonics, Bremen, Germany) equipped with the Smart Beam 3D laser. Spatial proteomic analysis were carried out through the utilization of chemical printer (CHIP-1000, Shimadzu, Kyoto, Japan) and the TriVersa Nanomate device (Advion Biosciences Inc, Ithaca, NY, USA). Samples were dried in a SpeedVac (SPD13DPA, Thermo Fisher Scientific, Waltham, Massachusetts, USA). nLC-MS/MS analysis were performed with TimsTOF Flex (Bruker) coupled to an EVOSEP One (EVOSEP).

### Sample preparation

Rat brains were obtained from our collaborator Dr. Dasa Cizkova (Institute of Neuroimmunology, Slovak Academy of Science, Bratislava). Male Wistar rats of adult age were sacrificed by CO_2_ asphyxiation and dissected. Brain tissues were frozen in isopentane at -50 °C and stored at -80 °C until use. Experiments on animals were carried out according to institutional animal care guidelines conforming to international standards and were approved by the State Veterinary and Food Committee of Slovak Republic (Ro-4081/17-221), and by the Ethics Committee of the Institute of Neuroimmunology, Slovak Academy of Science, Bratislava. For this study, FF rat brain tissues were cut using a cryostat at -20°C. All sections were obtained at the same time and stored at - 80°C until their use. Rat brain sagittal 12 µm sections were prepared, to finally reach 22 batch of 4 consecutive sections. Tissues were fixed on ITO slides and respectively intended to: back-up, lipid in negative and positive mode imaging, protein imaging and peptide imaging in positive mode^29,30^.

10 others consecutive rat brain sagittal sections of 12 µm were mounted on poly-lysine coated slide for lipid analysis carried out by SpiderMass technology. Three consecutive another 20 µm sections were fixed on polylysine coated slide for spatial proteomic analysis.

Finally, 3 different rat brain sagittal 12 µm section were fixed onto ITO coated slide as a validation cohort for the lipid predictive model.

For the analysis of horizontal rat brain tissues, 4 consecutives sections were prepared for multi-omics MSI analysis as describe bellow, followed by another consecutive sections for spatial proteomic analysis. This schema was repeated on 4 different rat brains.

### Lipid MALDI MS imaging

Tissues were dried in a desiccator before a matrix deposition. Norharman was used as MALDI matrix for positive and negative lipid imaging. The matrix was deposited at 7 mg/mL in CHCl_3_: MeOH (2:1, *v/v*). The HTX parameters for norharman spray were: spray at 30°C with 10 psi pressure, a pattern CC, a flow rate of 0.1 mL/min, a velocity of 1200 mm/min, for 12 passages with 2 mm track spacing. Lipid images were performed on the MALDI-TOF Rapiflex Tissuetyper mass spectrometer. The spectra were acquired within the *m/z* 200-1200 range in positive ion mode and the *m/z* 400-1500 range in negative ion mode. All data were performed in the delayed extraction reflectron mode with an average of 300 laser shots per pixel for a spatial resolution of 50 µm. The laser energy was set around 60 % and the voltages of the ion source were 20 kV and 11 kV for the lens. Same protocol was applied for 10 µm lipid imaging.

Other images were performed with DHB matrix in positive ion mode. The matrix was deposited at 10 mg/mL in MeOH: TFA 0.1% (7:3, *v/v*). The HTX parameters for DHB spray were: spray at 75°C, tray at 55°C, with 10 psi pressure, a pattern CC, a flow rate of 0.1 mL/min, a velocity of 1200 mm/min, for 8 passages with 2 mm track spacing. Lipid images were performed on the MALDI-TOF Rapiflex Tissuetyper mass spectrometer. The spectra were acquired within the *m/z* 200-1200 range in positive ion mode. All data were performed in the delayed extraction reflectron mode with an average of 300 laser shots per pixel for a spatial resolution of 50 µm. The laser energy was set around 85 % and the voltages of the ion source were 20 kV and 11 kV for the lens.

### Protein MALDI MS imaging

Tissues were vacuum dried before being subjected to delipidation using sequential baths of EtOH: H_2_O (70:30, *v/v*) for 30 s, EtOH 100% for 30 s, Carnoy solution (EtOH/Chloroform/Acetic acid, 3:6:1, *v/v/v*) for 2 min, EtOH 100% for 30 s, TFA 0.1%/H_2_O for 30 s and EtOH 100% for 30 s. After drying the sections, SA-Aniline (SA- ANI) MALDI matrix was deposited on tissue. SA-Aniline was prepared by dissolving sinapinic acid matrix at 10 mg/mL in ACN/TFA 0.1% (50:50, *v/v*) and adding 24.3 µL of aniline. The HTX parameters included a temperature of spray at 75°C with 10 psi pressure, a pattern CC, a flow rate of 0.1 mL/min, a velocity of 1100 mm/min, a temperature of tray at 55°C, for 8 passages with 2 mm track spacing. The slides were analyzed on the MALDI-TOF Rapiflex Tissuetyper mass spectrometer. MS spectra were acquired in the positive linear delayed extraction mode, on the *m/z* 2400-30,000 range with an average of 700 laser shots per pixel and at a spatial resolution of 50 µm. The laser energy was set around 90 %. The voltages of the ion source were 20 kV and 9 kV for the lens.

### Peptide MALDI MS imaging

For peptide imaging, the slides were dried and delipidated using a similar protocol as for protein MS Imaging. The tissue sections were then submitted to trypsin digestion. The tryptic digestion was performed by applying trypsin (40 µg/mL in NH_4_HCO_3_ 50 mM). The HTX parameters included a temperature of spray at 65°C with 10 psi pressure, a pattern CC, a flow rate of 0.1 mL/min, a velocity of 1100 mm/min, for 12 passages with 2 mm track spacing. Once the trypsin was deposited the slides were incubated overnight at 56°C in a humidified box containing MeOH/H_2_O. The slides were then dried under vacuum over the next day. An HCCA-aniline matrix was deposited by the HTX M5-Sprayer. Briefly, 43.2 µL of aniline were added to 5 mL of a solution of 10 mg/mL HCCA dissolved in ACN/TFA 0.1% (7:3, *v/v*). Slides were analyzed on a MALDI-TOF Rapiflex. Spectras were obtained in the positive delayed extraction reflector mode analysis, with a mass range of 700-3200 *m/z*, and averaged from 500 laser shots per pixel for a spatial resolution of 50 µm. The laser energy was set around 40 %. The voltages of the ion source were 20 kV and 11 kV for the lens.

### Multi-Omics MSI segmentation

The raw MALDI MSI data for lipids in both ionization modes, peptide and protein data were initially converted into the imzML format^31^ using SCiLS lab software. Subsequently, the imzML converter, version 1.3.3, was employed to import these datasets into MATLAB R2019a. It’s worth noting that MSI data is characterized by high dimensionality, often reaching sizes of up to 100 GB per image. This magnitude makes it infeasible to analyze such data. To address this issue and prevent data loss using peak list generation, data compression was implemented as a preprocessing step before segmentation. Several data reduction (compression) algorithms were explored, including t-SNE (t-distributed Stochastic Neighbor Embedding), NNMF (Non-Negative Matrix Factorization) and SVD (Singular Value Decomposition). For the segmentation process, the *k*-means++ algorithm was utilized, implemented as the ‘*k-*means’ function in the MATLAB Statistics Toolbox. *K*-means++ offers an improved initialization of centroids, enhancing the quality of clustering^32^. The cosine distance metric was employed to calculate the cosine angle between two spectra for quantifying the similarity. For visualization, each cluster’s pixels are uniformly assigned a specific color, facilitating the creation of a segmentation map. This map delineates the cluster or region of interest to which each pixel (spectrum) belongs. To estimate the right numbers of clusters, the Silhouette criterion was used. After predefining the number of clusters, the silhouette plot method was used to assess the stability of the clusters. The silhouette plot displays a measure of the proximity of each point in a cluster. This measure has a range (-1, 1). A value close to 1 indicates that the cluster is distant from neighboring clusters (the spectra are very compact within the cluster to which it belongs and distant from other clusters). A value of 0 indicates that the sample is very close to the decision boundary between two neighboring clusters (overlapping clusters). Negative values indicate that these samples may have been assigned to the wrong cluster^33^. Silhouette plot was calculated using the function silhouette in Matlab. Subsequently, each centroid within these clusters is thoughtfully exported in CSV format, ready for further in-depth analysis and exploration.

### Differential analysis between clusters

The centroids generated from the image segmentation were imported into Python using the panda’s library. All centroid data was structured into a data frame. A custom script was developed to automate the execution of a statistical test. This script iterates over all *m/z* variables, identifying ions that exhibited statistical significance between the regions of interest (ROIs). To enhance data quality, a peak picking algorithm was employed. Specifically, the find_peaks_cwt function from the Scipy library was utilized to effectively remove instrument noise. A nonparametric statistical test, the Kruskal-Wallis test with Bonferroni correction, was conducted. Only features deemed statistically significant, with a p-value equal to or less than 0.05, were retained. A manual step is added to isolate and retain only the mono-isotopic peaks. The seaborn library was utilized to generate corresponding box plots.

### Prediction model based on lipid MALDI imaging and associated proteins pathways

The previously developed pipeline^27^ served as the foundation for constructing the optimal model adapted to the dataset based on highest accuracy and F1-score. These predictive models are designed to classify new MSI-lipid samples pixel by pixel, or the centroid of clusters after segmentation. While models cannot directly predict protein pathways, clusters previously associated with detected proteins using spatially resolved proteomics can indicate these pathways. Therefore, a logical algorithm was integrated into the prediction process. When a model predicts clusters, it also highlights the associated pathways and the corresponding list of proteins.

The three selected models for both rat brain optimization and glioblastoma applications were Stochastic Gradient Descent (SGD)^34^, RidgeClassifier^35^ and Light Gradient Boosting Machine (LGBM)^36^. The **Table 1** summarize the performance of each model in both rat brain and glioblastoma analysis. In addition, LIME (Local Interpretable Model-agnostic Explanations) was used for each model to understand the decision-making process of the models and thus identify the molecules that contribute most to predicting each cluster. The highest-contributing molecules are considered potential biomarkers.

**Table 1:**
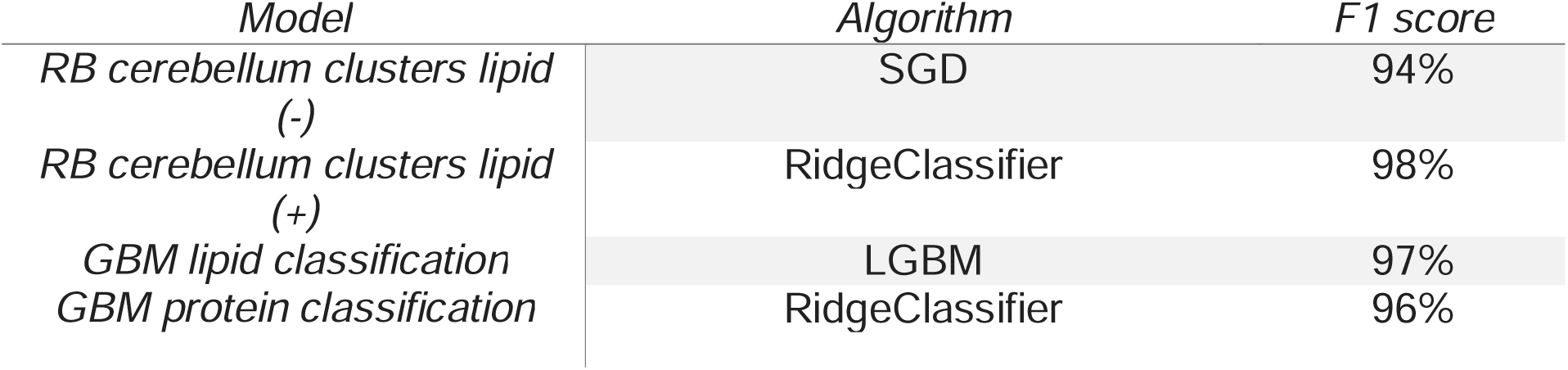
Model algorithms implication.

| Model | Algorithm | F1 score |
| --- | --- | --- |
| RB cerebellum clusters lipid (-) | SGD | 94% |
| RB cerebellum clusters lipid (+) | RidgeClassifier | 98% |
| GBM lipid classification | LGBM | 97% |
| GBM protein classification | RidgeClassifier | 96% |

### Lipid annotation by SpiderMass technology

The basic design of the instrument setup has been described in detail elsewhere^37^. In addition, here, the laser system used was an Opolette 2940 laser (OPOTEK Inc., Carlsbad, California, USA). The infrared laser microprobe was turned at 2.94 µm to excite the most vibrational band of water (O-H). The laser beam was injected into a 1 m reinforced jacketed fiber of 450 µm inner core diameter equipped at its extremity with a handheld including a focusing lens with 4 cm focal distance to get a 500 µm spot on the tissue. To aspirate and analyze the ablated material, a Tygon^Ⓡ^ tubing (Akron, OH, USA) is directly connected to Q-TOF mass spectrometer (Xevo, Waters, UK) through a REIMS interface. Each rat brain cerebellum clusters, observed by MSI, were analyzed by SpiderMass with four independent biological repetitions. Briefly, the laser was directly placed above the region of interest at the 4 cm focal distance. The laser energy was fixed to 4 mJ/pulse^38^. On each spot, three acquisitions of 10 repetitive laser shots (10 Hz) were performed which resulted in 3 individual MS spectra. The data were acquired in both negative and positive polarities, in the sensitivity mode over a *m/z* 100-2000 range. The previously identified discriminative ions were selected for MS/MS analysis with 0.1 *m/z* isolation window. MS/MS was performed using collision induced dissociation (CID) with argon as collision gas and a collision energy of 25 eV.

### Spatially resolved proteomics extraction

The different clusters identified by the segmentation process were submitted to spatially resolved proteomics. Each cluster was analyzed in triplicate from the same tissue section as describe bellow. A localized digestion was carried out by deposing a trypsin solution (40 µg/mL in NH_4_HCO_3_ 50mM), on a region of 500 µm^2^ of tissue (4 x 4 droplets of 200 µm in diameter), using CHIP-1000. The deposition method comprises approximately 1205 cycles per digestion spot, i.e., 3 hours of deposition, with a drop volume of 150 pL. Finally, each spot was digested with 0.112 µg of trypsin. Following the micro-digestion, each spot was extracted by liquid micro-junction using the TriVersa Nanomate device, with LESA (Liquid Extraction and Surface Analysis) parameters^1^. The tryptic peptides were extracted by performing 2 consecutive extraction cycles for three different solvents mixtures (TFA 0.1%; ACN/0.1% TFA (8:2, *v/v*); and MeOH/0.1% TFA (7:3, *v/v*)) for a total of 6 extractions. For each cycle, 2 µL of solvent was drawn into the tip of the pipette, of which 0.8 pL was brought into contact with the surface. 15 back and forth movements were performed to extract the peptides before collecting the solution in a recovery tube. All extracts were pulled in one tube and 50 µL of ACN were finally added before drying the samples in a SpeedVac. The samples were then stored at -20°C prior to nLC-MS/MS analysis.

### nLC-MS/MS bottom-up analysis

All sample analysis was performed on a timsTOF fleX mass spectrometer online coupled to an Evosep One nano-flow liquid chromatography system. Peptides were separated using an 8 cm x 150 µm C18 column with 1.5 µm beads and the 60 samples per day method from Evosep One. The mobile phases comprised 0.1% FA in water as solution A and 0.1% FA in ACN as solution B. To perform DIA analysis in PASEF mode^39^, one MS1 scan was followed by 10 dia-PASEF scans from *m/z* 100 to 1700. The ion mobility range was set to 1.42 and 0.65 V.s/cm^−2^. The accumulation and ramp times were specified as 100 ms. As a result, each MS1 scan and each MS2/dia-PASEF scan last 100 ms plus additional transfer time, and a dia-PASEF method with 22 dia-PASEF scans has a cycle time of 1.06s. The mass spectrometer was operated in high sensitivity mode, with a collision energy ramped linearly as a function of the ion mobility from 59 eV at 1/K_0_=1.6Vs.cm^−2^ to 20 eV at 1/K_0_=0.6Vs.cm^−2^. The ion mobility was calibrated with three Agilent ESI Tuning Mix ions (*m/z*, 1/K_0_: 622.02, 0.98 V.cm^−2^, 922.01, 1.19 V.cm^−2^, 1221.99, and 1.38 V.cm^−2^).

### Proteomic data analysis

DIA-NN version 1.8.1 was used to search DIA raw files and dia-PASEF files. A Rattus library was generated with the software parameters set as following: complete proteome of Rattus norvegicus from UniProt database (Release January 2024, 92958 entries), Trypsin protease with 2 missed cleavages and a maximum number of variable modification at 3, methionine oxidation as variable, peptide length range from 7 to 30, precursor charge range from 1 to 4, precursor *m/z* range comprised between 100 and 1700, fragment ion *m/z* range between 200 and 1700, 0.1% precursor FDR, protein inference set on ‘genes’, neural network classifier on single-pass mode, quantification strategy set on robust LC (high accuracy), RT-dependent cross-run normalization, and library generation fixed on the ‘IDs, RT & IM profiling’ ruban. Samples were interrogated according the resulting Rattus library with the same options. Data are available via ProteomeXchange with identifier PXD054488. Statistical analyses were carried out using Perseus software v2.0.5.0. ANOVA tests were performed with p-value ≤ 0.01 to be statistically significant and generate heat maps of differentially expresses proteins across sample. Gene Ontology (GO) analysis were performed using ClueGO^40^ with GO term database, on Cytoscape v3.10.2^41^.

## RESULTS

The main goal of this study was to develop a machine learning pipeline capable of automatically identifying tissue heterogeneity clusters from lipid MSI data and providing associated protein networks without requiring additional protein experiments. To this end, the first challenge was to demonstrate that identified clusters are specifically spatially localized by MSI, regardless of whether lipid or protein imaging is used. Following this idea, if a cluster is identical on these omics images, it should possess specific lipid and protein pathways, tending to the basis of the dry proteomics concept.

### Segmentation workflow development on RB cerebellum omics MSI

#### Clustering multi-omics MALDI MSI workflow optimization

The machine learning clustering processing was the first development to adapt a workflow for multi-omics MALDI MSI. This step was focused on RB cerebral lump, a model whose anatomical and molecular characteristics are already well referenced. For the latter, four main clusters are described (**Fig 1B**): the white matter (WM) and the grey matter (GM), composed of the molecular layer (ML), Purkinje cells and the granular layer (GL) (**Fig. S1**). The first aim was to demonstrate that these clusters could be observed with the same spatial localization in each omics image, using an adapted segmentation process script.

**Fig. 1:**
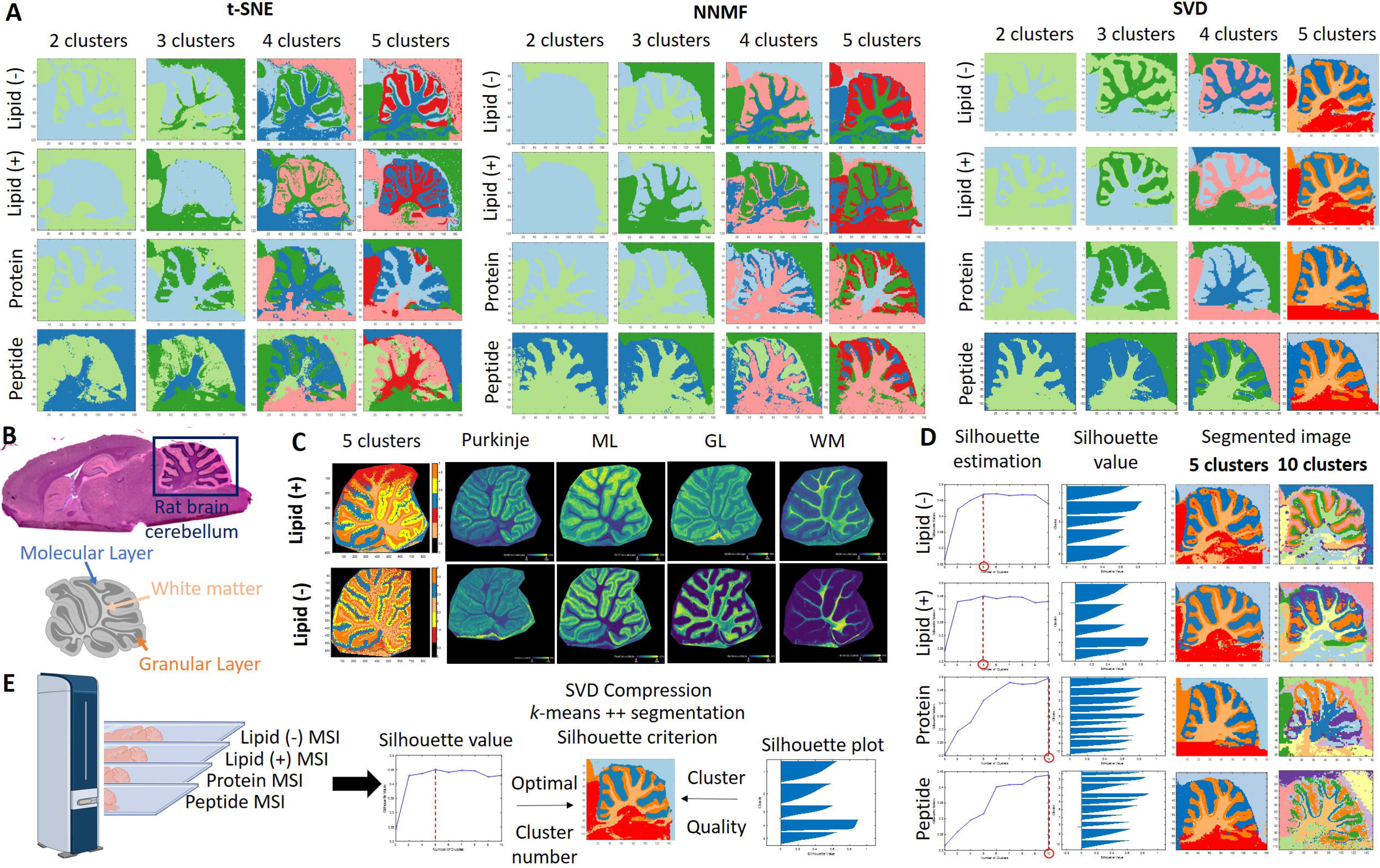
Omics MALDI MSI clustering procedure optimization on rat brain cerebellum. **A)** Comparison of t-SNE, NNMF and SVD data compression followed by *k*-means++ segmentation for 2 to 5 clusters applied to lipid negative mode, lipid positive mode, protein, and peptide MSI. **B)** Rat brain sagittal section HPS coloration and cerebellum annotations. **C)** Lipid MALDI MSI in negative and positive mode with 10 µm spatial resolution with image segmentation composed by 5 clusters, and ion spatial distribution specific to Purkinje cells, ML, GL and WM. **D)** Use of Silhouette criterion for the number of cluster estimation and each cluster value determination applied to lipid negative mode, lipid positive mode, protein, and peptide imaging. **E)** Optimal segmentation workflow developed on MATLAB integrating a SVD compression data with 10 principal components, combined with a *k*-means++ segmentation using a cosine score with a Silhouette criterion

For that, 22 RB sagittal sections were analyzed for lipid in negative (-) and positive (+) ion mode, while 12 slides were analyzed for protein and peptide, focusing on the RB cerebellum area. First, the MS spectra revealed different molecular fingerprints regarding WM, GL, and ML clusters for each molecular MSI (**Fig. S2**), confirmed for lipid (-) and protein data by t-SNE revealing clear separations of the different ROIs. On the contrary, the t-SNE obtained for lipid (+) and peptides did not show a clear separation of the different ROIs, which could predict difficulties for data processing of the latter.

To generate the most relevant segmented images, the image data was first analyzed on SCiLS lab software using RMS (Root Mean Square) normalization. The SCiLS software allows to play with different clustering parameters. Several segmentation methods were tested, including bisecting *k*-means, hierarchical clustering and *k*-means segmentation using correlation or Euclidean distance metrics. As shown in **Fig. S3**, the use of bisecting *k*-means and hierarchical clustering were ruled out due to the difficulty of interpreting the results for several reasons. First, the complexity of manually determining the desired number of clusters, which can be difficult in the case of a complex and unknown image. In addition, the spatial connectivity limitations of bisecting *k*-means do not adequately account for the connectivity between pixels in an image. This oversight can lead to segmentation discontinuities that undermine the overall accuracy and coherence of the segmentation process. *k*-means segmentation appeared to be more user-friendly, with multiple clusters defined subjectively. Unfortunately, it seems that poor centroid initialization led to insufficient clustering performance, rendering the segmentations of lipid, protein, and peptide images incomparable despite using the same number of clusters

To find a more transparent and robust strategy, data from SCiLS was imported into MATLAB software. To improve the previous clustering performance, the *k*-means++ segmentation algorithm with cosine distance metric was used. This algorithm ensures more intelligent centroid initialization, thereby improving the overall quality of the clustering. Beyond the initialization step, the rest of the algorithm remains consistent. To overcome the high dimensionality MSI data problem and to avoid data loss due to peak list generation, a pre-processing step involving data compression was introduced prior to segmentation.

For this purpose, several data reduction algorithms were investigated, including t-SNE, NNMF and SVD. As for PCA, t-SNE and NNMF are common preprocessing methods used for MSI data processing, compared to SVD. PCA and SVD are known to be suitable for linear dimensionality reduction and preserving global structure, NNMF is useful for non-negative data and part-based representation, while t-SNE excels in visualizing highdimensional data. As shown in **Fig. 1A**, SVD compression was found to be optimal. Indeed, even if t-SNE presented good segmentations for lipid (-) and peptide images, it was difficult to distinguish the GL from the ML and WM in lipid (+) and protein cases. The results using NNMF and SVD were correct, observing the three RB areas in each omics image (**Fig. 1A**). It can be added that the images generated by the latter have a better resolution and are more looking alike. Therefore, the SVD compression was kept for the future to obtain the best possible segmentations.

It is noteworthy that within the context of this investigation, only three out of the four primary cerebellum clusters were discernible using a 50 µm MSI spatial resolution: the ML, WM, and GL in conjunction with Purkinje cells. Additionally, lipid cerebellum images were captured at a finer resolution of 10 µm (**Fig. 1C**) and subsequently processed, thereby confirming the distinct visualization of all four cerebellum clusters. This underscores the crucial role that spatial resolution plays in the generation and differentiation of clusters, yet the spatial resolution was fixed at 50 µm since proteins imaging needs a higher resolution to get enough signals. Despite the potential for finer resolution to improve cellular component discrimination within the cerebellum, it was pragmatically determined that the 50 µm resolution sufficed for the objectives of this study, given the constraints and goals at hand.

#### Unsupervised cluster number estimation

We have shown that the three regions of the cerebellum can be observed in a similar way on lipid or protein image constructed with five clusters. However, the choice of the number of clusters was made in a semisupervised manner. To automate the process of lipid-based proteomics, it was necessary to implement a tool capable of estimating the optimal number of clusters. To estimate the correct number of clusters in a nonsubjective way, the silhouette criterion was used. The advantage is that it can be used multiple times, both to find the optimal number of clusters and to assess their stability and compactness.

As shown in **Fig. 1D and S4**, Silhouette estimated the optimal number of clusters at 5 for the lipid images, which was a coherent result with respect to the previously selected semi-supervised number. Furthermore, the fact that the same results were obtained for the lipids in negative or positive mode was expected due to their identical nature and metabolism. The 5 estimated clusters included 4 corresponding to the ML, GL, WM and brainstem regions of the rat brain, while 1 cluster represented a tissue-free area containing only matrix. These clusters were also observable for protein and peptide images with 5 clusters.

The same was true for predictions using protein and peptide data sets, which yielded a consistent and identical number of clusters. The heterogeneity predicted for peptides and proteins was more important, with a cluster number between 9 and 10. This can be explained by the diversity of proteins compared to lipids. Proteins are made up of a combination of 20 different amino acids, which may explain the presence of more protein clusters in the depth of the tissue compared to what is observed by lipid imaging or immunohistochemistry. Moreover, this over-segmentation in protein and peptide data may result from artefacts, particularly in the tissue-free regions. While we expect a single cluster to represent the matrix, as seen in the lipid data, we instead observed three distinct clusters, likely due to inhomogeneous crystallization (due to the nature of the matrix i.e. Norharman for lipids and HCCA-aniline vs SA-aniline for proteins). HCCA-aniline and SA-aniline are ionic matrices based on two component which explain the fact that we have 3 clusters instead to get only one (corresponding to HCCA, HCCA-aniline and aniline clusters or for SA, SA-aniline and aniline clusters). Taking account that in proteins and peptides due to the nature of the ionic matrix giving 3 additional clusters we can remove them and at the end we only have 7 clusters related to the tissue. Subdivisions were also observed in two clusters for molecular layer (possibly linked to the presence of Purkinje cells in some pixels) and brainstem, which were also found with lipids images with 10 clusters. Thus, in total we have 7 clusters for lipids and 7 clusters for peptides and proteins, as it can be seen in the **Figure 1D** for the 10 clusters images, still suggesting a degree of concordance between lipid and protein clustering in tissue regions.

For a simpler process, dry proteomics on lipid images was used because lipid imaging does not require additional sample preparation steps, protects the tissue from artifacts and potential degradation, and is less time consuming for routine analysis. Consequently, the clusters identified in lipid images are more representative of the RB cerebellum anatomy. In the present case, we relied on the principle of dry proteomics by lipid imaging and thus selected the 5 cluster omics images for the rest of the study.

Finally, the optimal segmentation workflow developed (**Fig. 1E**) was a MATLAB script, integrating a SVD compression of data with 10 principal components, combined with a *k*-means++ segmentation using a cosine distance with a silhouette criterion. This approach allowed the visualization of the three main clusters of the RB cerebellum (ML in blue, GL in orange, WM in light orange), in an identical and specific spatial localization, from the 5 cluster images respectively generated for: lipid (-) and (+) MSI with Norharman matrix (**Fig. S5 and S6**), lipid (+) MSI with DHB matrix (**Fig. S7**), protein MSI (**Fig. S8**) and peptide MSI (**Fig. S9**), with semi-supervised observation.

### Prediction model on lipid MALDI imaging

To automatically identify each cluster present in a tissue from a lipid image, a machine learning algorithm was trained on the 22 positive and negative lipid imaging datasets previously obtained. The ML, GL and WM centroids were extracted from the 5-cluster segmented lipid images and imported into Python. The datasets were subjected to peak picking and a non-parametric Kruskal-Wallis test to compare the significance of each ion between each ROI. Only features with a p-value equal to or less than 0.05 were retained as discriminant ions (**Fig. S10 and S11**). After isotope filtering, a final list of 36 lipid (-) and 19 lipid (+) discriminant ML, GL and WM ions were identified (**Fig. 2A-B**). The spatial distribution of each ion also confirms its specificity to its assigned cluster (**Fig. S12 and S13**). The annotation of the discriminant lipid ions was performed by SpiderMass MS/MS experiments, as its highest lipidomic similarities with the MALDI^38^ (**Fig. S14 and S15**). All the specific ions of the lipids in a region are listed in an internal database, which is used to predict regions on MALDI lipid images. Various prediction models were evaluated, respectively for each lipid mode analysis, taking in account the discriminant ions previously set out. In case of lipid (-) datasets, the SGD^42,43^ algorithm was selected as optimal model (**Fig. S16A**) and was validated using a 5-fold cross-validation^44^ with an accuracy of 94% (**Fig. S16B**). The Ridge classifier model was the one adapted to the lipid (+) datasets (**Fig. S17A**), with 98% accuracy after 5-fold cross-validation (**Fig. S17B**). The robustness of the developed models were subsequently evaluated by blind cohort validation, which included three different datasets of cerebellum RB lipid images for both polarity modes. Notably, in each instance, the model achieved 100% accuracy in its classifications (**Fig. S16C and S17C**).

**Fig. 2:**
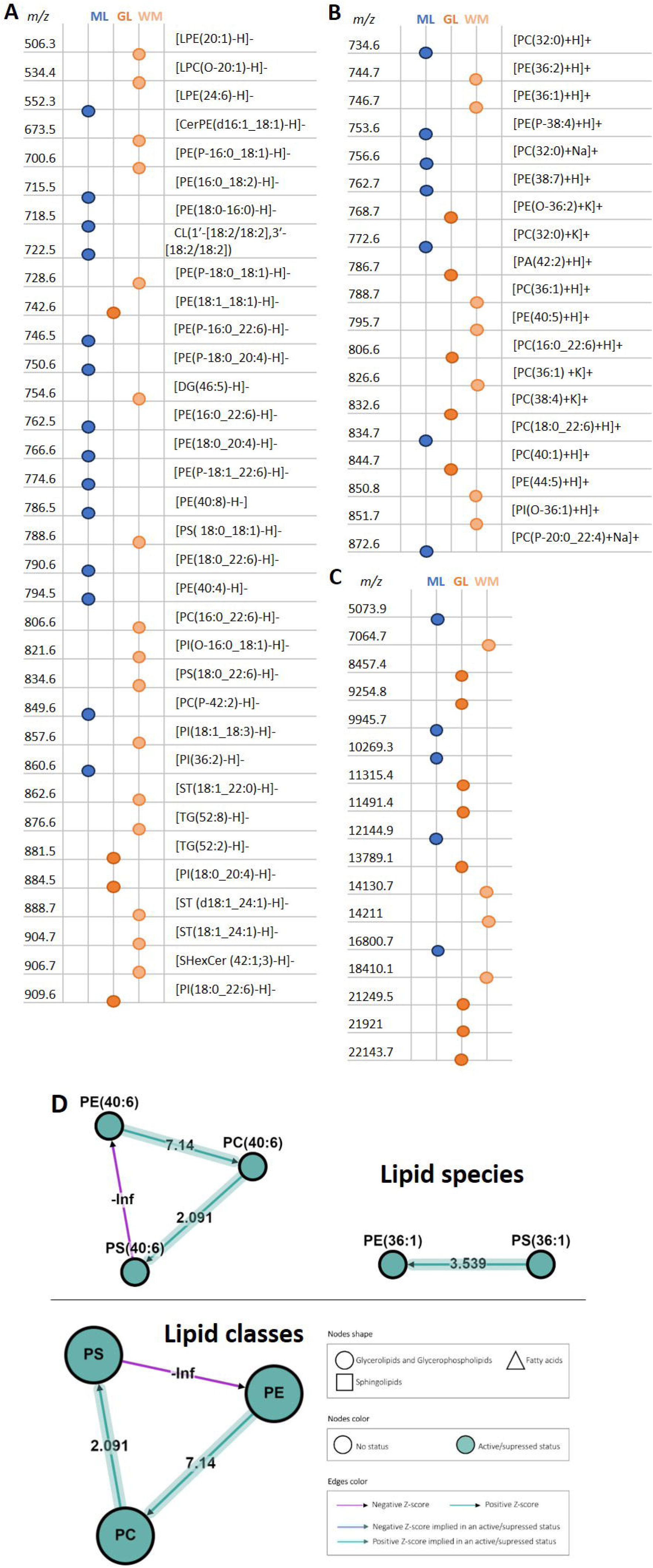
Discriminant lipid and protein ions present in RB cerebellum with BioPAN lipid pathways. Exhaustive list of **A)** 36 lipid (-), **B)** 19 lipid (+) and **C)** protein discriminant ML, GL, and WM cerebellum ions. **D)** BioPAN biological lipid pathways involved in white matter represented according to lipid species and lipid classes, with nodes legend.

The same data processing was performed on the protein imaging datasets to provide the corresponding discriminant protein ions for each RB cerebellum clusters (**Fig. 2C and S18**). The list of discriminant protein ions was then added to discriminant lipid ions for each cluster in order to create discriminant protein and lipid ions dataset reference for each RB cerebellum area.

### Lipids biological network analysis

Based on the compilation of annotated lipids, a greater number of lipids have been specifically identified in WM compared to GM. This observation is in line, considering that the WM is predominantly comprised of myelin, a substance containing a higher lipid content (78-81%) than both white (49-66%) and grey matter (36-40%)^45^. In the same way, myelin is composed of a high percentage of galactoceroboside and cholesterol compared to GM, which is why more diglycerides (DG), triglycerides (TG) and fatty compounds are identified in the latter. On the other hand, GM presents a higher percentage of phosphatidylethanolamines (PE) and phosphatidylcholines (PC), which correlate with presented annotations^45^.

To highlight the biological process involved by lipid data, a comparison between WM and GM discriminant lipid was performed on BioPAN^46^. The results, shown in **Fig. 2D** and **Fig. S19A**, revealed PC biosynthesis as the most active pathway in WM (with the involvement of PEMT predicted gene), whereas PE biosynthesis was observed as the most active pathway in GM (with the involvement of PISD predicted gene). These results clearly indicate discriminants lipids involved in specific biological pathways associated to distinct cerebellum regions.

Biologically, phosphatidylcholine is an essential choline reservoir for brain function^47^. In fact, choline is an important molecule for neurotransmission in neurons, which may explain the high activation of PC biosynthesis in WM. Phosphatidylethanolamine’s biological function is more due to its small chemical structure, which allows fluidity of the neuronal membrane^48^. The hypothesis is that this facilitates vesicle budding and membrane fusion^49^, a key step in synaptic transmission in GM. Finally, biological pathway based on lipids analysis showed that PC may be involved in the neurotransmission process in WM, whereas PE is more involved in synaptic transmission in GM^50^. These conclusions were further corroborated by Reactome analysis of the lipid dataset (**Fig. S19B**), which demonstrated their involvement in the neural system, signal transduction, small molecule transport or metabolism of protein and vesicle-mediated transport pathways.

### Consolidation method by protein pathway analysis

As discriminant biological pathways were defined for different regions of the cerebellum RB based on lipid species, the proteomes of these regions were analyzed to validate the hypothesis of a correlation between lipids and proteins within the same biological network. This analysis aimed to consolidate the dry proteomics processes.The ML, GL and WM regions observed in the multi-omics MALDI MSI were therefore subjected to spatial proteomics using the micro-proteomics workflow on three different RB sections^51,52^. By regrouping the triplicates for each cluster, a total of 5270 proteins were identified for WM, 5390 for GL and 5354 for ML (**Supplemental Spreadsheet S1**). This study confirmed the spatial heterogeneity of proteins previously observed in imaging. The results showed that discriminant lipid species for each ROIs are consistently linked to specific proteins in the same ROI, thereby forming region-specific pathways and functions.

Indeed, the Venn diagram, shown in **Fig. 3A**, considers the protein diversity between each region by the presence of proteins exclusive to each of them. In total, 85 proteins were exclusive, of which 7 were specific for WM, 11 for ML and 67 for GL (**Supplemental Spreadsheet S2**). It must be noted it was found among the 11 specific proteins in ML, two important enzymes involved in lipids metabolism *e.g.* Phosphoinositide phospholipase C and Inositol monophosphatase 1 whereas in GL, the Gamma-butyrobetaine dioxygenase know to catalyze the formation of L-carnitine and the Plasmolipin in WM, a main component of the myelin sheath involved in intracellular transport, lipid raft formation, and Notch signaling were identified^53^. The GL contain several neuropeptides or neuropeptide hormone activity such as Corticotropin-like intermediary peptide, Somatostatin-14; Pro-thyrotropin-releasing hormone, cholecystokinin-12; Neurokinin-B, Cocaine- and amphetamine-regulated transcript protein; Pituitary adenylate cyclase-activating polypeptide 27 or Ephexin-1^54^. In ML, among the identified protein the lamin B-binding protein (BAF: Barrier-to-autointegration factor) and Myogenin are of particular interest. In fact, BAF is required during brain development as a regulator of nuclear migration during neurogenesis of the CNS^55^. Myogenin is also detected in Allan brain atlas and is linked to motor neurons^56^. Similarly, in WM among the specific proteins identified, the Lymphocyte specific 1 is recently known to be correlated with tissue resident memory T cells^56^ and T cell infiltration^57^. Interestingly, Phosphatidylserine decarboxylase proenzyme (PISD) was also found in both WM and ML regions and was a predicted gene previously reported in BioPAN GM lipid pathway (**Fig. S19A**). The presence of PISD protein may explain the amount of PE identified in the ML region. In this context, PE may contribute to the integrity and function of neuronal membranes, influence synaptic transmission, and participate in signaling events. This again demonstrated the relevance of the different clusters by MSI, which predicted their own lipid/protein pathway and therefore biological heterogeneity.

**Fig. 3:**
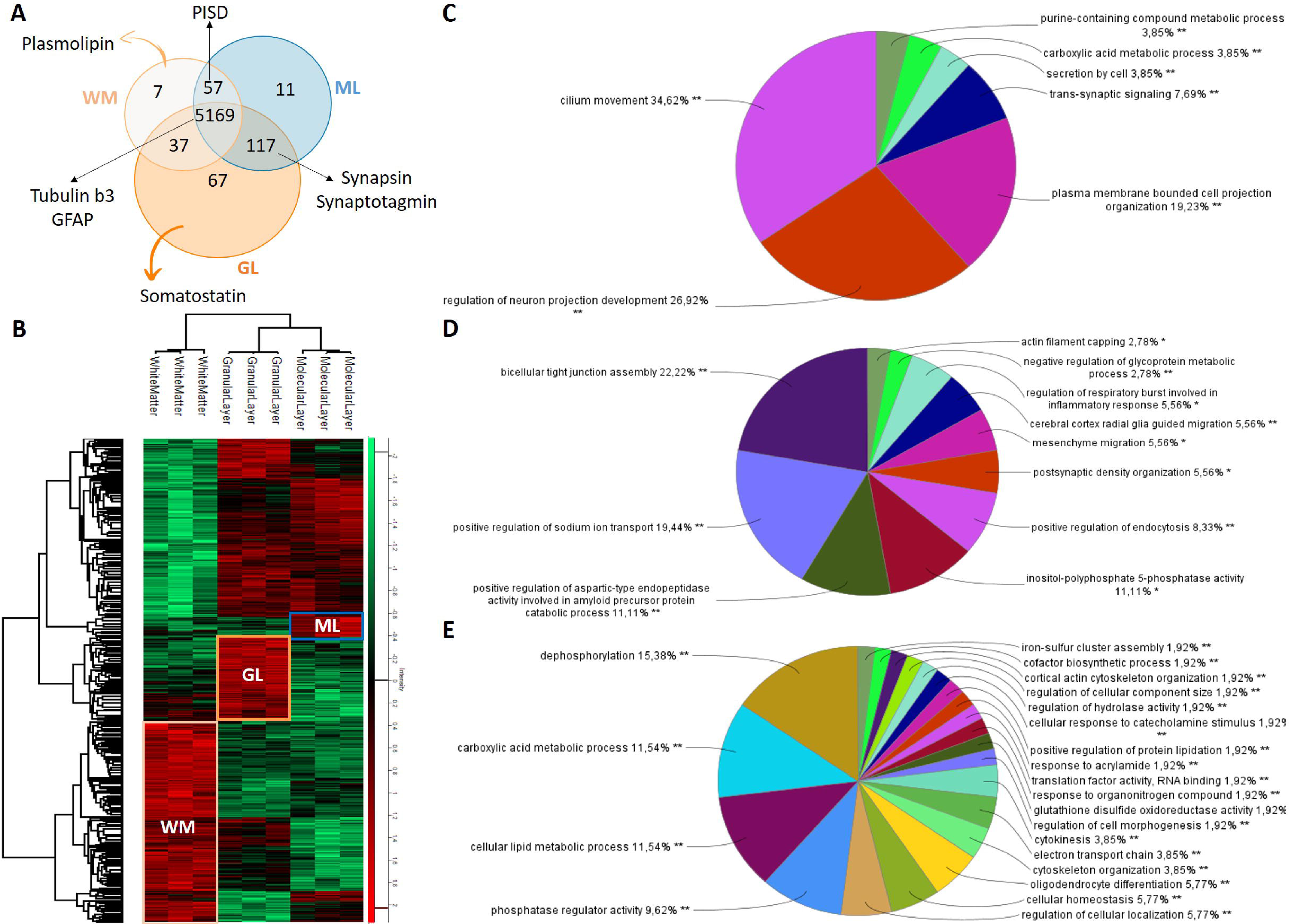
Rat brain cerebellum regions spatial proteomic analysis. **A)** Venn diagram of the specific proteins per layer. **B)** Heatmap after ANOVA (p-value <0.01) analysis demonstrated the presence of different of overexpressed proteins. ClueGO biological pathways involving the significant proteins found in **C)** granular layer**, D)** white matter, and **E)** molecular layer of the cerebellum.

To go further, the common proteins were subjected to an ANOVA test (p-value < 0.01) and showed that 2204 out of 5465 proteins have a significant variability of expression (**Supplemental Spreadsheet S3**). According to Allan brain Atlas, based on transcriptomic analyses, 196 genes are Cerebellum enriched gene and 59 out of those genes show highest expression levels in cerebellum. 90% of their corresponding proteins have been identified such as CBLN1 and CBLN3. Among them, some are known to be specifically located to the Purkinje layer which was regrouped with the GL after clustering. We were able to identify specific proteins from the Purkinje cells (MYH10, HOMER3, KIT, QKI, MX1, PCP-2, PP1R17, ARGEF33). For example, QKI protein expressed by radial astrocytes (Bergmann glia) with processes through the molecular layer all the way to the pial surface of the cerebellar cortex has been identified. MX1 is known to be in the dendritic processes of Purkinje cells. Moreover, other specific proteins of granular layer, GABRB2, TMEM6 and KCNIP4, markers of synaptic glomeruli from granular cells are also detected.

This was reflected by the presence of different clusters of over- or under-expressed proteins between each RB cerebellum area (**Fig. 3B**). The gene lists corresponding to over-expressed protein clusters were analyzed using ClueGO software to identify the biological pathways associated with the significant proteins identified in each distinct cluster. It turns out that the overexpressed proteins in the WM are mainly involved in myelination, glucose and neurofilament metabolism (**Fig. 3C and S20A**), which is a consistent result according to the bibliography^58^. In fact, WM consists of myelinated axons, so it’s involved in the transmission of nerve impulses by axons. The presence of glucose metabolism is also interesting when correlated with the galactoceroboside myelin composition previously suggested by lipid WM analysis. Furthermore, iron metabolism is another important biological process in the white matter, e.g. for myelin formation, redox reactions or neuronal development and synaptic plasticity^59–61^. This information can be linked to biological pathways previously found by lipids analysis, which also highlighted the neurotransmission pathway in WM. Regarding the GL, the neuropeptide hormone activity pathway was found to play a role in the processing and regulation of peptides that influence synaptic transmission, neural signaling and modulation of neuronal activity (**Fig. 3D and S20B**). Purine metabolism also plays a crucial role thereby influencing various physiological processes such as neurotransmission, synaptic plasticity, and energy metabolism. Dysregulation of purine metabolism in the brain has been implicated in several neurological disorders, including epilepsy, Parkinson’s disease, and neurodegenerative diseases. Similarly, the relevance of synaptic organization and sodium ion transport pathways involved in the molecular layer (**Fig. 3E and S20C**) were expected results given their role in neurotransmission and synaptic signaling between these cell types.^62^. It’s interesting to remember that the biological processes of synaptic transmission, vesicle transport and signaling were also predominant pathways in the previous lipid study. Thus, it has been shown that the ML, WM, and GL have their own specific proteome that can be correlated with specific lipid associated to distinct biological pathways.

### Dry proteomics based on RB horizontal lipid imaging application

#### Multi-omics RB horizontal sections generation

To validate the dry proteomics workflow to more complex tissue, the analysis was widened to total horizontal RB sections. As previously, multi-omics MALDI MSI were performed on 4 different sets of consecutive horizontal RB sections, and resulting data were submitted to the imaging data processing workflow, excluding matrix signal. The Silhouette criterion was around 11 for each lipid replicate, leading to multi-omics images composed of 11 clusters (**Fig. 4A and Fig. S21**). A similar spatial clustering shape was observed for each lipid image, including the well-known areas of the cerebellum, as well as other specific areas such as: the corpus callosum subdivided into clusters white, green and yellow, the cerebral cortex and thalamus in purple, red and pink, and the ventricular system in brown. These specific regions were also observed on the protein and peptide images built with 11 clusters, again confirming the lipid/protein pathway cluster appurtenance.

**Fig. 4:**
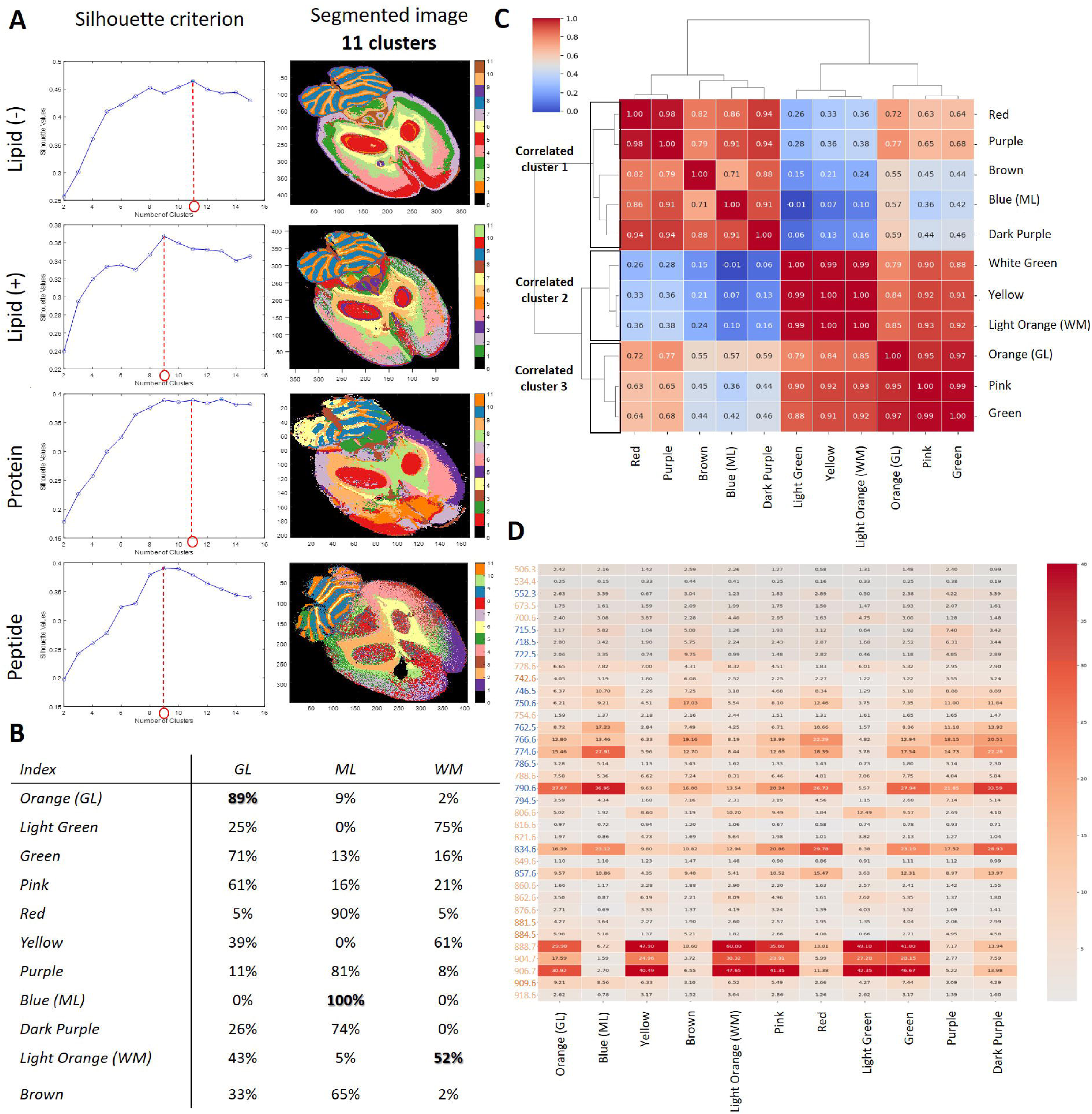
Horizontal rat brain section omics MALDI MSI analysis. **A)** Lipid (-), lipid (+), protein and peptide MSI segmentation images with 11 clusters and Silhouette criterion. **B)** Clusters mean scores prediction based on rat brain cerebellum lipid (-) model. **C)** Clusters Pearson’s correlation. **D)** Prediction lipid (-) model peaks involvement.

#### RB cerebellum lipid classification model: prediction on horizontal sections

Four replicate lipid (-) horizontal RB images were blindly analyzed using the pre-built classification model trained on 22 RB cerebellum lipid (-) MSI datasets (**Fig. S22**). The model returned a confidence score for predicting each ROI. Since the model was trained on three ROIs, the default confidence score to predict an ROI was >33%. The model successfully predicted the ML area with a mean confidence score of 100%, WM with a confidence score of 52%, and GL with 89% (**Fig. 4B**). For WM, although 52% is significantly higher than 33%, the lower confidence score may be due to the discrepancy in surface area between the sagittal and horizontal brain slices of the rats, with the former showing a significantly greater extent of WM. Other clusters were also analyzed using the predictive model (**Fig. 4B**) with interesting results. The light green and yellow clusters (corpus callosum region) were predicted as WM with confidence scores of 75% and 61%, respectively. Similarly, the green cluster (colliculus regions) was predicted as GL with a confidence score of 71%. A Pearson’s correlation of the discriminant lipid negative ions, shown in **Fig. 4C**, further validated these predictions. Two main clustering branches were identified: one leading to correlated cluster 1 associated with ML, and another leading to two separate clusters, correlated cluster 2 associated to GL and correlated cluster 3 associated to WM. In correlated cluster 1, dark purple and brown ROIs were grouped with ML, sharing the 774.6 and 790.6 lipid (-) ions (**Fig. 4D**). In correlated cluster 2, WM was grouped with the yellow and light green ROIs, as predicted by the model, with the main involvement of the 888.7 and 906.7 lipid (-) ions (**Fig. 4D**). Biologically, these results were expected. The corpus callosum (light green and yellow clusters) forms the largest commissural WM bundle in the brain, which has a distinct molecular composition due to its significant size and role, explaining the observed clustering^63^. Similar observations were valuable for the colliculus (green cluster), which also contains a superficial grey layer^63^. This explains the presence of orange color in both granular layer and colliculus clusters, corresponding to GM, and accounts for the 71% confidence score prediction explaining similarity^65^.

With the aim to justify the images segmentation, discriminant lipid (-) ions were identified for different cluster observed on the horizontal RB section lipid (-) image (**Fig. S23**). Many peaks were spatially distributed regrouping multiple clusters. For example, common ions were spatially distributed in ML, cerebral cortex and hypothalamus regions, like *m/z* 790.4, 834.4 and 886.5. The ion *m/z* 599.4 was collocated in GL and green cluster. Same observations for both the WM and the corpus collosum, for which ions as *m/z* 701.6, 889.6 and 904.7 were also spatially present. These ions could explain the correlation clusters highlighted by the Pearson’s correlation graph in **Fig. 4D**. However, discriminant ions were also extracted for specific regions, explaining their segmentation as single cluster during MSI data processing. The ventricular region (brown cluster) possessed various discriminant ions such as *m/z* 473.2 and 615.1. Specific ions were also discriminant for the red cluster, regrouping cerebral cortex and hypothalamus regions (*m/z* 746.6, 766.6, and 834.*5*), as well as for the corpus callosum like *m/z* 806.6. This ions list was added to the prediction model, to refine predictions.

#### Proteome horizontal RB section cluster comparison

To have a look at the proteome specificity of the RB horizontal section clusters, spatial proteomic analysis was also performed on the 7 clusters observed from the rat brain 11-cluster segmentation image (excluding the cerebellum cluster, already analyzed) (**Fig. 5A**). Proteins from red cluster were extracted from hypothalamus region, while proteins from purple and pink clusters were extracted from cerebral cortex. The green cluster was extracted from colliculus area, brown cluster from ventricular system and yellow and light green clusters from corpus callosum. Experiments were performed in biological triplicate. Data were processed with ML, WM and GL previous data in DIA-NN software for protein identification, quantification and correlation. By regrouping the triplicates for each cluster, more than 17243 proteins were identified, among them 5498 were proteotypics (**Fig. 5B**) (**Supplemental Spreadsheet S4**). Common proteins were subjected to an ANOVA test (p-value < 0.0001) and showed that 4481 out of 7223 proteins had a significant variability in expression. This was represented by the presence of different clusters of over- or under-expressed proteins between each extracted region (**Fig. 5C**). The resulting heatmap highlighted different clusters of overexpressed proteins (**Supplemental Spreadsheet S5**). First, cluster i consisted of proteins overexpressed in the cerebellum regions (ML, GL, and WM), while cluster v consisted of proteins overexpressed in the other regions. Specific overexpressed protein clusters were also highlighted for the ventricular system in cluster ii and for the corpus callosum in cluster iii. It was also observed that cluster iii was involved in WM, confirming their correlation in the previous lipid Pearson’s analysis (**Fig. 4C**). The overexpressed protein cluster iv was involved in the cerebral cortex and hypothalamus brain regions, explaining their similar image segmentation in the red cluster (**Fig. 5A**), Pearson’s correlation and prediction model using lipid (-) data (**Fig. 4C**).

**Fig. 5:**
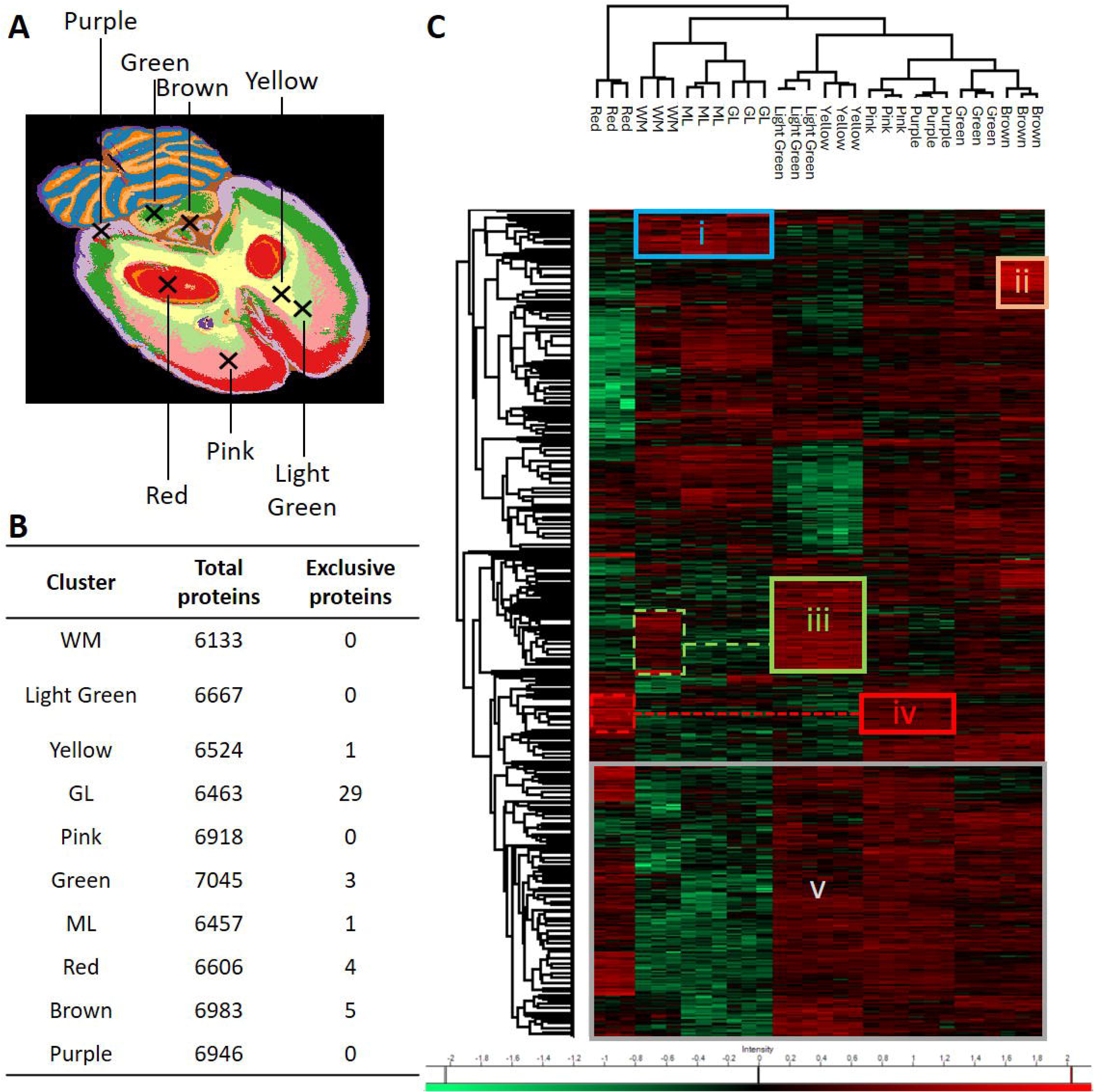
Spatial proteomic analysis of rat brain horizontal clusters. **A)** 10 different clusters identified thanks to lipid (-) lipid MSI and spatial proteomic extraction points. **B)** Protein Venn diagram. **C)** Heatmap after ANOVA (pvalue 0.0001) analysis demonstrated the presence of different of overexpressed proteins.

Biological pathway analysis of these later over-expressed protein clusters also confirms this observation (**Fig. S24**). Indeed, biological pathways involved in cerebellum (cluster i) were mainly concentered around synapse metabolism, with myelination, paranodal metabolism, neurofilament assembly, and calcium/sodium transport (**Fig. S24A**). We could notice that this biological process also resumed the one’s independently found for ML, GL and WM (**Fig. 3**). At the opposite, biological process involved in the cerebral cortex (cluster v) contributed to at least NMDA selective glutamate receptor signaling, regulation of neurotransmitter receptor transport (endosome to postsynaptic membrane) (**Fig. S24B**). This distinction of biological process well defined and distinguished the cerebellum and cerebral cortex regions of the brain. Indeed, the cerebellum is primarily involved in coordinating motor movements, maintaining posture and balance, and motor learning^64^, whereas the cerebral cortex is responsible for higher cognitive functions including perception, memory, attention, language, and consciousness^65^. Same conclusions were observable analyzing biological pathways specifically involved in cerebral cortex and hypothalamus, in cluster iv, where mains pathways regrouped vocal and auditory learning, memory and feeling process with serotonin metabolism (**Fig. S24D**). Likewise, myelination and neurofilament pathways were involved in cluster iii, for corpus callosum RB area (**Fig. S24E**), which was linked to WM biological pathways. The biological pathways for cluster ii, specific of ventricular system RB region, was also analyzed. It turned out that cholesterol, triglyceride, and blood coagulation regulation were the most relevant pathways (**Fig. S24C**). These results fit with the neuroanatomy of ventricular system, where cerebrospinal fluid flows in the regions thanks to blood pulsations in surrounding blood vessels^66^. Furthermore, triglycerides cross the blood-brain barrier and are found in cerebrospinal fluid helping in satiety and cognition mechanisms^67^.

In this way, we were able to show from a protein pathway point of view that cerebellum regions are distinct from the cerebral cortex regions, which itself consists of several specific areas. Their proteomes were also integrated into the model with their paired lipid clusters. In addition, proteomic data of this study were in line with previous analysis already performed on RB regions from published studies. This allowed to add more information to the RB dry proteomics model. First, we compared proteins identified here in bottom-up, with proteins identified by top-down in the hippocampus and corpus callosum RB areas, presented in a previous study^3^. According to Delcourt, V. *et al.,* 2018^3^, 16 over 22 proteins identified in top-down for the corpus callosum were also identified and over-expressed in this area according to presented protein dataset. Same observations for 15 proteins over the 20 identified in top-down for the hippocampus (**Supplemental Spreadsheet S6**).

#### 52Workflow robustness

The robustness of the dry proteomics workflow was thoroughly assessed by examining the redundancy of spectral lipidome and proteome identifications within each cluster across independent triplicates. To ascertain clustering repeatability, the spectral lipid (-) dataset from each cluster was compared among triplicates, as illustrated in **Fig. S25A**. Impressively, an average of 99% of common lipid (-) ions was consistently identified across replicates within clusters (refer to **Fig. S25C**). Similarly, an in-depth analysis of the spatial proteomic dataset, with a specific focus on distinct clusters, revealed a remarkable consistency, with 93% of the proteins consistently identified across each replicate extraction point within a cluster. This robustness is highlighted in **Fig. S25C**, which succinctly summarizes the percentage of common protein identifications in replicates for each cluster (**Fig. S25B**). Notably, it’s worth mentioning that proteins involved in cluster-specific pathways, as previously depicted in **Fig. 4A** were fully recovered at a 100% rate in subsequent analyses. This underscores the reliability and reproducibility of the methodology employed in capturing proteomic signatures associated with distinct cellular clusters. This reproducibility is the essence of dry proteomics. For future analyses, there’s no need to redo spatially resolved proteomics. Simply start with a lipid image and query the dry proteomics model to reliably determine the cluster type, associated proteins, and relevant biological pathways.

### Glioblastoma tumoral heterogeneity analysis

#### Lipid and peptide MSI segmentation correlation

Finally, we performed the dry proteomics workflow on a prospective and retrospective cohort of glioblastoma (GBM), re-using collected data from Duhamel, M. *et al*., 2022 study^27,28^. The previous study performed patient’s stratification based on spatial proteomic and spatial lipidomic guided by MALDI MSI associated to patient survival^27,28^. The cohort consisted of 50 GBM patient tissues, referenced to P1 to P53 (**Fig. S26 and S27**). Peptide MALDI MSI was performed for all samples, and lipid MSI was conducted for 13 of these tissues. Thus, peptide and paired lipid images were collected for these 13 patients and were processed through developed data imaging workflow. Initially, each tissue was analyzed individually to assess its heterogeneity using Silhouette criterion and generate segmented images. Subsequently, peptide and lipid images were created with 8 to 13 clusters each. The findings of this study revealed an intriguing correlation between lipid and peptide distributions in samples labeled P1 to P14, as evidenced by the generation of highly similar numbers of clusters in both types of images. This correlation underscores the inherent link between the spatial heterogeneity of peptides and lipids within the tissue microenvironment (refer to **Fig. 6A and Fig. S26**). Furthermore, segmentation analysis effectively mirrored histological annotations, enabling the delineation of distinct regions of tumoral proliferation from necrotic or inflammatory areas (as depicted in **Fig. 6A**), as it was also evocated by Duhamel, M. *et al*., 2022 in previous studies^27,28^. Prior investigations have primarily relied on lipid and protein to differentiate between these three main tissue types based on specific molecular signatures. In contrast, dry proteomics segmentation workflow offers a more detailed representation of the intricate composition of biological tissues. This enhanced segmentation, not only facilitates the precise identification of pathological features but also reveals previously undetected levels of heterogeneity within tumor, necrotic, and inflammatory regions. This not only achieved improved delineation between annotated areas but also unveiled a greater-than-expected level of heterogeneity within these regions. This heightened resolution enhances understanding of tissue composition and offers valuable insights into the underlying biological processes driving tumor progression and response to treatment (**Fig. 6A**). For instance, in the case of P12, a nuanced examination of the proteomic extraction points unveiled intriguing insights. Points 12.1, 12.3, and 12.4, which were annotated as tumoral in the histopathological scan, exhibited a complex molecular landscape. Notably, point 12.2 was identified as bearing both tumor and inflammation characteristics. However, upon closer inspection using lipid and peptide MSI, it became evident that point 12.4 shared similar molecular profiles with point 12.2, in stark contrast to points 12.1 and 12.3. This striking observation was further corroborated by protein extraction analysis, which revealed distinct correlations among the points. Specifically, points 12.1 and 12.3 exhibited a notable correlation, indicating shared molecular features, while points 12.2 and 12.4 formed a separate correlated cluster (**Fig. 6D**). This delineation underscored the intricate heterogeneity within the tumor microenvironment, where discrete molecular signatures delineated different regions, potentially indicative of diverse biological processes or cellular compositions.

**Fig. 6:**
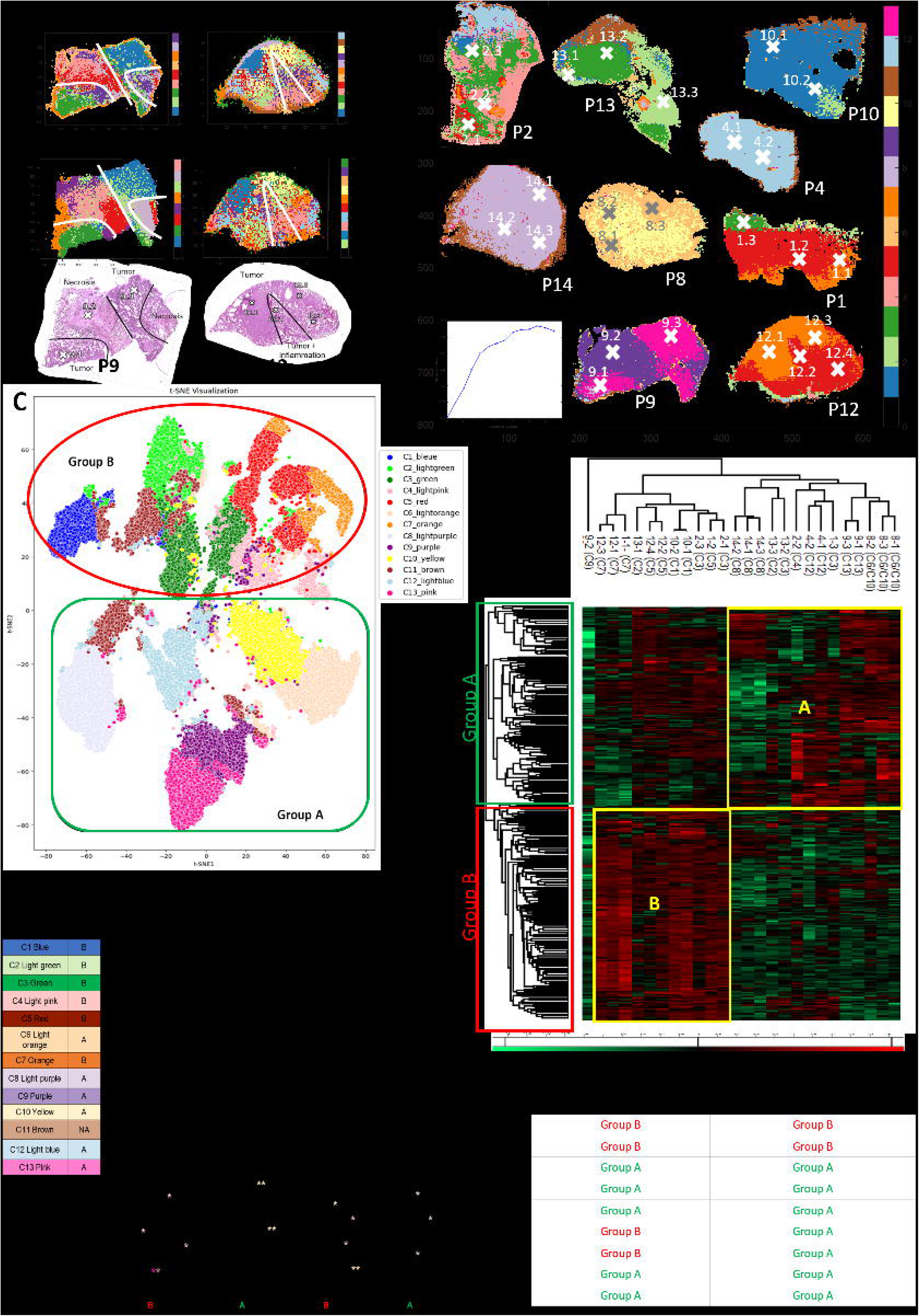
Glioblastoma patient lipid and protein heterogeneity analysis. **A)** P9 and P12 lipid and peptide MSI with histopathological annotations. **B)** Co-segmentation of 9 tissues previously analyzed by lipid MALDI MSI. **C)** t-SNE representation of each cluster identified through lipid co-segmentation. **D)** Protein heat map after ANOVA (pvalue 0.01) analysis demonstrating the presence of different of over-expressed proteins according lipid clusters. **E)** Patient classification group A and B prediction according lipid and protein model. **F)** Lipid cluster and associated protein blind prediction on patient P3, P5, P6 and P11.

#### Lipid-MSI clusters classification and proteomic correlation

To have a large view on the general heterogeneity on the whole cohort, a co-segmentation was performed on 9 lipid images dataset. It turned out that 13 different clusters were shared between these 9 patients’ tissues (**Fig. 6B**). Some clusters were correlated to biological specific tissues regions according to histopathological annotations. In this way, clusters 4 (light pink) and 9 (dark purple) were identified as necrosis tissues, clusters 1 (blue), 2 (light green) and 7 (orange) seemed to be specific tumors, whereas clusters 3 (green) and 5 (red) were tumoral areas near to inflammation and clusters 6 (light orange), 8 (light purple), 10 (yellow), 12 (light blue) and 13 (pink) were tumoral areas with necrosis. Clusters were predominantly identified within specific tissues, such as cluster 9 primarily present in P9, or shared across multiple tissues, as observed with cluster 3 in P1, P2, and P13. Once more, the segmentation underscored the molecular diversity within necrotic and tumoral regions, revealing a mosaic of numerous clusters.

A t-SNE representation of tissue lipid imaging clusters allowed to distinguish two mains groups of clusters based on lipid MSI (**Fig. 6C**): group A was regrouping clusters 6, 8, 9, 10, 12 and 13, while group B regrouped clusters 1, 2, 3, 4, 5, and 7. Cluster 11 was shared between the two groups. A correlation heatmap, presented in **Fig. 6D**, also highlighted the correlation between lipid clusters regrouped in group A and B.

The proteomic data obtained from nine distinct tissue samples were leveraged to conduct a comparative analysis of the various clusters identified through lipid imaging segmentation. Notably, specific extraction points analyzed in this study correlated with clusters identified in lipid imaging (**Fig. 6B**). Through statistical analysis, employing an ANOVA test with a significance threshold set at p < 0.01, we identified 373 out of 3616 proteins exhibiting significant variability in expression levels (**Supplemental Spreadsheet S7**). First, biological pathways were identified for each cluster through ClueGo analysis (**Fig S28**), based on the overexpressed proteins present in each. Interestingly, some pathways were specific to particular lipid clusters. For example, the RAC3 GTPase cycle pathway was unique to cluster 7 (**Fig S28F**), playing an important role in neuronal development and tumor progression^68^. L1CAM expression was particularly found in cluster 1 (**Fig S28A**), underscoring the tumor aggressiveness of this cluster. This pathway is a focal point of active investigation in GBM due to its profound implications for tumor aggressiveness, invasion, therapeutic resistance, and poor prognosis. Similarly, overexpressed proteins in cluster 5 were specifically involved in the axon guidance pathway^69^, which is currently a therapeutic area of research for the treatment of malignancy. On the other hand, some biological pathways were common across multiple clusters. Notably, the interleukin-12 family signaling pathway^70^, a current therapeutic target in cancer immunotherapy, was identified in clusters 6, 9, 10, and 13 (**Fig S28E, H and J).** Similarly, the ECM proteoglycans pathway^71^ associated with tumor development in GBM was found in clusters 4 and 9 (**Fig S28C and H**). Finally, the biological pathway analysis of each cluster revealed distinct characteristics: some clusters exhibited a more aggressive GBM pattern, whereas others showed a less aggressive pattern and identified potential therapeutic targets.

Further investigation on protein data allowed to compare the proteome of each cluster and identify correlations between them. It revealed the presence of 2 distinct clusters of over-expressed proteins, namely protein cluster A and B (**Fig. 6D**). Of particular interest, protein cluster A was found to correspond to regions of necrotic tissue, encompassing the imaging clusters 9 and 4 previously described. To gain deeper insights into the biological processes associated with these necrotic regions, we performed ClueGO analysis on protein cluster A, utilizing GOterms and Reactome databases (**Fig. S29A**). This analysis unveiled a multitude of signaling pathways implicated in necrosis processes. Notably, pathways such as platelet degranulation, blood coagulation, MyD88 deficiency, and IRE1 chaperone activation emerged as significant contributors in modulating cell death processes, including necrosis and can influence tissue damage and disease progression in various pathological conditions such as GBM. In the same way, an intriguing correlation in protein cluster A was observed among protein extracted from lipid imaging clusters 6, 8, 10, 12, and 13, as depicted in **Fig. 6D**. The later result confirmed the lipid image cluster classification in group A, proposed previously according lipid MSI co-segmentation analysis (**Fig. 6C**). This cluster notably encompassed tumoral clones characterized by the presence of necrotic regions. Through ClueGO analysis, the significant implication of selenoamino acid metabolism within this cluster was unveiled, shedding light on its pivotal role in the pathogenesis of glioblastoma. This pathway was also individually identified previously in **Fig. S28E and J** in lipid cluster 6, 10 and 13. Selenoamino acids, such as selenocysteine and selenomethionine, are fundamental constituents of selenoproteins, where selenium, an essential trace element, is incorporated. These selenoproteins orchestrate a myriad of cellular processes, including antioxidant defense, redox regulation, and DNA synthesis and repair. The dysregulation of selenoamino acid metabolism has been implicated in the intricate progression of GBM through various mechanisms, contributing to disease aggressiveness and resistance to therapy. Similarly, the over-expressed proteins identified within protein cluster B, primarily comprising lipid imaging clusters 3, 4, 5, and 7, yielded significant insights, particularly regarding the involvement of L1CAM interactions (**Fig S29B**) from cluster 1 **(Fig. S28A).** Protein cluster B suggested a more aggressive tumor phenotype compared to those within protein cluster a, with implications for poor prognosis or short survival prediction. The intricate interplay between selenoamino acid metabolism and L1CAM interactions underscored the multifaceted nature of GBM pathogenesis, highlighting potential avenues for targeted therapeutic interventions and personalized treatment strategies aimed at mitigating tumor progression and improving patient outcomes.

Finally, two distinct classification groups, labeled group A and group B, were highlighted and cross-validated between lipid MSI and proteomic analysis. Proteins from the over-express protein cluster A were associated to lipid cluster A, resulting in group A. Thus, group A was associated to the lipid clusters 6, 8, 9, 10, 12, 13, and protein, involving specific protein pathways with a pivotal role in GBM, such as selenoamino acid metabolism. In another hand, group B regrouped lipid clusters 1, 2, 3, 4, 5, 7, and the over-expressed protein cluster B, which possessed more aggressive protein pathways with the implication L1CAM interactions.

#### Patient proteome blind prediction based on lipid cluster classification

To predict patient proteome through group A and B, two distinct classifications models were developed. Firstly, a model was trained on the lipid-MSI data from the 13 clusters comprising groups A and B. The aim of this classification model was to classify patient tissue according to lipid images, and associate their paired protein pathway. The resulting model was built with LGBM algorithm with an accuracy of 97% after 5-fold cross validation with an individual accuracy up to 95% for each cluster (**Table 1**, **Fig. S30B-C**). Specific lipid ions involved in the model were extracted and identified in specifics clusters using LIME algorithm. The top lipid biomarkers implicated to classify each cluster with 82.3% of contribution were summarized in **Fig. S30E and S31**. For example, lipid with *m/z* 770.35 was specific to group A, with a significative presence in lipid cluster 8 with highest contribution weight at 70% (**Fig. S30C**). Likewise, the *m/z* 798.64 was specially distributed in lipid MSI cluster 5 (**Fig. S32**) with a contribution weight at 61% (**Fig. S30C**), associating it with group B specific marker.

Thus, the classification of all 9 patients was carefully reviewed according to the patient group A or B classification model, based on the presence of specific lipid clusters in tumoral tissue. In scenarios where tissue samples exhibited clusters overlapping both group A and B, they were unequivocally classified into group B, prioritizing the presence of markers indicative of unfavorable outcomes. This approach ensured a rigorous and systematic evaluation, wherein each case was subjected to thorough examination, with particular emphasis placed on identifying and prioritizing markers associated with poorer prognostic indicators. By adhering to the following patient classification method, the prognostic assessment process maintained an exemplary level of precision and consistency, empowering clinicians to render well-informed decisions regarding patient management and treatment strategies. As depicted in **Fig. 6E**, 4 patients were classified in group B and 5 patients in group A. It’s noteworthy that previous investigations have emphasized the importance of assessing patient classification based on the expression levels of key proteins^28^. In 9 patient’s cohort (outlined in **Fig. 6E**), prior studies classified 2 patients in group B and 7 patients in group A using this protein panel^28^. However, resulting analysis unveiled a nuanced disparity in group classification for patients P10 and P12. This discrepancy can be attributed to the incorporation of molecular heterogeneity into analysis, offering additional insights into survival prediction. Furthermore, upon scrutinizing the co-segmentation analysis illustrated in **Fig. 6B**, it became evident that P12 and P1 shared significant cluster composition. Given P1’s association with group B, it was reasonable to surmise that P12, sharing similar cluster characteristics, would also be classified within group B. This observation underscores the importance of integrating molecular heterogeneity and comprehensive data analysis techniques to refine classification assessments and enhance clinical decision-making processes.

The 4 last patient tissues, for which lipid-MSI and protein data were available (P3, P5, P6 and P11), were blindly interrogated in classification model based on lipid-MSI clusters. P3 and P11 presented the IDH1 mutation and were not considered in the studies of ^27,28^. Upon blind interrogation of the lipid cluster images, patients 3 and 6 harbored a non-negligible percentage of lipid clusters 4 and 2, leading to the prediction to proteins associated to wound healing, or ECM proteoglycans biological pathways for example (**Fig. 6F**). The presence of the latter lipid cluster and biological pathways in patient 3 and 6 were thus indicative of the group B classification. Conversely, patients 5 and 11 mainly predicted with high percentage of lipid cluster 6 and 8, allowed the prediction of proteins associated to biological pathways such as Interleukin-12 family signaling, peptide chain elongation or RHO GTPase active ROCKS (**Fig. 6F**). In this way, patient 5 and 11 were classified in group A. This result also correlated with a lipid MSI co-segmentation performed on the 13 tissues (**Fig. S33**). The resulting image was composed of 14 clusters according Silhouette criterion. Interestingly, P6 and P3 were segmented apart of the rest of the cohort, suggesting a possible new lipid class. P5 and P11 tissue associated to group A were sharing specific clusters with P8 and P14, already previously classified in group A.

Complementary, a second classification model was constructed using RidgeClassifier with group A and B protein data (**Fig. S34**). The objective was first to intricately cross-validate the lipid MSI-based classification model. The resulting model had an accuracy of 96%, with a 5-fold cross validation (**Table 1** and **Fig. S34F**). Specific proteins involved in the model decision-making were identified in specifics clusters (**Fig. S35**). Among them, group A and group B biomarkers were distinct, referring to selenoamino acid metabolism or L1CAM interaction pathway for instance. This sophisticated approach underscored the synergy between lipidomic and proteomic analyses in refining group A and B classification for glioblastoma patients, thus paving the way for personalized therapeutic interventions tailored to individual risk profiles.

Thus, previous finding was further reinforced by the protein classification model, which concurred in its classification assessment, designating patients P3 and P6 to group B, whereas P11 and P5 were classified to group A. Hence, both the lipid-MSI clusters and protein models converged in classifying these patients within group A, or B. This alignment serves to authenticate the reliability and validity of the classification model, as well as enhancing the dry proteomics concept on clinical study as GBM^28^

#### Groups classification and patient outcome correlation

The dry proteomics developed pipeline, in both using lipid-MSI data and proteins classification models, led to the discernment of two distinct classes in GBM study, labeled as classification group A and B, illustrated in **Fig. 6-D**. Leveraging patient survival data, prognostic outcomes were correlated with specific lipid-MSI clusters. The clinical characteristics of the patient, evocated in studies^27,28^, revealed that patients involved in group A, through lipid-MSI clusters 6, 8, 9, 10, 12, and 13 implication, were upper the survival interquartile range with a survival outcome surpassing 32 months. In the same logic, patients associated to group B, with the presence of lipid clusters 1, 2, 3, 4, 5, and 7, had a poorer survival prognosis of less than 30 months.

Indeed, some of lipid biomarkers involved in lipid-MSI classification model were already recognized as prognostic markers in previous research^27^. For instance, lipid ions with *m/z* of 864.7, 866.7, and 881.7 were identified in both studies as markers for survival outcomes exceeding 36 months, primarily present in clusters 8 and 9 from group A. Conversely, lipid ions such as *m/z* 760.6, 788.6, and 810.6 were associated with shorter survival durations, less than 30 months, and were distinctly present in clusters 2 and 5 from group B. Moreover, these significant findings were consistent with prior investigations, reinforcing the notion that protein group B typically correlates with a poorer prognosis compared to group A. Particularly notable was the identification of over-expressed proteins ANXA6 and GPHN within group B, both previously implicated as unfavorable prognostic indicators ^28^. Conversely, group A exhibited elevated expressions of proteins RPS14 and MTDH, associated with more favorable prognostic outcomes^28^. Thus, the identification of group A and B lipid features by MALDI MSI, would automatically provide the paired protein pathways (**Supplemental Spreadsheet S8**), associated to short or long survival patient outcome.

As the left 37 patients were only analyzed through peptide MALDI MSI and spatial proteomics due to the data reuse, the later were interrogated through classification model with proteomics data, to predict their appurtenance to group A and group B, and thus their protein networks and prognosis (**Fig. S36**). Finally, among the cohort of 50 patients, 11 patients were classified in group A with a prognosis survival outcome >32 months, whereas 39 patients were classified in group B with a survival outcome <30 months. The latter results correlated with the clinical characteristics of the patient evocated in study ^28^. Indeed, 4 patients with IDH mutation were excluded, 12 patients were upper the survival interquartile range (IQR) set at 13.5 and 32 months, 23 patients were intermediate IQR, and 11 patients were lower IQR.

### Dry proteomics limitations

Although the dry proteomics model is robust, fast, and simplifies the analysis of complex heterogeneous tissues, it has some experimental and predictive limitations.

Technically, it is impossible to obtain identical consecutive tissue sections due to the z-dimensional factor related to tissue depth during cryostat sectioning. For example, we observed less structural changes between consecutive sections of the cerebellum. However, in horizontal sections, where the anatomy is more complex and variable, differences between consecutive sections are noticeable. These differences affect the imaging of lipids, proteins and peptides due to anatomical changes with depth. To address this issue, we performed spatially resolved proteomics on the same section used for lipid imaging. Once the model is trained, dry proteomics becomes a useful tool because only one lipid image is needed to assess the heterogeneity, identify the clusters, and associate the proteome, avoiding issues related to anatomical changes in consecutive sections. The second limitation concerns the predictive ability of the model, which is based on experimental data of clusters obtained by segmentation of lipid images. A reliable and accurate model requires a large cohort with representative replicates of the studied population. Building a generalizable model is challenging because some tissue-specific clusters may not be represented in our analyses. When the model encounters an unknown cluster that it hasn’t been trained on, it will likely misclassify it by approximating a known cluster. There are two ways to address this problem. First, by checking the approximation of an unknown cluster by the unsupervised k-means++ and t-SNE models. This involves plotting the matrix of this cluster on the k-means++ and t-SNE axes to see which known cluster it is close to, thereby confirming or disproving the model’s predicted approximation. Second, consider the use of self-training algorithms in the future ^72^. This involves retraining our model with known labeled clusters and new unknown and unlabeled clusters to improve and update the model specifically for clinical routine use. In this case, it will also be necessary to update the proteomic data for the new unknown clusters.

To extrapolate the strategy of dry proteomic to other tissue types or diseases, different learning model approaches are possible. The first one consists in a specific model for a specific tissue type or disease. In this case, the model would be trained on clusters specific to a particular tissue type or disease. While this approach is limited to the heterogeneity of that single tissue or disease, it offers greater accuracy by focusing on fewer clusters, which reduces the risk of false positive predictions (fewer classes in a multi-class classification task). This results in a more targeted and precise model. The second possibility is to improve the model in an agnostic model. This is a global model designed to work across multiple tissue types or diseases. To improve its performance, the model would need to be trained on clusters from various diseases and tissue types. Such a model would be capable of predicting and identifying clusters specific to particular tissues or diseases, while also recognizing common clusters across different tissue types. This approach could be especially useful for large-scale studies, such as PAN-cancer research. However, agnostic models are typically less accurate and require sophisticated feature engineering to enhance their performance. Another strategy to improve agnostic models is to use a transfer learning approach, where specific models are trained on individual diseases and then adapted for broader applications. Once refined, this type of agnostic model could also be applied to study metastasis and help trace the origin of cancers.

## DISCUSSION

We presented an automated dry proteomics approach based on lipid MALDI-MSI, addressing several challenges to establish cluster-specific lipid and protein correlations in terms of imaging and pathways. Optimizations in the segmentation pipeline using SVD data compression, k-means++ segmentation and the Silhouette criterion enabled the correlation of multi-omics MALDI-MSI data. The integration of the Silhouette criterion proved useful for determining tissue heterogeneity, identifying the most optimized number of clusters in a fully automated and unsupervised manner.

Using the RB cerebellum tissue model, we demonstrated the workflow’s suitability for lipid, protein, and peptide imaging, outperforming other segmentation algorithms. The robustness of our MS image processing model was confirmed through numerous experimental replicates. Multi-omics segmented images revealed the presence of RB cerebellum clusters ML, GL, and WM, each with specific spatial localizations, distinct lipid and protein compositions, and associated biological pathways. A predictive model was developed based on these specific lipid fingerprints, complemented by the unique protein compositions and paired biological pathways of each cluster. Pathway analysis validated the dry proteomics approach for GL, ML, and WM.

To extend the prediction model beyond cerebellum regions to more complex tissues, RB horizontal slices were analyzed using multi-omics MALDI MSI. This analysis successfully identified several clusters with unique spatial localizations, including cerebellum clusters. The model, trained with cerebellum lipid datasets, effectively annotated these areas and provided insights into their specific lipids, proteins, and associated biological pathways. Further analysis refined the model for more accurate predictions, improving our understanding of complex tissue composition and highlighting the potential of dry proteomics to elucidate intricate biological processes.

Applying the dry proteomics workflow to glioblastoma patient cohorts provided profound insights into the spatial heterogeneity of peptides and lipids within the tumor microenvironment. By combining previous research data with cutting-edge imaging techniques, we uncovered previously unexplored complexity within GBM tissues. The segmentation process accurately delineated pathological features and revealed nuanced variations within tumor, necrotic, and inflammatory regions, providing a detailed representation of tissue composition.

The observed correlation between lipid and peptide distributions underscores their potential as robust biomarkers for tumor characterization. Our analysis revealed distinct molecular signatures within different tumor regions, indicating distinct biological processes and cellular compositions. Co-segmentation identified 13 discrete clusters among patients that corresponded to specific biological tissue regions. Proteomic data integration enriched our understanding of the molecular landscape within these clusters. Statistical analyses revealed significant protein expression variability across clusters, identifying distinct biological pathways. Indeed, the ClueGO analysis highlighted the involvement of different pathways, such as selenoamino acid metabolism or L1CAM interactions, which have a remarkable impact on GBM pathogenesis.

By integrating dry proteomics with prognosis, this study culminated in the development of a sophisticated classification model for GBM patients to identify the type of clusters and corresponding proteomic data with region-specific pathways and functions to stratify different prognostic categories. Two resulting GBM patient groups, A and B, were predicted using a model based on GBM lipid MSI that incorporated molecular heterogeneity within tumor tissues. The model was also translated into a proteomic model capable of distinguishing groups A and B based on protein data. It validated the model based on lipid MSI data and classified patients using only proteomic data. The protein networks of group A correlated with survival of more than 32 months, while those of group B correlated with survival of less than 30 months. By classifying patients’ tumors into groups A or B, we were able to predict tumor protein networks that correlated with classification group membership and survival prognosis.

In essence, this project highlights the importance of integrating multi-omics approaches for comprehensive prognostic assessment in GBM. By unraveling the interplay between molecular features and clinical outcomes, the developed model provides invaluable insights to inform personalized treatment strategies and improve patient management in the complex GBM landscape.

Finally, the concept of dry proteomics, which first identifies tissue heterogeneity and distinct clusters by lipid imaging and then automatically associates specific proteins and biological pathways involved in each cluster, has proven essential for clinical applications. These insights facilitate the identification of potential therapeutic targets or prognostic markers, as demonstrated in the glioblastoma study, paving the way for improved patient outcomes and personalized treatment strategies.

## Supporting information

Graphical Abstract

## ABBREVIATIONS

CID: Collision Induced Dissociation
DG: Diglyceride
FF: Fresh Frozen
GBM: Glioblastoma
GL: Granular Layer
GM: Grey Matter
GO: Gene Ontology
LIME: Local Interpretable Model-agnostic Explanations
LGBM: Light Gradient Boosting Machine
MALDI: Matrix Assisted Laser Desorption/Ionization
ML: Molecular Layer
MSI: Mass Spectrometry Imaging
NNMF: Non-Negative Matrix Factorization
PC: Phosphatidylcholine
PCA: Principal Component Analysis
PE: Phosphatidylethanolamine
RB: Rat Brain
RMS: Root Mean Square
ROIs: Regions of Interest
SGD: Stochastic Gradient Descent
SVD: Singular Value Decomposition
TG: Triglyceride
t-SNE: t-distributed Stochastic Neighbor Embedding
WM: White Matter

## DATA AVAILABILITY

The data from this study, including MS DIA raw files, DIA-NN files, and annotated MS/MS datasets, have been deposited to the ProteomeXchange Consortium via the PRIDE partner repository with the dataset identifier PXD054488 (**Username:** - **Password:** vg74fZoVimg3)

Lipid MS/MS spectra are available at: https://doi.org/10.7910/DVN/MFKI8I Code developed can be retrieved at: https://github.com/yanisZirem/Spatial_multi-omics_guided_by_SVD_kmeans_clustering_and_statistical_estimation_of_heterogeneity.git

The data from Duhamel, M, *& al,.* for glioblastoma study, including MS raw files, MaxQuant files, and annotated MS/MS datasets, have been deposited to the ProteomeXchange Consortium Via PRIDE partner repository with the accession code PXD016165.

## SUPPLEMENTAL DATA

Supplementary information’s represents the 36 supplementary figures.

## ACKNOLEDGMENTS

This research was supported by grants from Ministère de l’Enseignement Supérieur et de la Recherche (MESR), Inserm specific funding for SpiderMass project (I.F.)**, Regional Council Hauts de France, The Metropole Europeenne de Lille (MEL) and I-site ULNE (grant MutiOmics)**, Inserm and Institut Universitaire de France (I.F. and M.S.). L.R. PhD was funded by University of Lille. Y.Z. is supported by ANR Click-Detect AAP CE29 (designed and built the machine learning workflow). L.L., Y.Z., I.F., M.S. and E.LR. corrected the manuscript. I.F. and M.S. supervised the project and provided the funding.

