## Supplementary material for "Heterogeneity Assessment and Protein Pathway Prediction via Spatial Lipidomic and Proteomic Correlation: Advancing Dry Proteomics concept for Human Glioblastoma Prognosis": Supplemntary Figures Legend

*Co-corresponding

**SUPPLEMENTARY FIGURES LEGENDS**

**Figure S1:** Rat brain annotations. A) Sagittal rat brain section annotations. B) Cerebellum rat brain sagittal section annotations. C) Rat brain sagittal section HPS coloration. D) Horizontal rat brain annotations. E) Horizontal rat brain section HPS coloration.

**Figure S2**: Cerebellum rat brain WM, ML and GL mean spectra and t-SNE separation for A) Lipid (-), B) Lipid (+), C) Protein, and D) Peptide MSI.

**Figure S3**: Comparison of different SCILS clustering methods applied to lipid negative mode, lipid positive mode, protein and peptide imaging. A) Scan of rat brain cerebellum analysed tissues and mean MSI spectra. Segmented images for each omics MSI analysis processed with B) Hierarchical clustering, C) Bisecting K-means with correlation distance, D) Bisecting K-means with Euclidian distance, E) K-means with correlation distance for 2 to 5 clusters, or E) Bisecting K-means with Euclidian distance for 2 to 5 clusters.

**Figure S4:** Unsupervised number of cluster selection according Silhouette interrogation applied to three rat brain cerebellum lipid, protein and peptide imaging data.

**Figure S5:** Lipid imaging of 22 rat brain cerebellum in negative mode with segmentation for 2 to 10 clusters.

**Figure S6:** Lipid imaging of 22 rat brain cerebellum in positive mode with segmentation for 2 to 10 clusters.

**Figure S7:** Lipid imaging of rat brain cerebellum in positive mode with DHB and segmentation for 2 to 10 clusters.

**Figure S8:** Protein imaging of 12 rat brain cerebellum with segmentation for 2 to 10 clusters.

**Figure S9:** Peptide imaging of 12 rat brain cerebellum with segmentation for 2 to 10 clusters.

**Figure S10:** Rat brain lipid negative mode A) peak picking datasets, followed by B) unsupervised discriminant ion list , and radar plot of the selected discriminant ions involved in C) GL, D) WM, or E) ML area .

**Figure S11:** Rat brain lipid positive mode A) peak picking datasets, followed by B) unsupervised discriminant ion list , and radar plot of the selected discriminant ions involved in C) GL, D) WM, or E) ML area .

**Figure S12:** Lipid (-) discriminant ions spatial distribution.

**Figure S13:** Lipid (+) discriminant ions spatial distribution.

**Figure S14:** Spidermas MS spectra for lipids in negative mode and positive mode in each cerebellum areas

**Figure S15:** Spidermas MS/MS spectra and discriminant ions annotations for lipids in A) negative mode, and B) positive mode examples.

**Figure S16 :** Rat brain lipid negative mode A) prediction model selection, with the B) 5-fold cross validation based on the 22 RB cohort, and a C) blind model validation performed on three rat brain MALDI MSI new datasets.

**Figure S17 :** Rat brain lipid positive mode A) prediction model selection, with the B) 5-fold cross validation based on the 22 RB cohort, and a C) blind model validation performed on three rat brain MALDI MSI new datasets.

**Figure S18:** Rat brain protein A) peak picking datasets, followed by B) unsupervised discriminant ion list , and discriminant ions spatial distribution in C).

**Figure S19:** Lipid pathway involved in white matter (WM) and gray matter (GM). A) Lipid subclass and species active pathway comparison between WM and granular matter GM discriminant lipids performed with BioPAN software. B) Lipid pathways involvment according REACTOME software.

**Figure S20**: ClueGO biological pathway involving proteins in A) white matter , B) gray matter and C) Molecular layer.

**Figure S21:** Rat brain horizontal section lipid (-), lipid (+), protein and peptide MSI segmentation images with 11 clusters and Silhouette criterion. A) Horizontal rat brain 1, B) Horizontal rat brain 2, C) Horizontal rat brain 3 and D) Horizontal rat brain 4.

**Figure S22:** White matter (WM), molecular layer (ML) and granular layer (GL) blind prediction on extracted rat brain horizontal section clusters from lipid (-) images. Results presented for the 4 different rat brain replicate.

**Figure S23:** Discriminant lipid (-) ions on rat brain horizontal section.

**Figure S24:** ClueGO biological pathways involving the significant proteins found in A) cerebellum, B) all clusters excluding cerebellum, C) Ventricular system, D) cerebral cortex and E) corpus callosum.

**Figure S25:** Robustness of A) spectral ions lipid (-) and B) protein between cluster triplicates with C) assessment in percentage.

**Figure S26:** Individual segmentation of 13 individual tumours by A) lipid and B) peptide MALDI-MSI and comparison with pathologist annotations.

**Figure S27:** Individual segmentation of 37 tumors by peptide MALDI-MSI and comparison with pathologist annotations.

**Figure S28:** ClueGO biological process and reactome pathways analysis for A) lipid cluster 1, B) lipid cluster 2, C) lipid cluster 4, D) lipid cluster 5, E) lipid cluster 6 and 10, F) lipid cluster 7, G) lipid cluster 8, H) lipid cluster 9, I) lipid cluster 12, and J) lipid cluster 13 according to K) lipid co-segmentatioin of 9 GBM tissue patient images.

**Figure S29:** ClueGO biological process and reactome pathways analysis for A) Tumoral Group A and B) Tumoral Group B.

**Figure S30:** Lipid cluster model prediction with A) 13 lipid cluster correlation heatmap highlighting group A and B; B) 5 fold cross validation based on 13 lipid clusters with C) correlation matrix. D) number of pixel involved in training model per cluster. E) Weight of lipid ions involved in the lipid cluster prediction model.

**Figure S31:** Top lipid biomarkers which contribute to each Group A cluster, statistically significant according Kruskal-Wallis test with p-value <0,005.

**Figure S32:** Top lipid biomarkers which contribute to each Group B cluster, statistically significant according Kruskal-Wallis test with p-value <0,005.

**Figure S33:** Blind cluster prediction involved for patient 3, 5 , 6 and 11 according A) lipid MSI prediction model and B) 13 lipid patient co-segmentation.

**Figure S34:** 13 lipid GBM tissue patient co-segmentation composed by 14 clusters.

**Figure S35:** Top proteinID biomarkers which contribute to each Group A and B cluster, statistically significant according Kruskal-Wallis test with p-value <0,005.

**Figure S36:** Blind survival prognosis prediction for the 50 GBM patients.
