## Supplementary material for "Heterogeneity Assessment and Protein Pathway Prediction via Spatial Lipidomic and Proteomic Correlation: Advancing Dry Proteomics concept for Human Glioblastoma Prognosis": Supplemntary data

### Slide 1
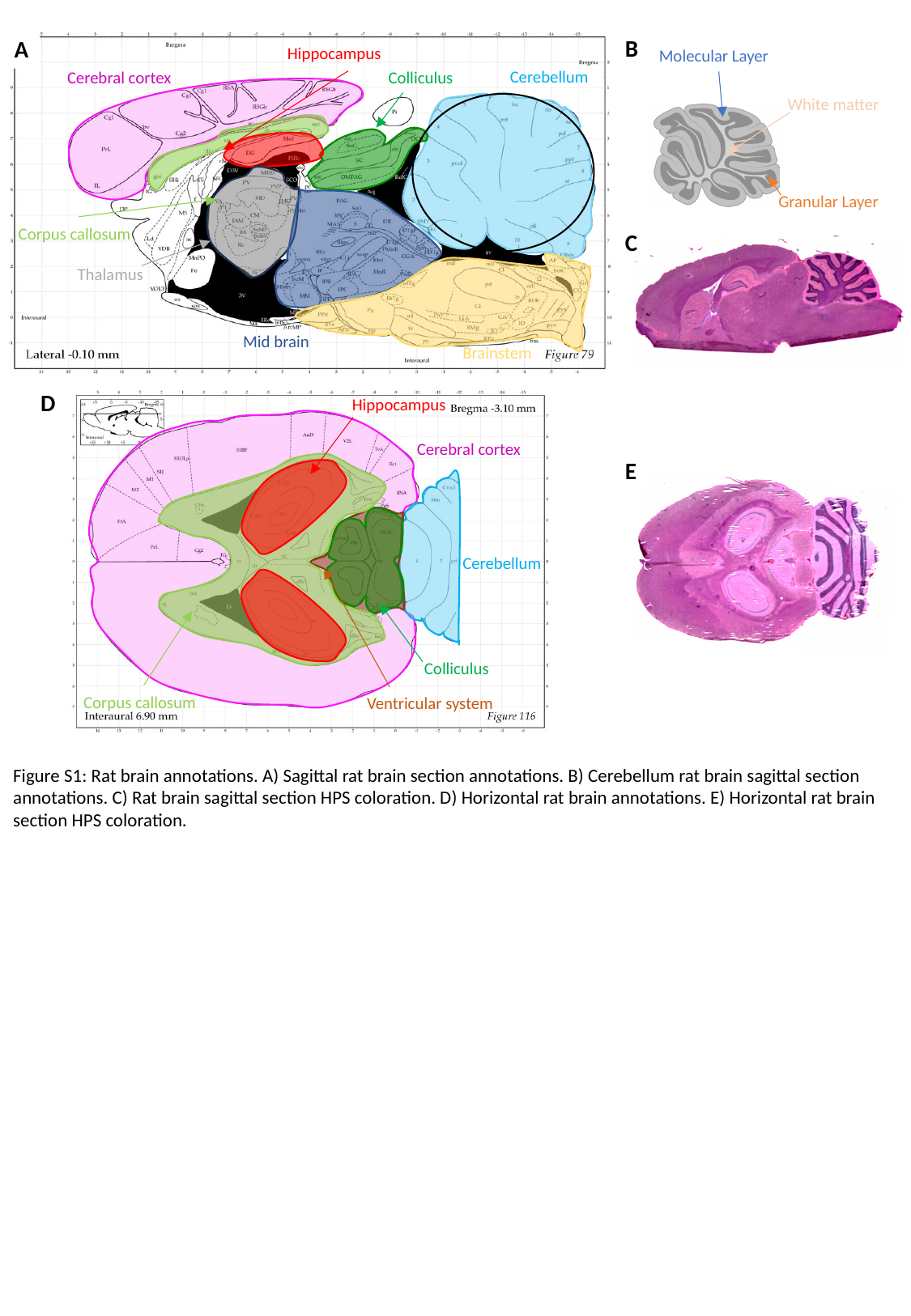

B
A
C
Hippocampus
Molecular Layer
White matter
Granular Layer
Cerebellum
Colliculus
Cerebral cortex
Corpus callosum
Thalamus
Mid brain
Brainstem
D
Hippocampus
Cerebral cortex
E
Cerebellum
Colliculus
Corpus callosum
Ventricular system
Figure S1: Rat brain annotations. A) Sagittal rat brain section annotations. B) Cerebellum rat brain sagittal section annotations. C) Rat brain sagittal section HPS coloration. D) Horizontal rat brain annotations. E) Horizontal rat brain section HPS coloration.

### Slide 2
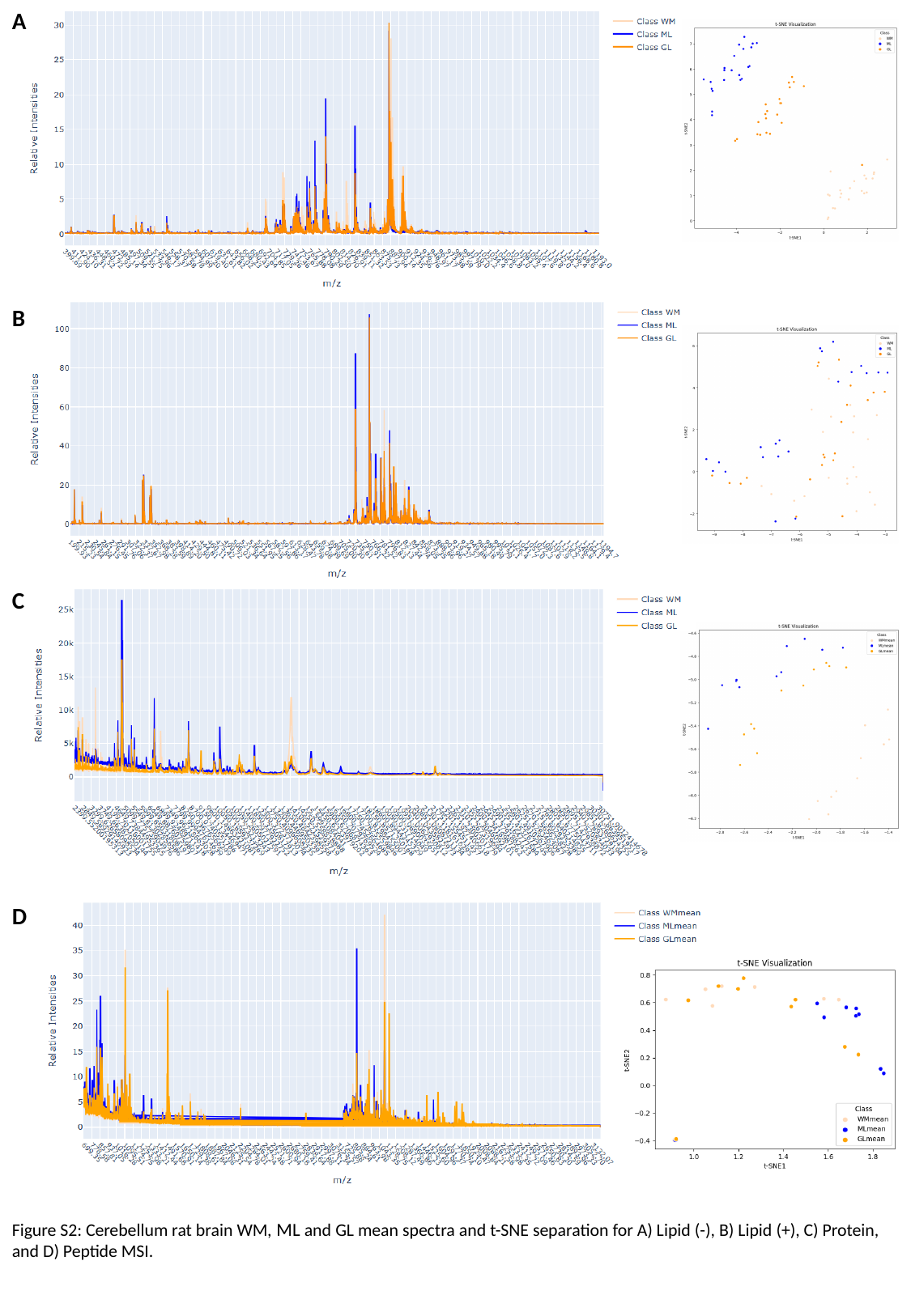

A
B
C
D
Figure S2: Cerebellum rat brain WM, ML and GL mean spectra and t-SNE separation for A) Lipid (-), B) Lipid (+), C) Protein, and D) Peptide MSI.

### Slide 3
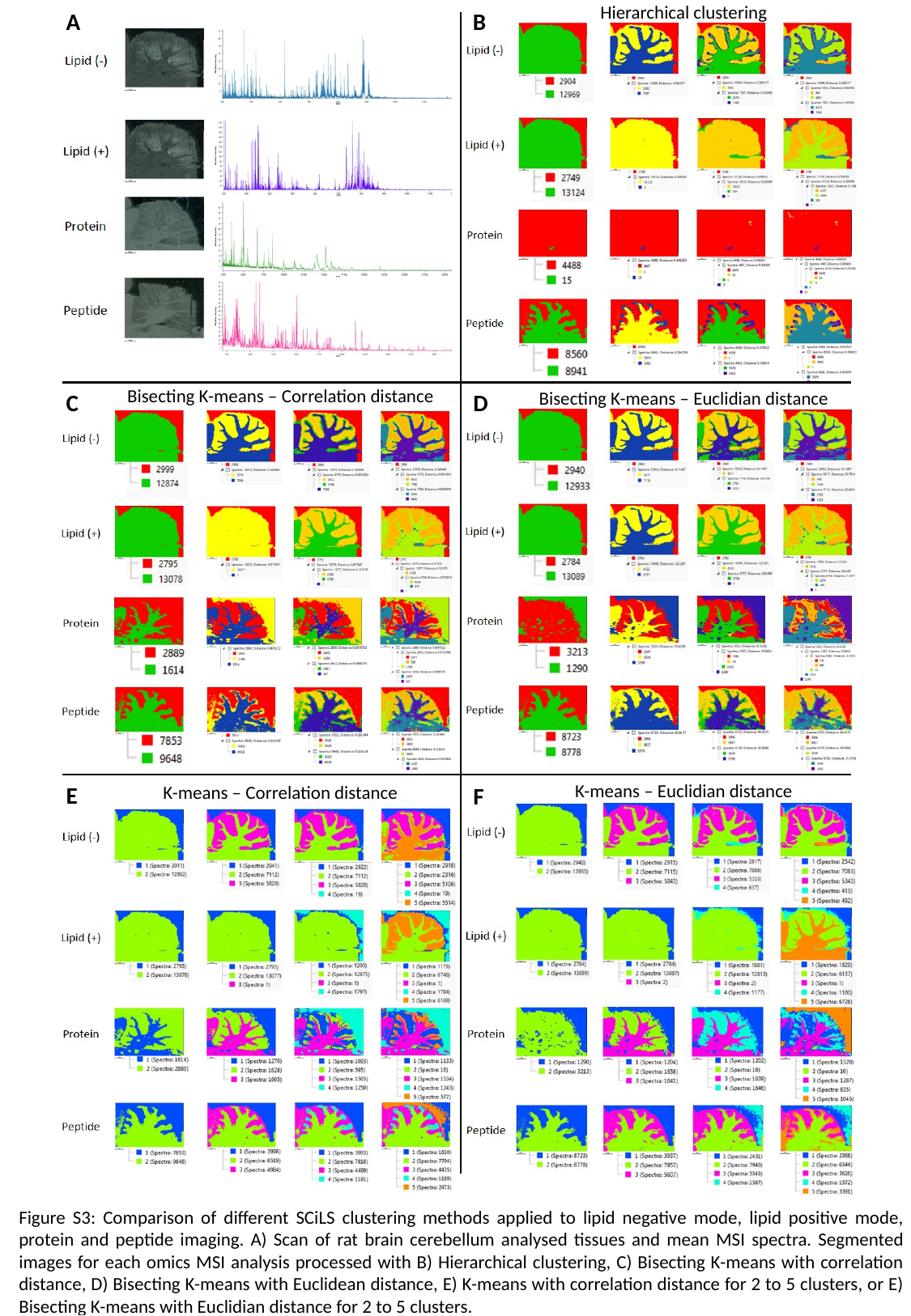

Hierarchical clustering
A
B
Bisecting K-means – Correlation distance
Bisecting K-means – Euclidian distance
C
D
K-means – Euclidian distance
E
K-means – Correlation distance
F
Figure S3: Comparison of different SCiLS clustering methods applied to lipid negative mode, lipid positive mode, protein and peptide imaging. A) Scan of rat brain cerebellum analysed tissues and mean MSI spectra. Segmented images for each omics MSI analysis processed with B) Hierarchical clustering, C) Bisecting K-means with correlation distance, D) Bisecting K-means with Euclidean distance, E) K-means with correlation distance for 2 to 5 clusters, or E) Bisecting K-means with Euclidian distance for 2 to 5 clusters.

### Slide 4
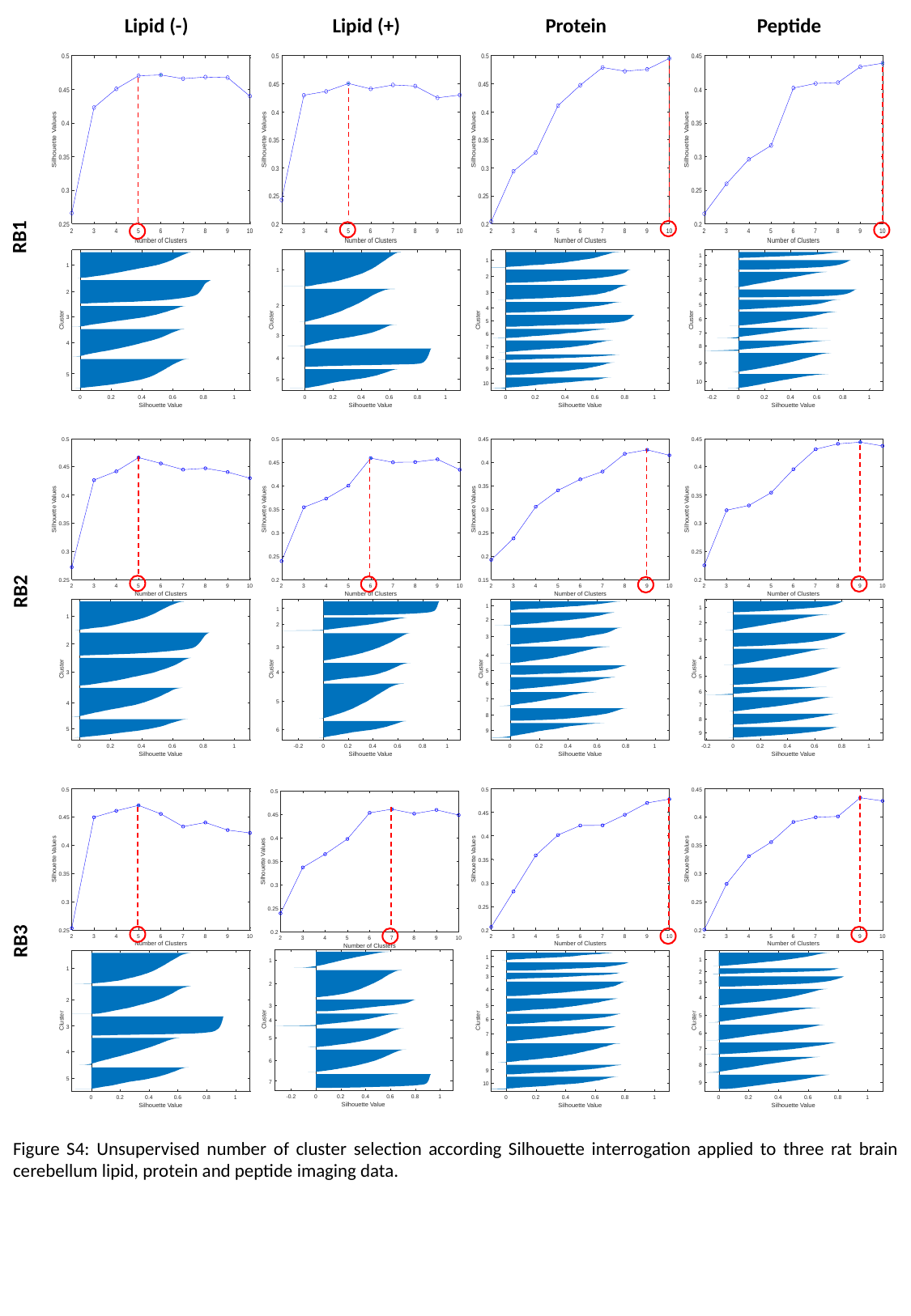

Lipid (-)
Lipid (+)
Protein
Peptide
RB1
RB2
RB3
Figure S4: Unsupervised number of cluster selection according Silhouette interrogation applied to three rat brain cerebellum lipid, protein and peptide imaging data.

### Slide 5
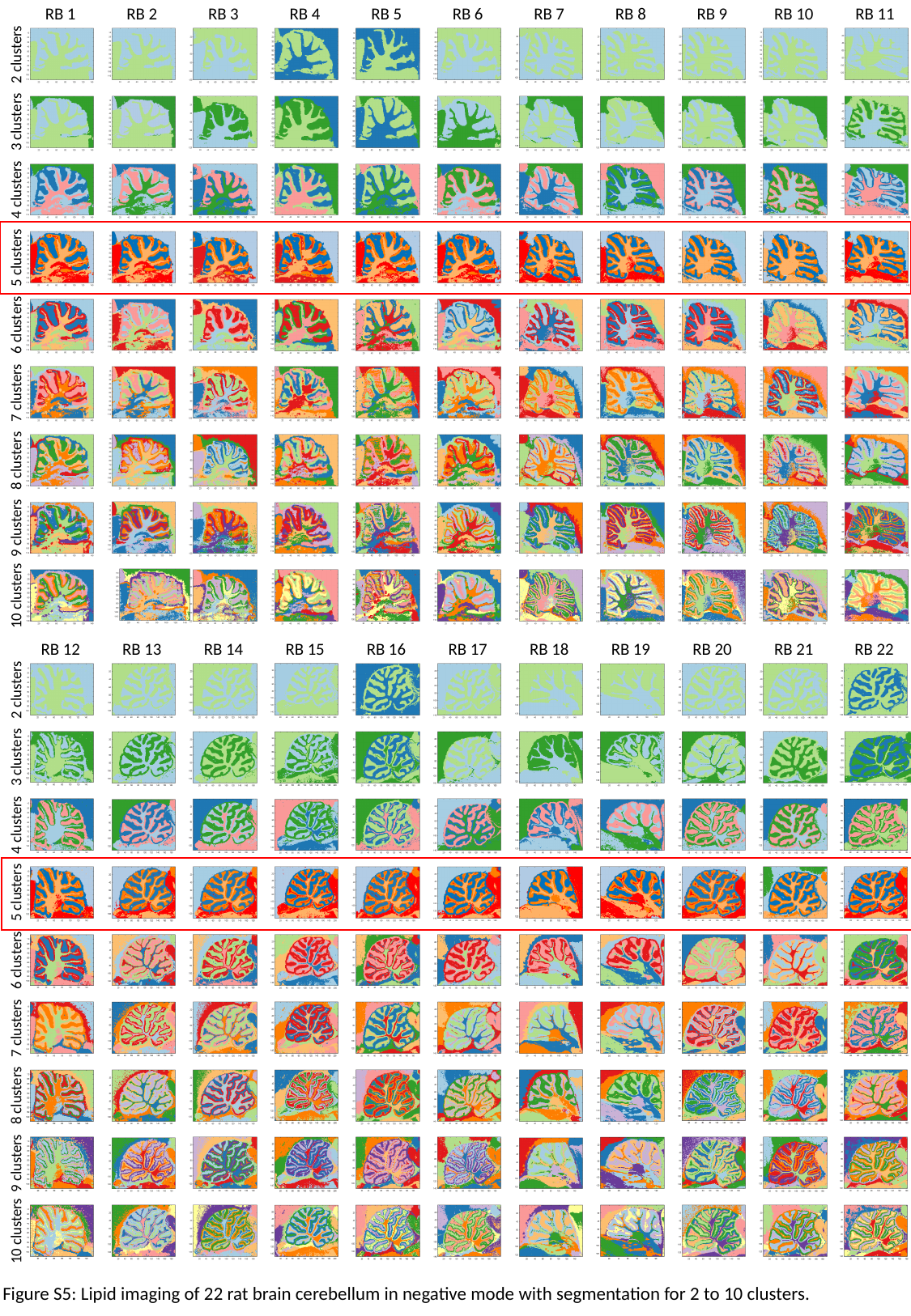

RB 1
RB 2
RB 3
RB 4
RB 5
RB 6
RB 7
RB 8
RB 9
RB 10
RB 11
2 clusters
3 clusters
4 clusters
5 clusters
6 clusters
7 clusters
8 clusters
9 clusters
10 clusters
RB 12
RB 13
RB 14
RB 15
RB 16
RB 17
RB 18
RB 19
RB 20
RB 21
RB 22
2 clusters
3 clusters
4 clusters
5 clusters
6 clusters
7 clusters
8 clusters
9 clusters
10 clusters
Figure S5: Lipid imaging of 22 rat brain cerebellum in negative mode with segmentation for 2 to 10 clusters.

### Slide 6
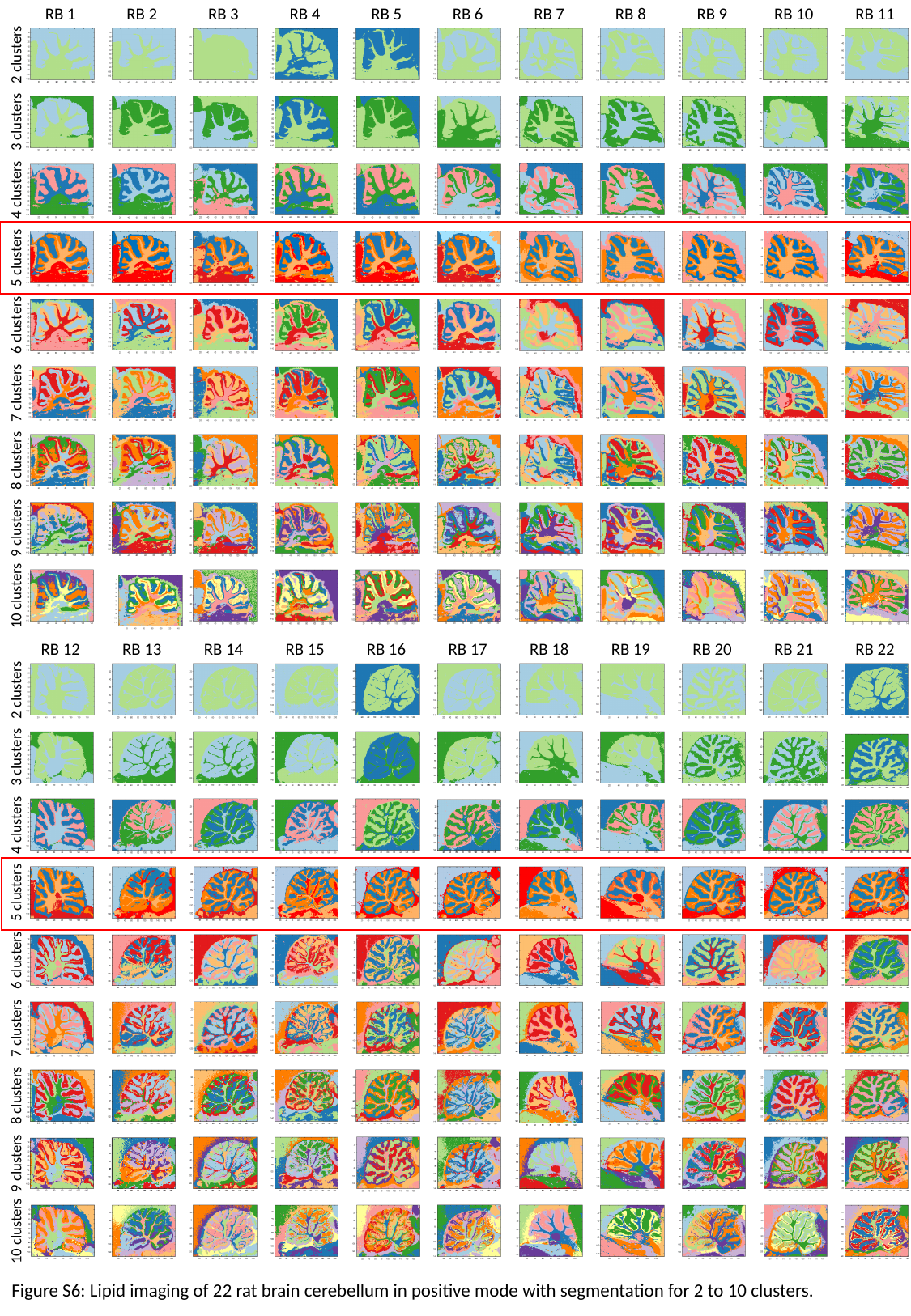

RB 1
RB 2
RB 3
RB 4
RB 5
RB 6
RB 7
RB 8
RB 9
RB 10
RB 11
2 clusters
3 clusters
4 clusters
5 clusters
6 clusters
7 clusters
8 clusters
9 clusters
10 clusters
RB 12
RB 13
RB 14
RB 15
RB 16
RB 17
RB 18
RB 19
RB 20
RB 21
RB 22
2 clusters
3 clusters
4 clusters
5 clusters
6 clusters
7 clusters
8 clusters
9 clusters
10 clusters
Figure S6: Lipid imaging of 22 rat brain cerebellum in positive mode with segmentation for 2 to 10 clusters.

### Slide 7
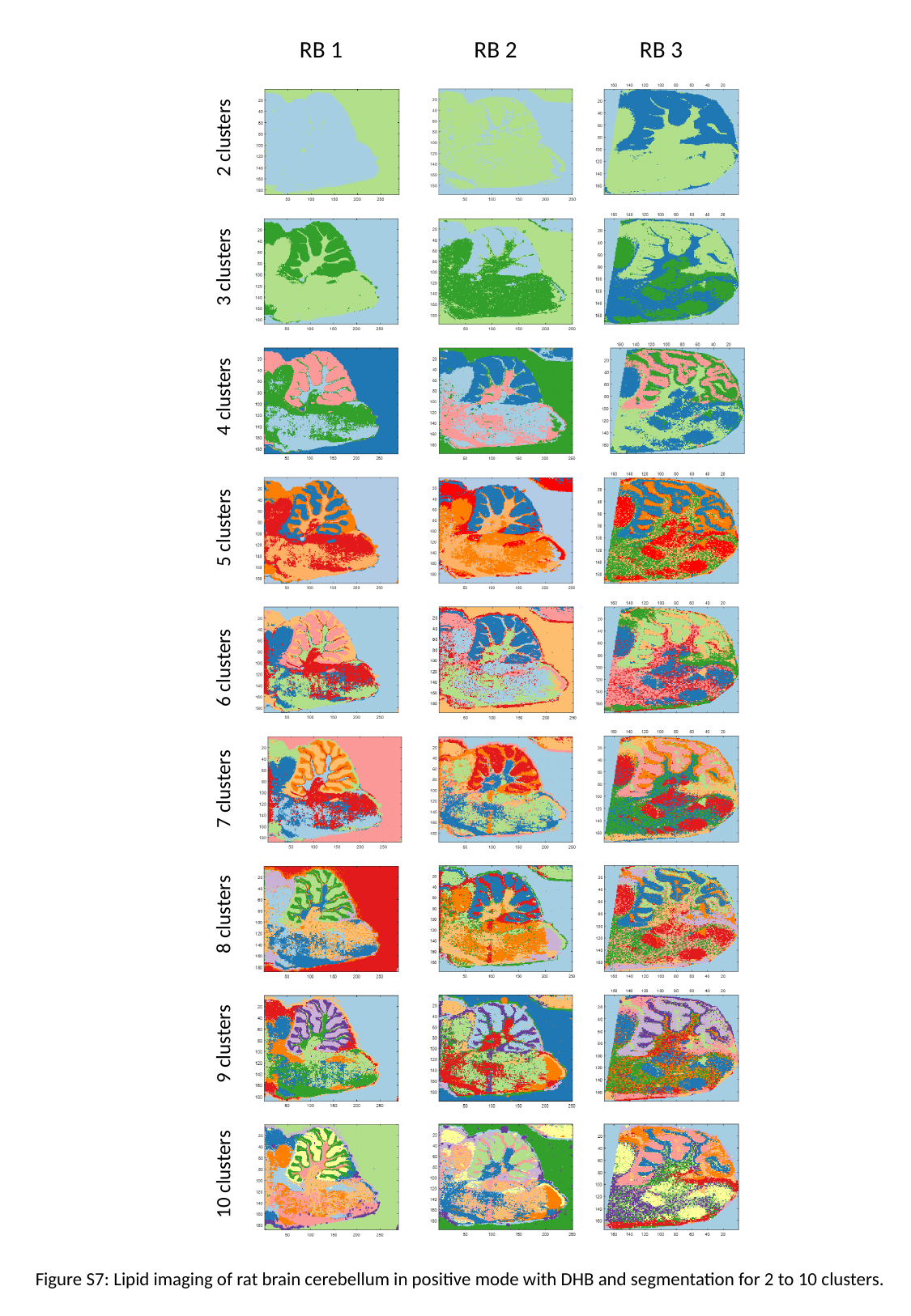

RB 1
RB 2
RB 3
2 clusters
3 clusters
4 clusters
5 clusters
6 clusters
7 clusters
8 clusters
9 clusters
10 clusters
Figure S7: Lipid imaging of rat brain cerebellum in positive mode with DHB and segmentation for 2 to 10 clusters.

### Slide 8
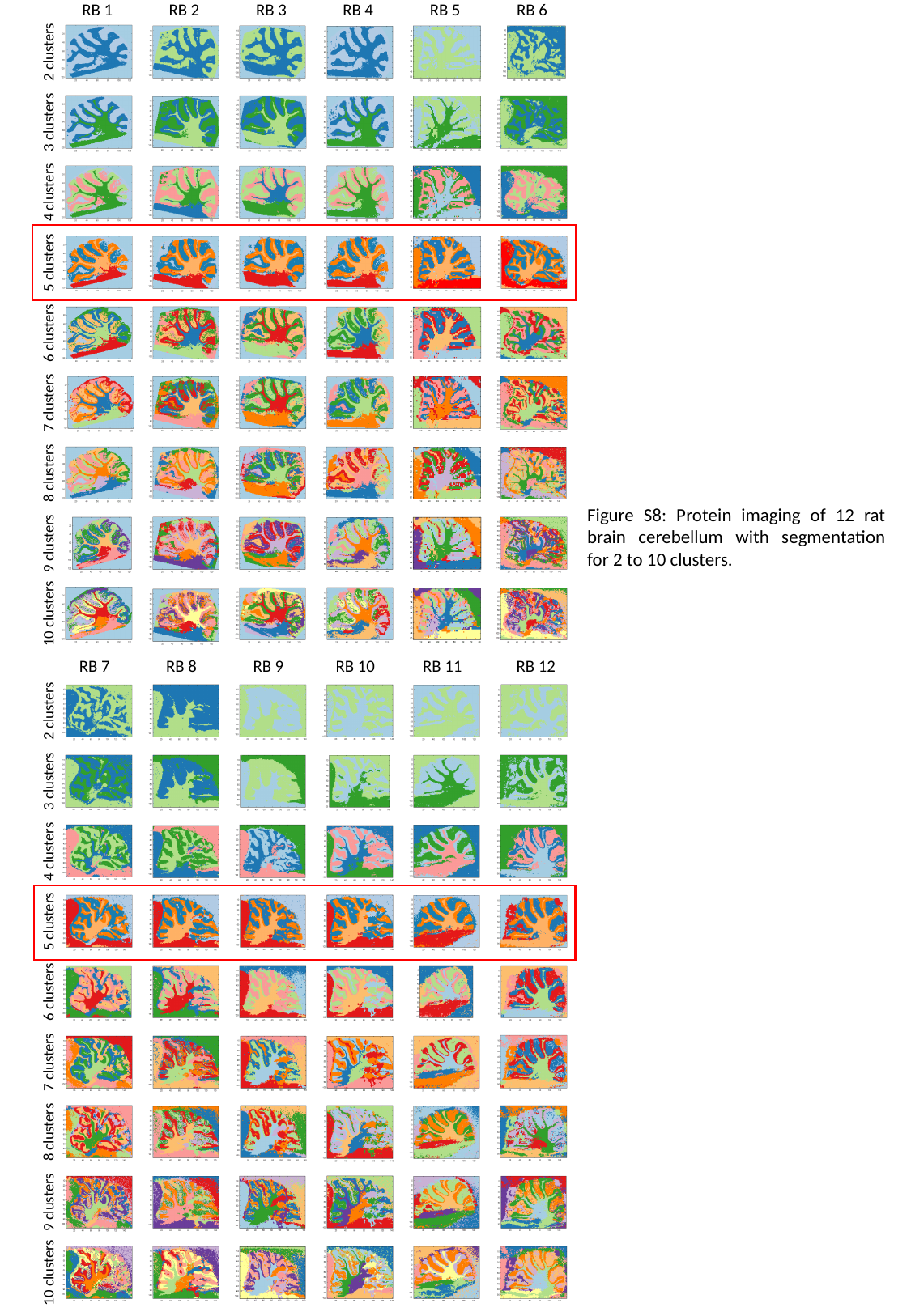

RB 1
RB 2
RB 3
RB 4
RB 5
RB 6
2 clusters
3 clusters
4 clusters
5 clusters
6 clusters
7 clusters
8 clusters
9 clusters
10 clusters
RB 7
RB 8
RB 9
RB 10
RB 11
RB 12
2 clusters
3 clusters
4 clusters
5 clusters
6 clusters
7 clusters
8 clusters
9 clusters
10 clusters
Figure S8: Protein imaging of 12 rat brain cerebellum with segmentation for 2 to 10 clusters.

### Slide 9
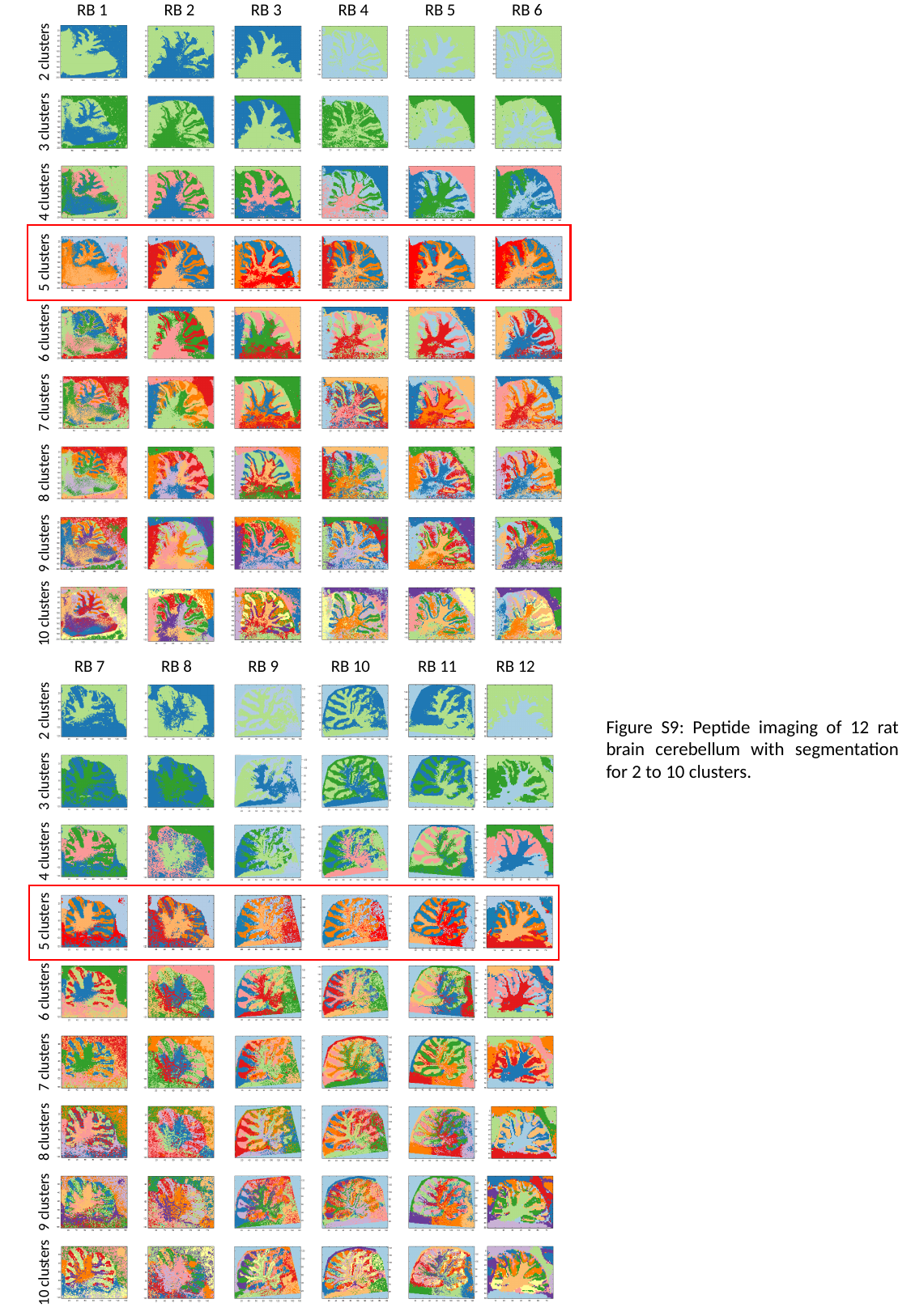

RB 1
RB 2
RB 3
RB 4
RB 5
RB 6
2 clusters
3 clusters
4 clusters
5 clusters
6 clusters
7 clusters
8 clusters
9 clusters
10 clusters
RB 7
RB 8
RB 9
RB 10
RB 11
RB 12
2 clusters
Figure S9: Peptide imaging of 12 rat brain cerebellum with segmentation for 2 to 10 clusters.
3 clusters
4 clusters
5 clusters
6 clusters
7 clusters
8 clusters
9 clusters
10 clusters

### Slide 10
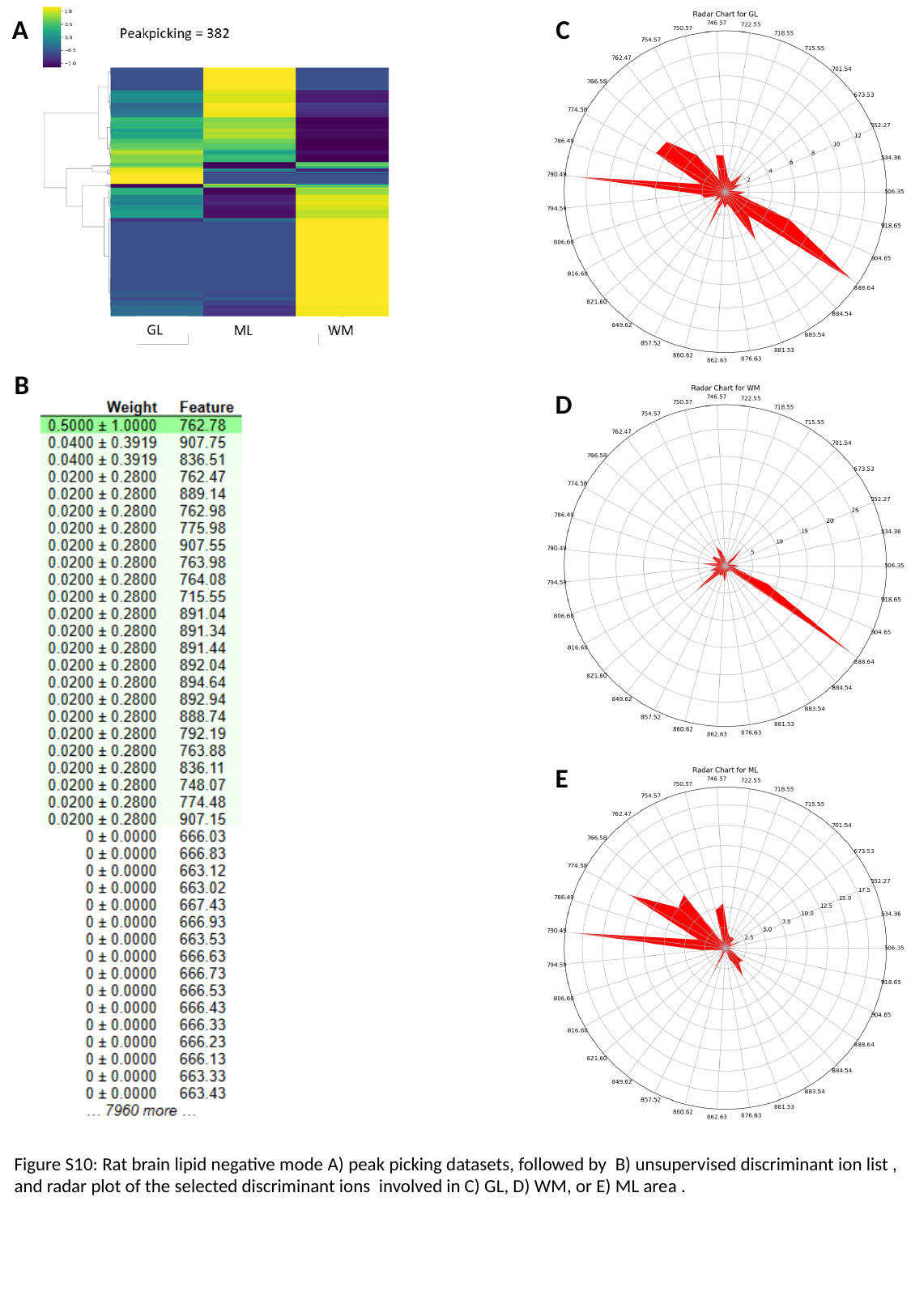

A
C
B
D
E
Figure S10: Rat brain lipid negative mode A) peak picking datasets, followed by B) unsupervised discriminant ion list , and radar plot of the selected discriminant ions involved in C) GL, D) WM, or E) ML area .

### Slide 11
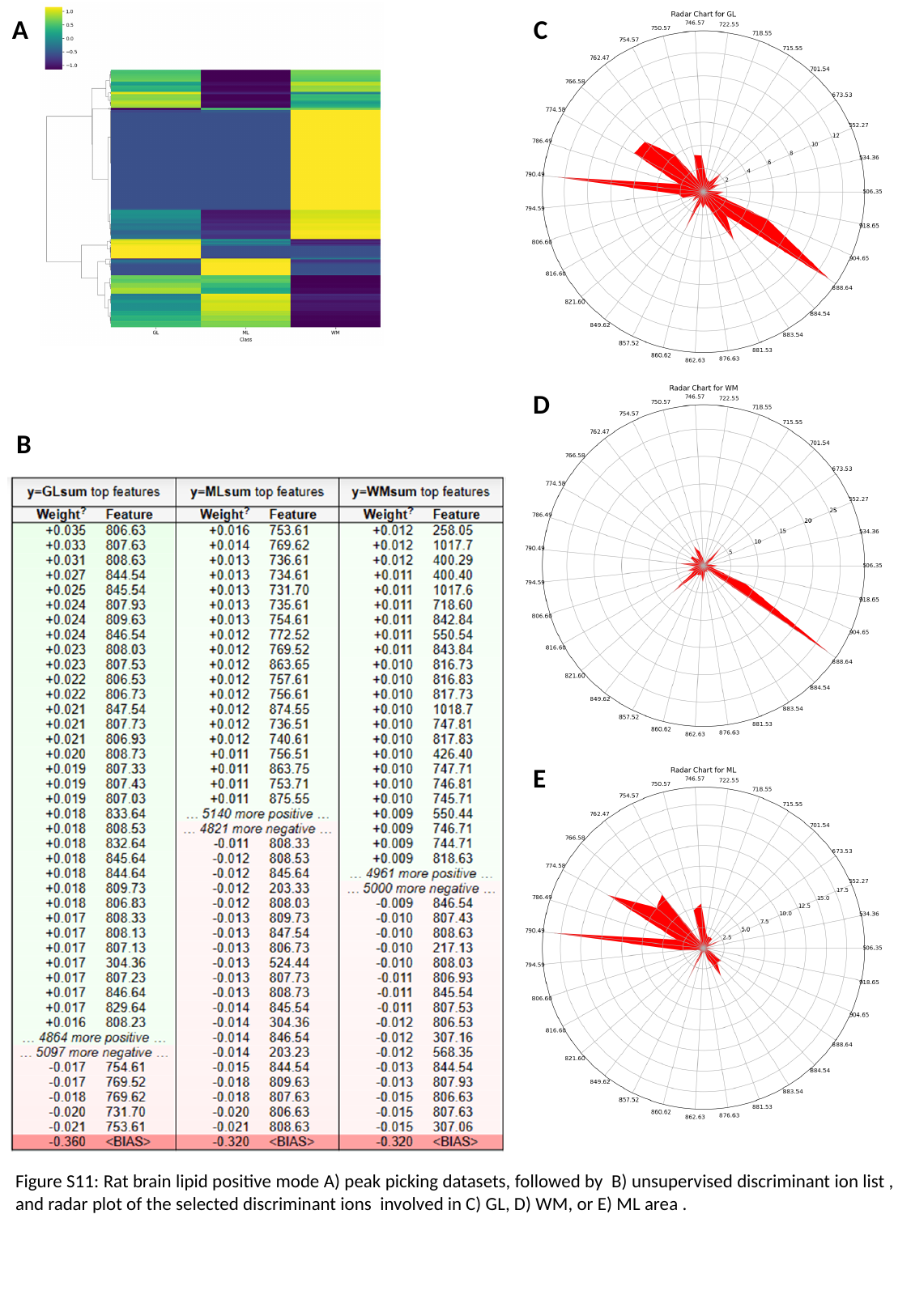

A
C
D
B
E
Figure S11: Rat brain lipid positive mode A) peak picking datasets, followed by B) unsupervised discriminant ion list , and radar plot of the selected discriminant ions involved in C) GL, D) WM, or E) ML area .

### Slide 12
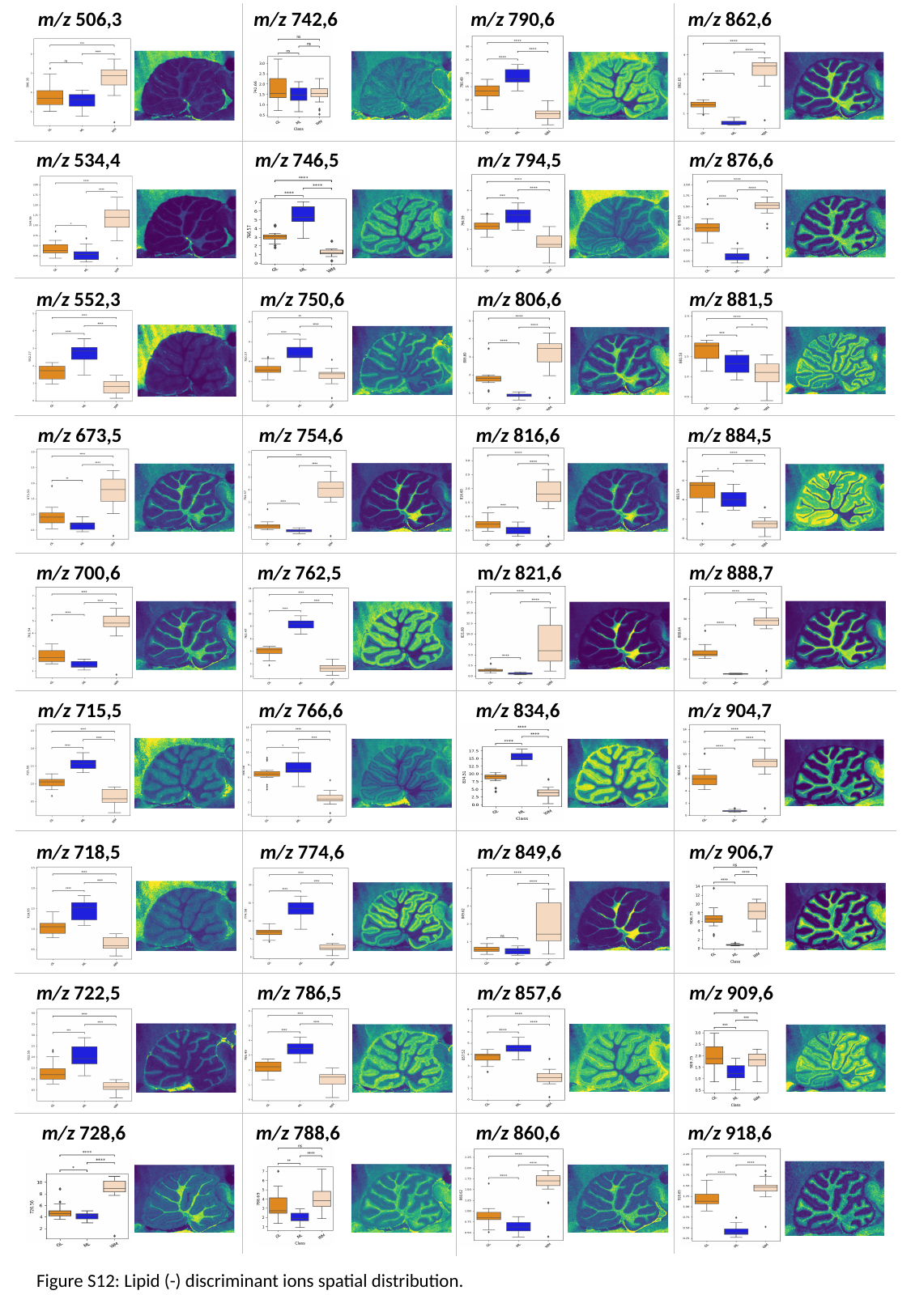

m/z 506,3
m/z 742,6
m/z 790,6
m/z 862,6
m/z 534,4
m/z 746,5
m/z 794,5
m/z 876,6
m/z 552,3
m/z 750,6
m/z 806,6
m/z 881,5
m/z 673,5
m/z 754,6
m/z 816,6
m/z 884,5
m/z 700,6
m/z 762,5
m/z 821,6
m/z 888,7
m/z 715,5
m/z 766,6
m/z 834,6
m/z 904,7
m/z 718,5
m/z 774,6
m/z 849,6
m/z 906,7
m/z 722,5
m/z 786,5
m/z 857,6
m/z 909,6
m/z 728,6
m/z 788,6
m/z 860,6
m/z 918,6
Figure S12: Lipid (-) discriminant ions spatial distribution.

### Slide 13
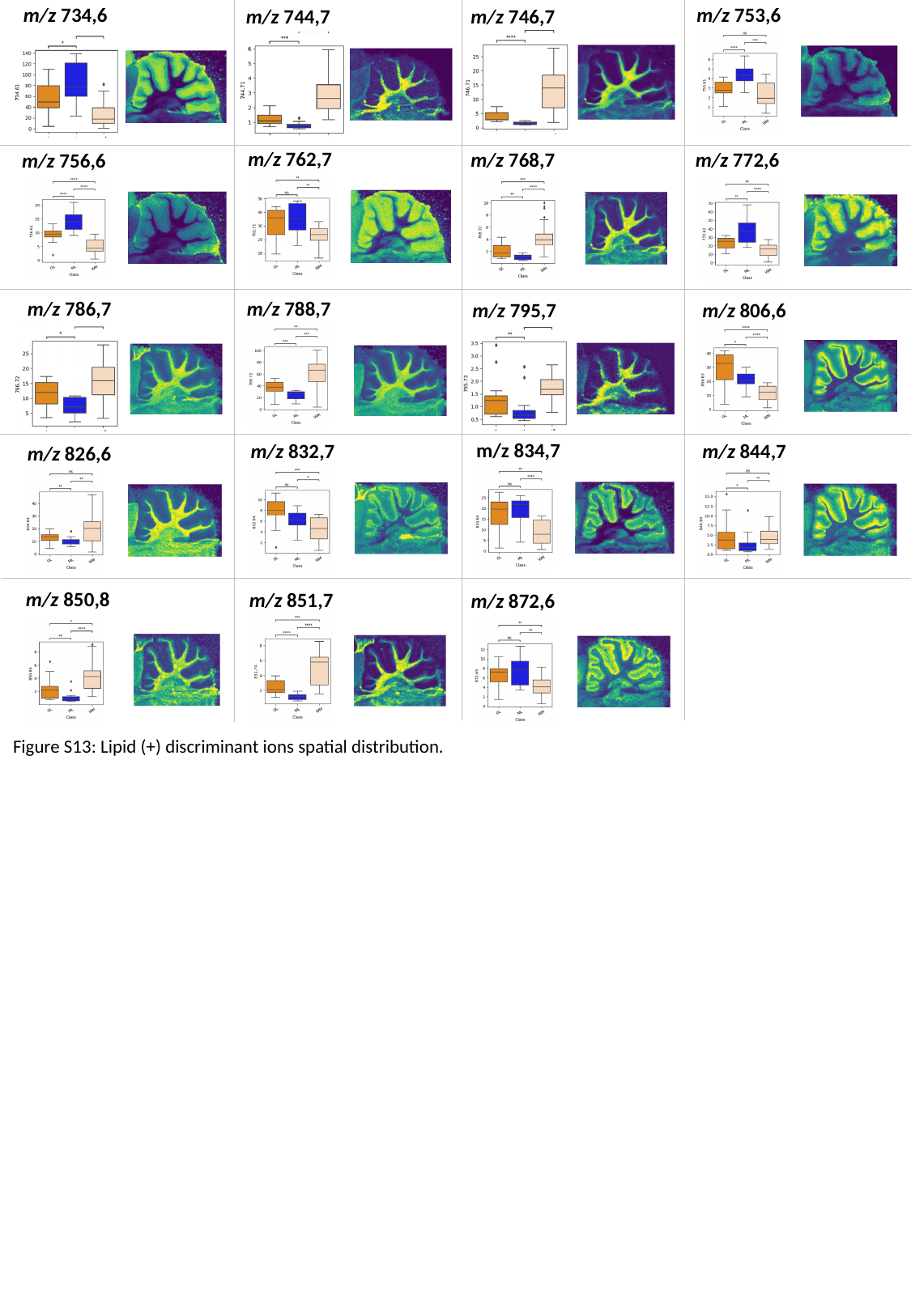

m/z 753,6
m/z 734,6
m/z 746,7
m/z 744,7
m/z 762,7
m/z 772,6
m/z 768,7
m/z 756,6
m/z 786,7
m/z 788,7
m/z 806,6
m/z 795,7
m/z 834,7
m/z 832,7
m/z 844,7
m/z 826,6
m/z 850,8
m/z 851,7
m/z 872,6
Figure S13: Lipid (+) discriminant ions spatial distribution.

### Slide 14
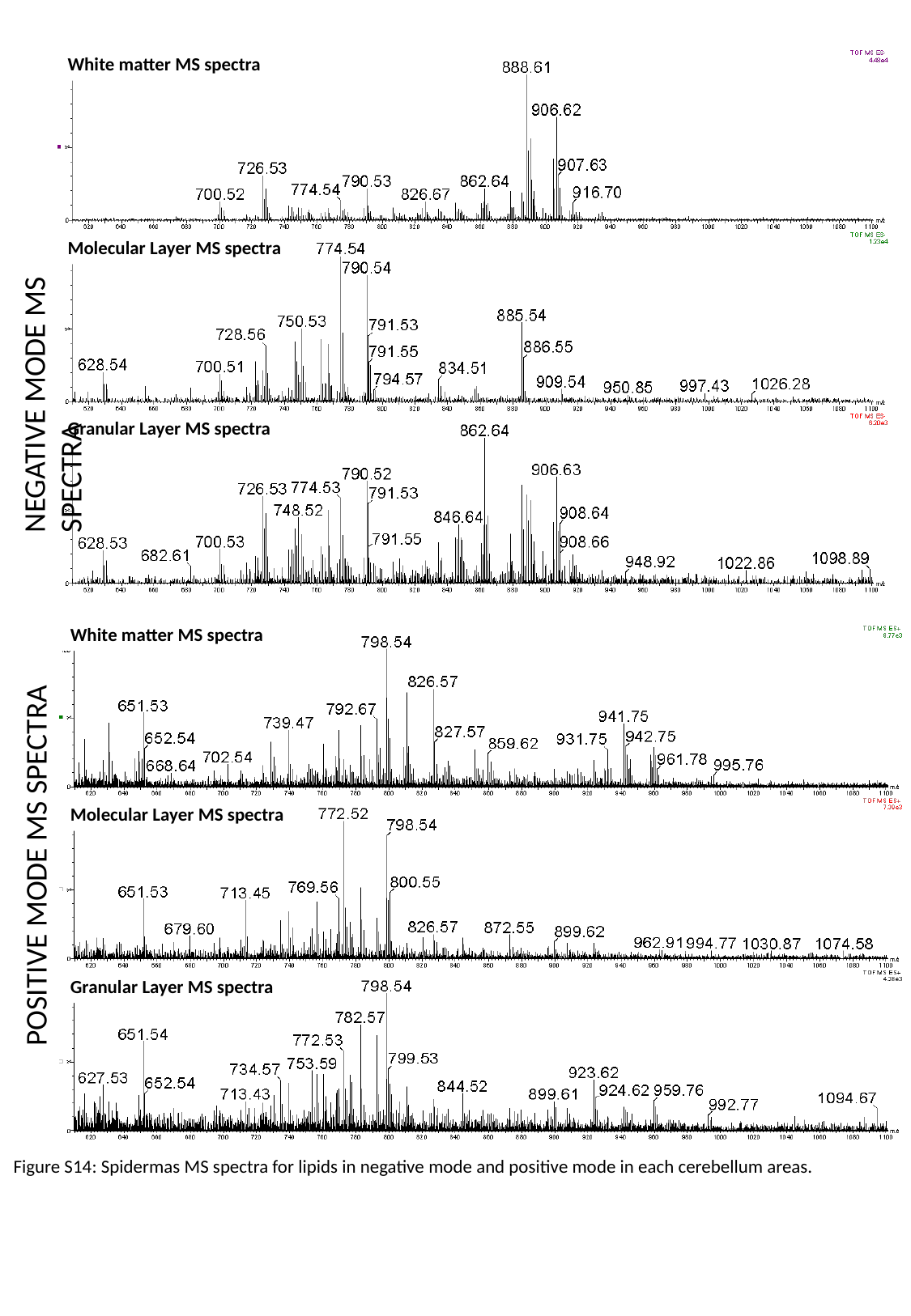

White matter MS spectra
NEGATIVE MODE MS SPECTRA
Molecular Layer MS spectra
Granular Layer MS spectra
White matter MS spectra
POSITIVE MODE MS SPECTRA
Molecular Layer MS spectra
Granular Layer MS spectra
Figure S14: Spidermas MS spectra for lipids in negative mode and positive mode in each cerebellum areas.

### Slide 15
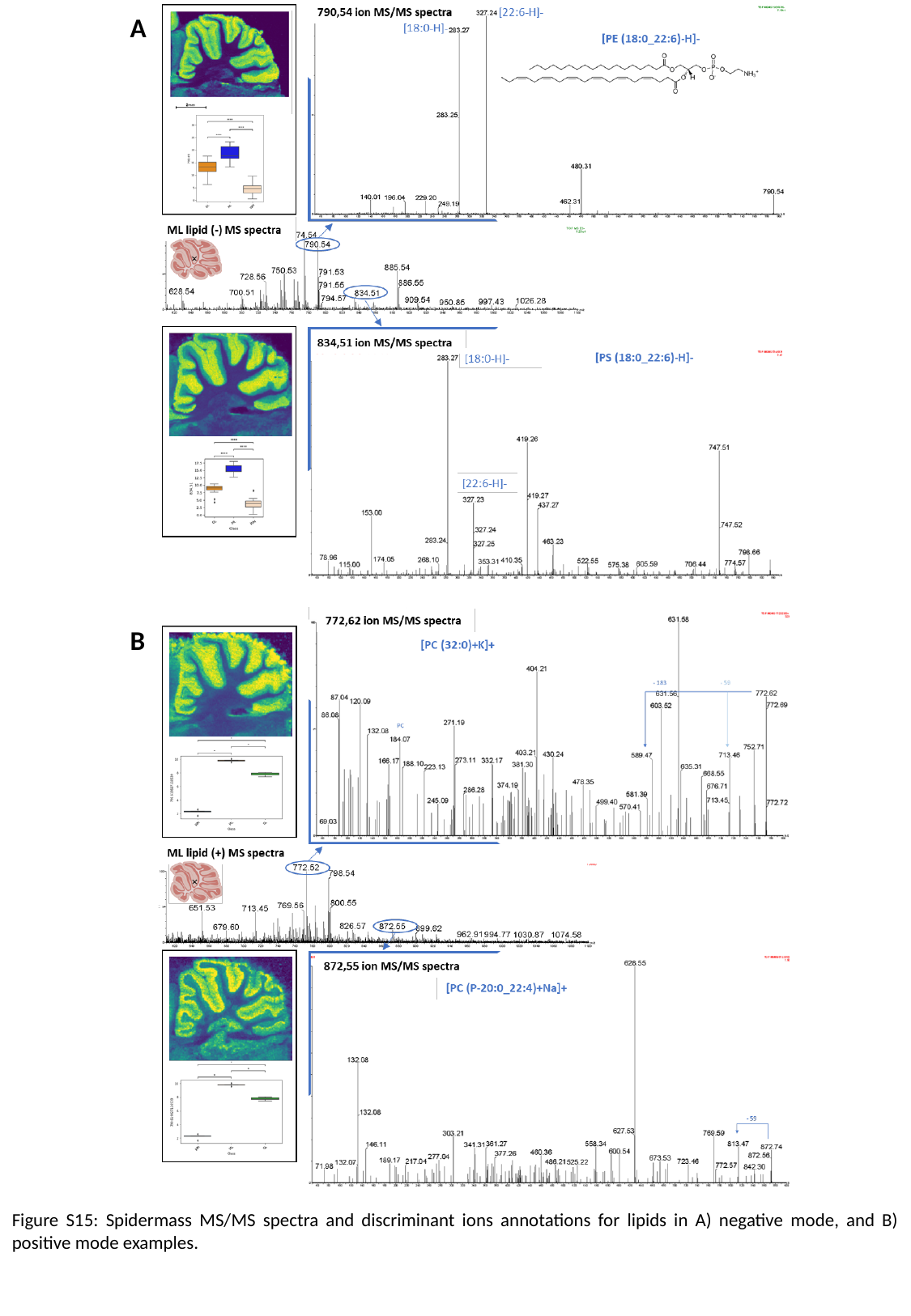

A
B
Figure S15: Spidermass MS/MS spectra and discriminant ions annotations for lipids in A) negative mode, and B) positive mode examples.

### Slide 16
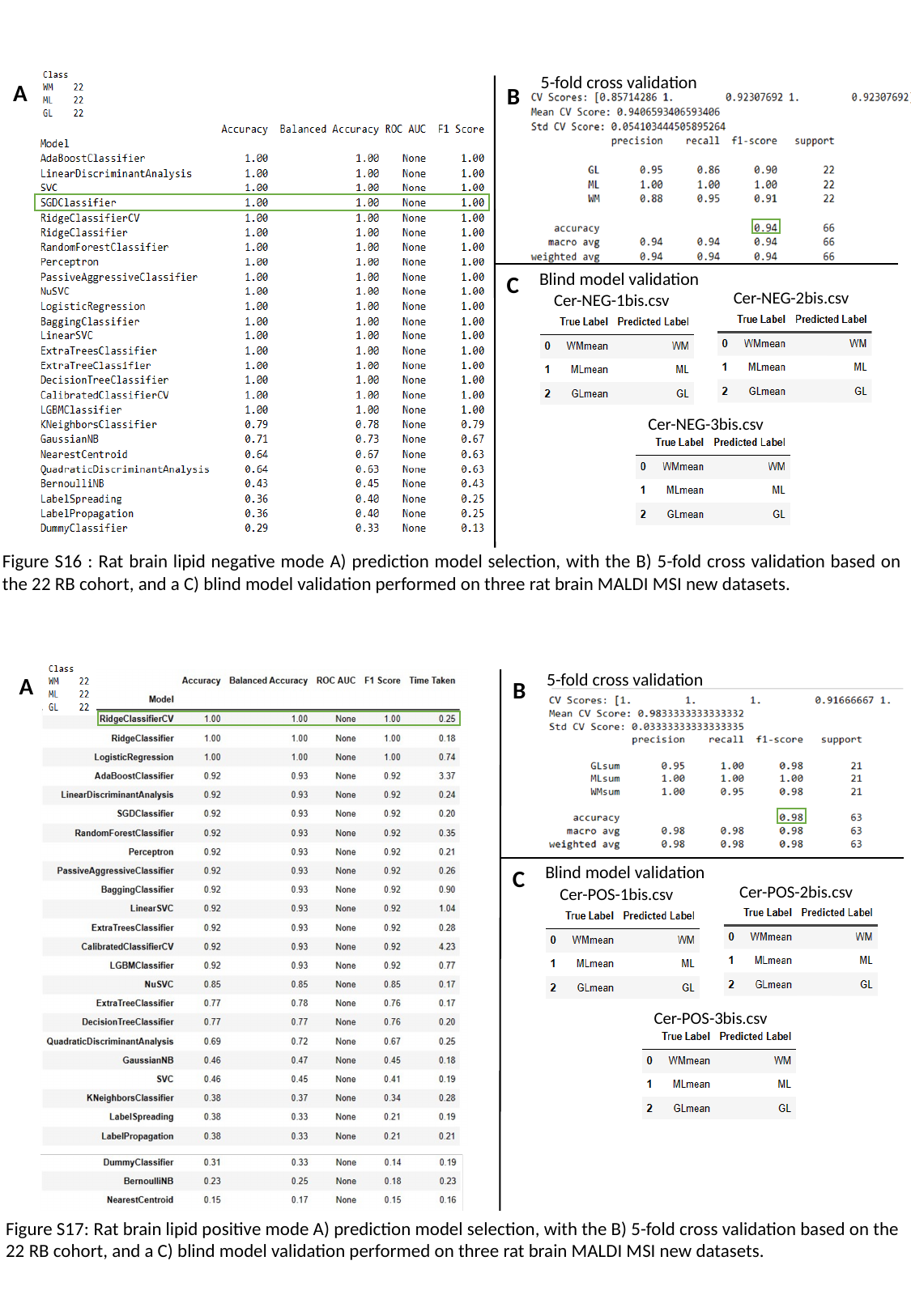

5-fold cross validation
A
B
Blind model validation
C
Cer-NEG-2bis.csv
Cer-NEG-1bis.csv
Cer-NEG-3bis.csv
Figure S16 : Rat brain lipid negative mode A) prediction model selection, with the B) 5-fold cross validation based on the 22 RB cohort, and a C) blind model validation performed on three rat brain MALDI MSI new datasets.
5-fold cross validation
A
B
Blind model validation
C
Cer-POS-2bis.csv
Cer-POS-1bis.csv
Cer-POS-3bis.csv
Figure S17: Rat brain lipid positive mode A) prediction model selection, with the B) 5-fold cross validation based on the 22 RB cohort, and a C) blind model validation performed on three rat brain MALDI MSI new datasets.

### Slide 17
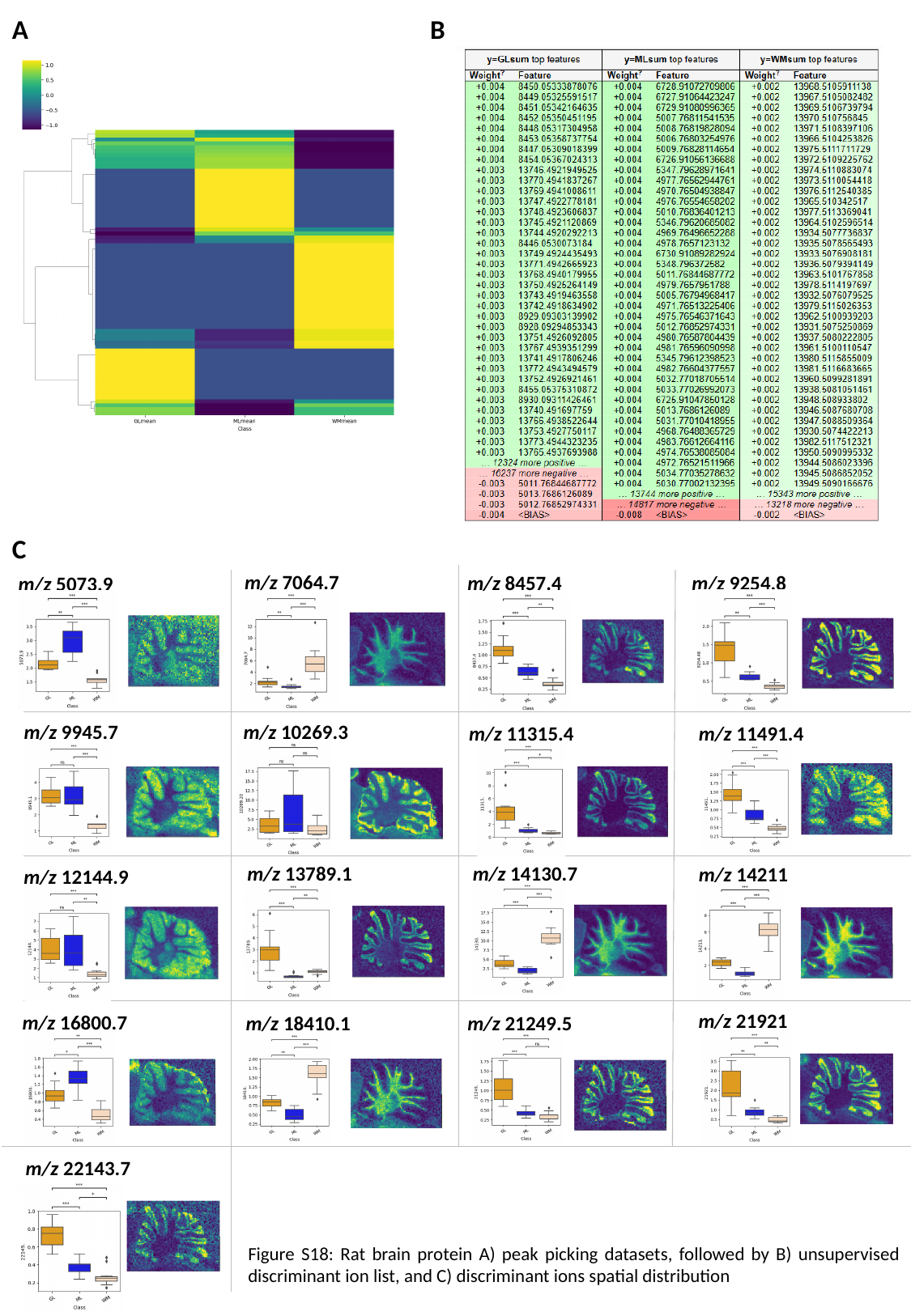

A
B
C
m/z 7064.7
m/z 9254.8
m/z 8457.4
m/z 5073.9
m/z 9945.7
m/z 10269.3
m/z 11491.4
m/z 11315.4
m/z 14130.7
m/z 13789.1
m/z 14211
m/z 12144.9
m/z 21921
m/z 16800.7
m/z 18410.1
m/z 21249.5
m/z 22143.7
Figure S18: Rat brain protein A) peak picking datasets, followed by B) unsupervised discriminant ion list, and C) discriminant ions spatial distribution

### Slide 18
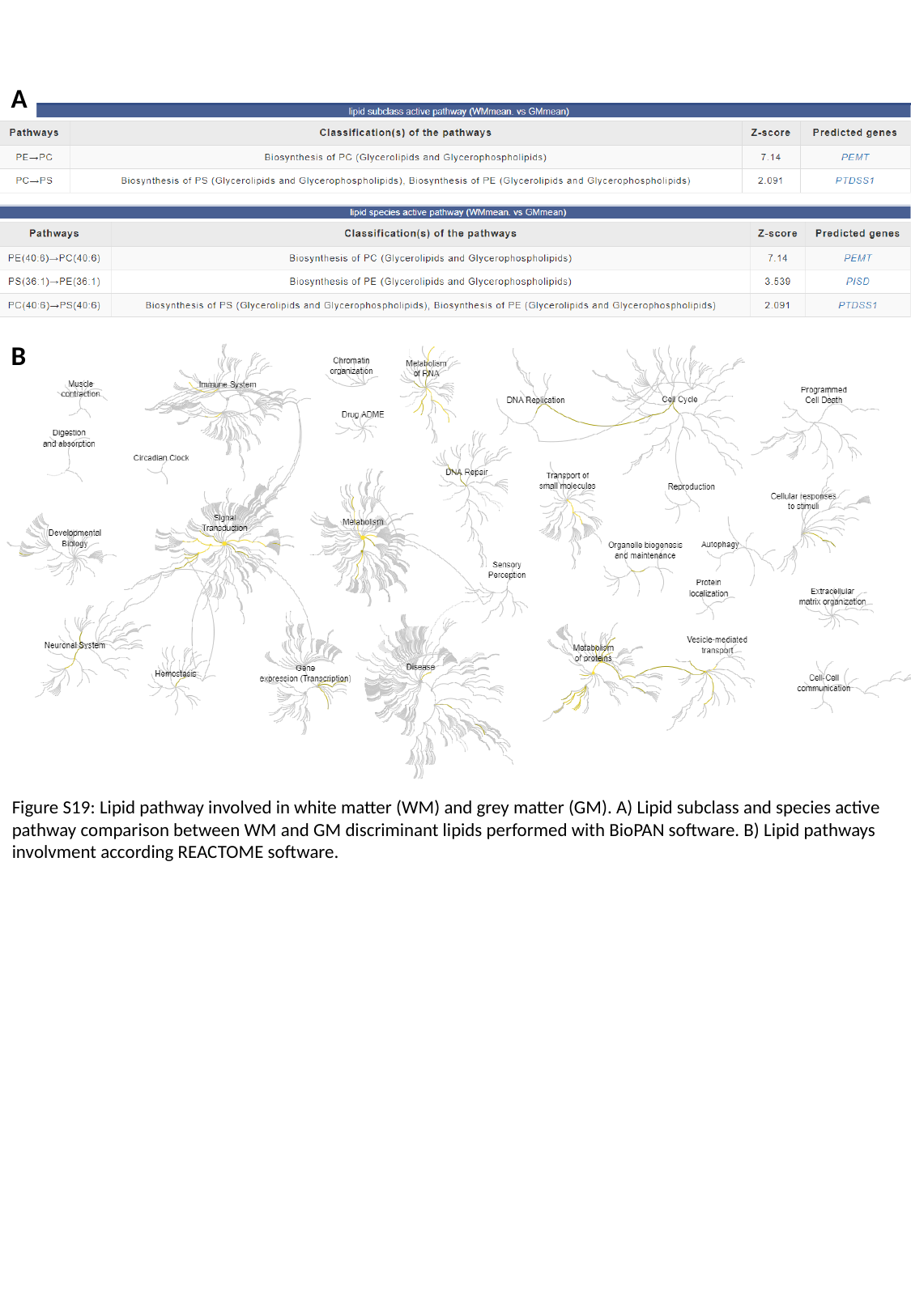

A
B
Figure S19: Lipid pathway involved in white matter (WM) and grey matter (GM). A) Lipid subclass and species active pathway comparison between WM and GM discriminant lipids performed with BioPAN software. B) Lipid pathways involvment according REACTOME software.

### Slide 19
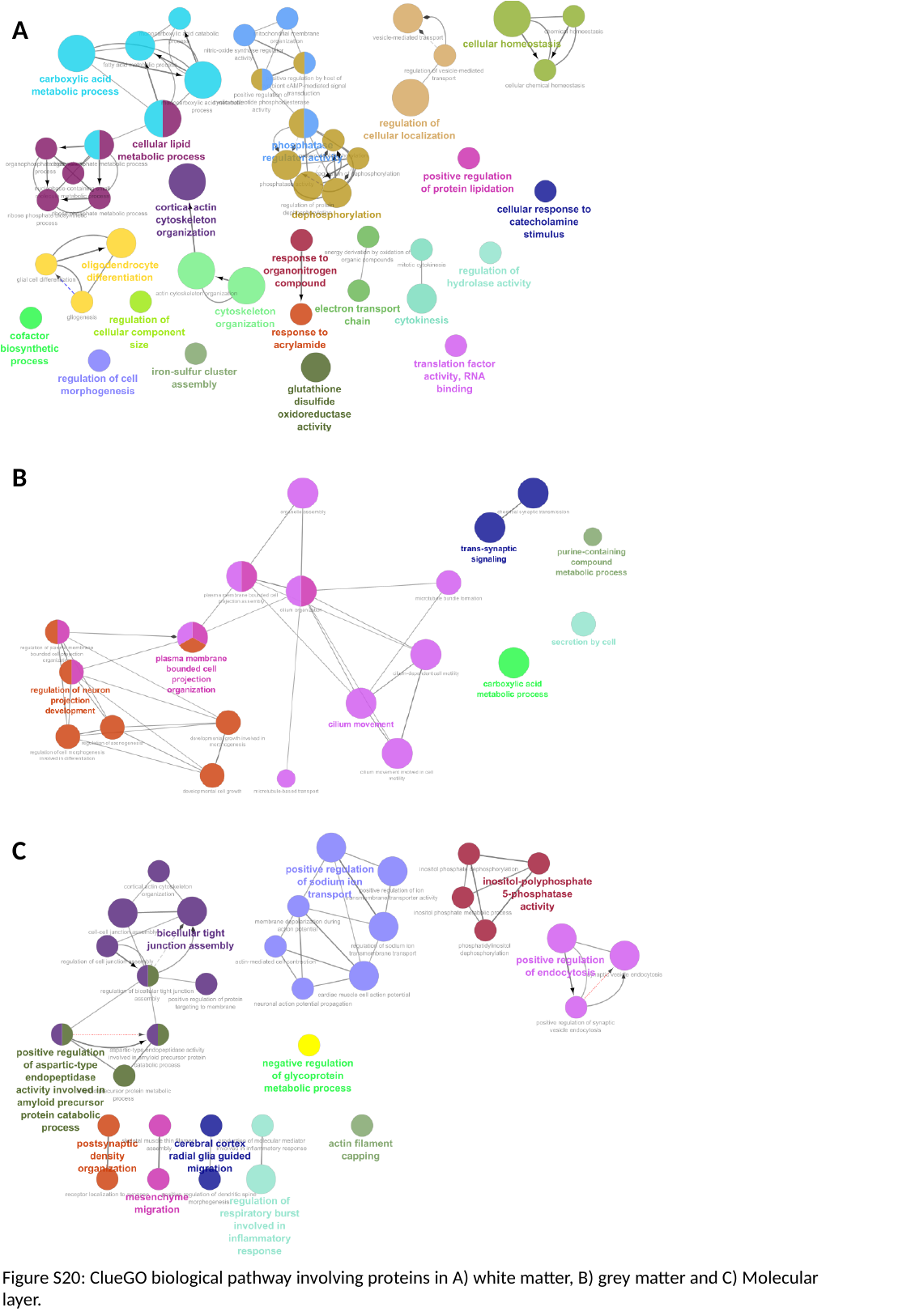

A
B
C
Figure S20: ClueGO biological pathway involving proteins in A) white matter, B) grey matter and C) Molecular layer.

### Slide 20
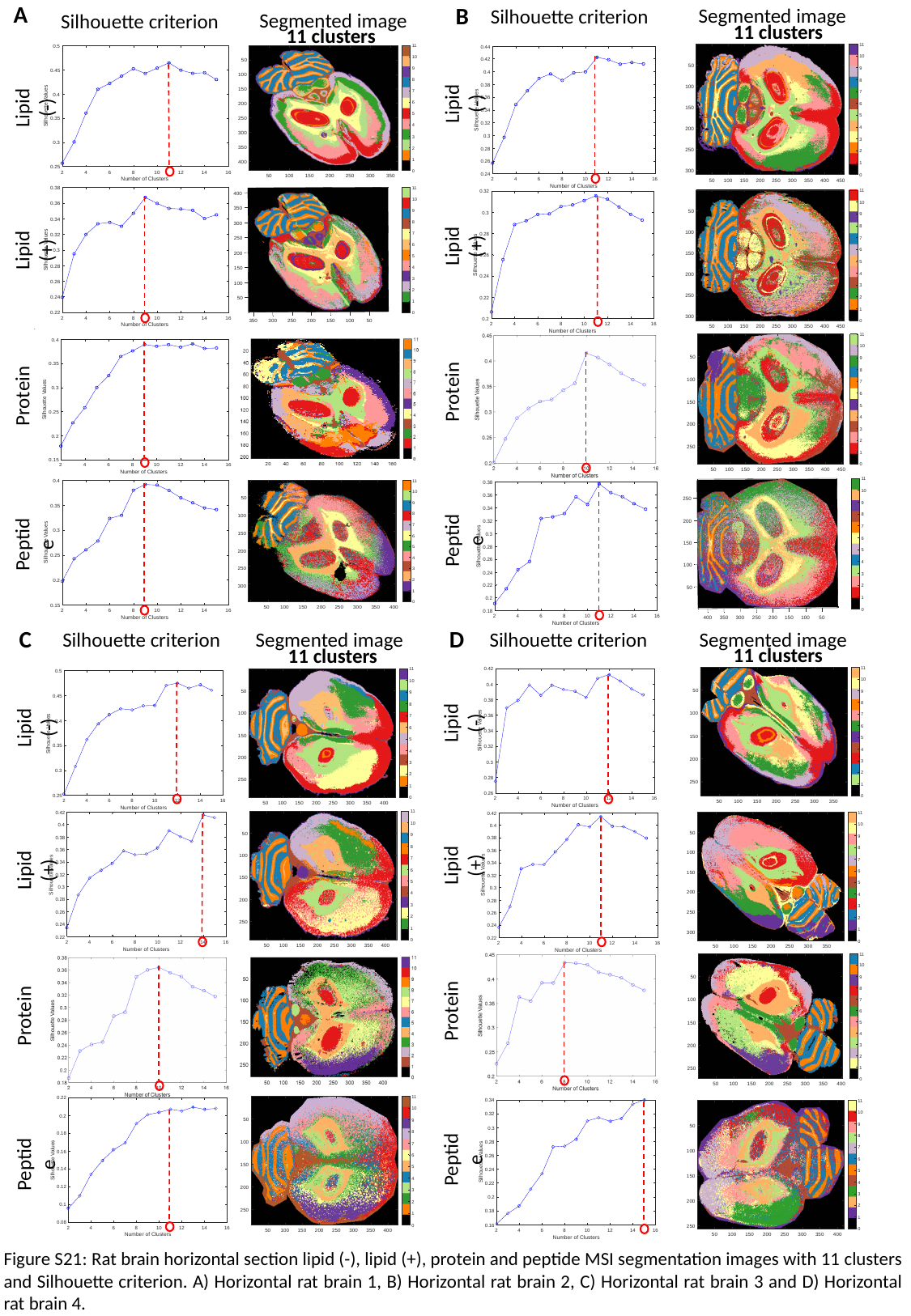

A
B
Segmented image
Silhouette criterion
11 clusters
Lipid (-)
Lipid (+)
Protein
Peptide
Segmented image
Silhouette criterion
11 clusters
Lipid (-)
Lipid (+)
Protein
Peptide
C
D
Silhouette criterion
Segmented image
Silhouette criterion
Segmented image
11 clusters
11 clusters
Lipid (-)
Lipid (-)
Lipid (+)
Lipid (+)
Protein
Protein
Peptide
Peptide
Figure S21: Rat brain horizontal section lipid (-), lipid (+), protein and peptide MSI segmentation images with 11 clusters and Silhouette criterion. A) Horizontal rat brain 1, B) Horizontal rat brain 2, C) Horizontal rat brain 3 and D) Horizontal rat brain 4.

### Slide 21
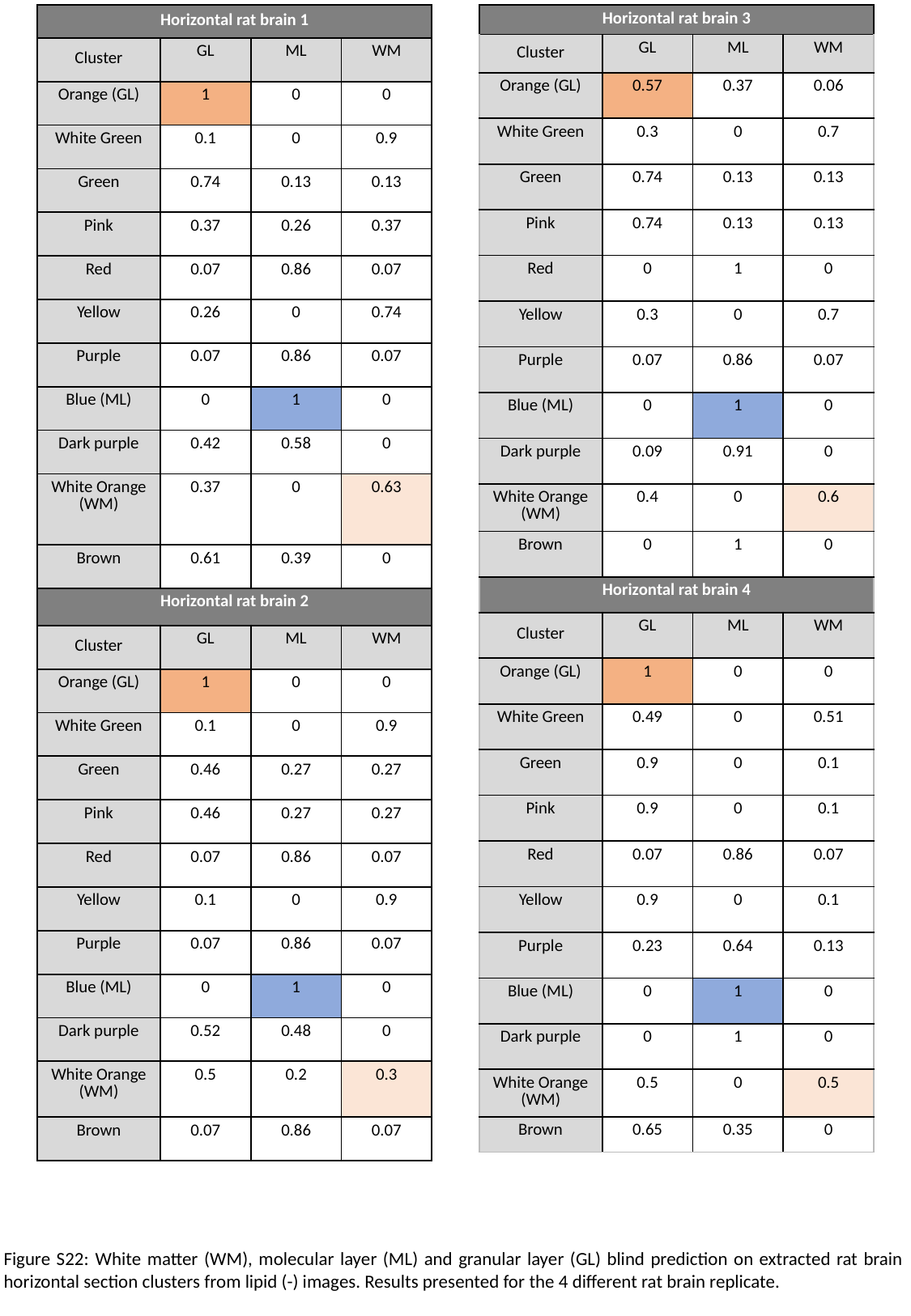

| Horizontal rat brain 1 | | | |
| --- | --- | --- | --- |
| Cluster | GL | ML | WM |
| Orange (GL) | 1 | 0 | 0 |
| White Green | 0.1 | 0 | 0.9 |
| Green | 0.74 | 0.13 | 0.13 |
| Pink | 0.37 | 0.26 | 0.37 |
| Red | 0.07 | 0.86 | 0.07 |
| Yellow | 0.26 | 0 | 0.74 |
| Purple | 0.07 | 0.86 | 0.07 |
| Blue (ML) | 0 | 1 | 0 |
| Dark purple | 0.42 | 0.58 | 0 |
| White Orange (WM) | 0.37 | 0 | 0.63 |
| Brown | 0.61 | 0.39 | 0 |
| Horizontal rat brain 2 | | | |
| Cluster | GL | ML | WM |
| Orange (GL) | 1 | 0 | 0 |
| White Green | 0.1 | 0 | 0.9 |
| Green | 0.46 | 0.27 | 0.27 |
| Pink | 0.46 | 0.27 | 0.27 |
| Red | 0.07 | 0.86 | 0.07 |
| Yellow | 0.1 | 0 | 0.9 |
| Purple | 0.07 | 0.86 | 0.07 |
| Blue (ML) | 0 | 1 | 0 |
| Dark purple | 0.52 | 0.48 | 0 |
| White Orange (WM) | 0.5 | 0.2 | 0.3 |
| Brown | 0.07 | 0.86 | 0.07 |
| Horizontal rat brain 3 | | | |
| --- | --- | --- | --- |
| Cluster | GL | ML | WM |
| Orange (GL) | 0.57 | 0.37 | 0.06 |
| White Green | 0.3 | 0 | 0.7 |
| Green | 0.74 | 0.13 | 0.13 |
| Pink | 0.74 | 0.13 | 0.13 |
| Red | 0 | 1 | 0 |
| Yellow | 0.3 | 0 | 0.7 |
| Purple | 0.07 | 0.86 | 0.07 |
| Blue (ML) | 0 | 1 | 0 |
| Dark purple | 0.09 | 0.91 | 0 |
| White Orange (WM) | 0.4 | 0 | 0.6 |
| Brown | 0 | 1 | 0 |
| Horizontal rat brain 4 | | | |
| Cluster | GL | ML | WM |
| Orange (GL) | 1 | 0 | 0 |
| White Green | 0.49 | 0 | 0.51 |
| Green | 0.9 | 0 | 0.1 |
| Pink | 0.9 | 0 | 0.1 |
| Red | 0.07 | 0.86 | 0.07 |
| Yellow | 0.9 | 0 | 0.1 |
| Purple | 0.23 | 0.64 | 0.13 |
| Blue (ML) | 0 | 1 | 0 |
| Dark purple | 0 | 1 | 0 |
| White Orange (WM) | 0.5 | 0 | 0.5 |
| Brown | 0.65 | 0.35 | 0 |
Figure S22: White matter (WM), molecular layer (ML) and granular layer (GL) blind prediction on extracted rat brain horizontal section clusters from lipid (-) images. Results presented for the 4 different rat brain replicate.

### Slide 22
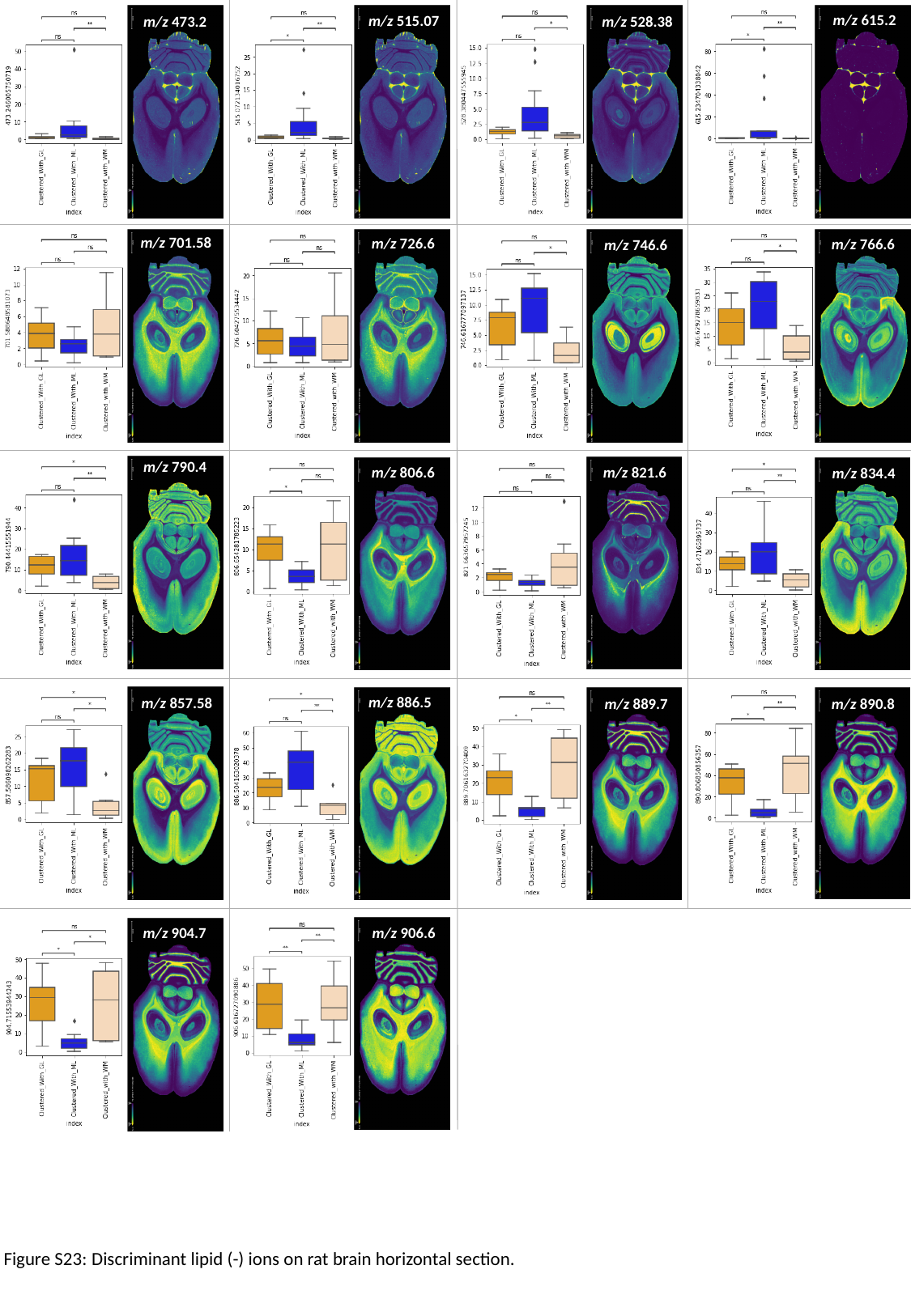

m/z 615.2
m/z 515.07
m/z 528.38
m/z 473.2
m/z 701.58
m/z 726.6
m/z 766.6
m/z 746.6
m/z 790.4
m/z 806.6
m/z 821.6
m/z 834.4
m/z 886.5
m/z 857.58
m/z 889.7
m/z 890.8
m/z 904.7
m/z 906.6
Figure S23: Discriminant lipid (-) ions on rat brain horizontal section.

### Slide 23
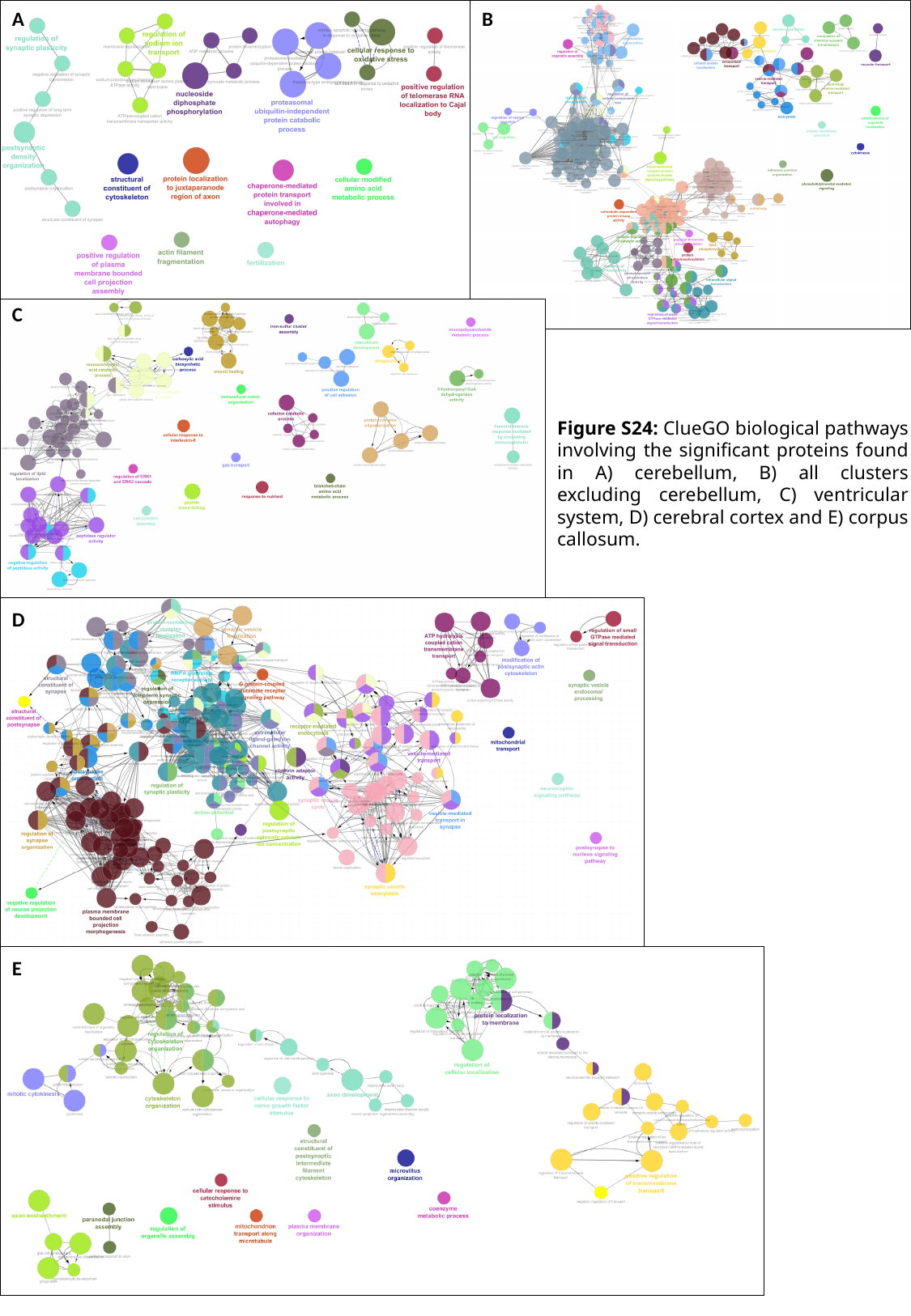

A
B
C
Figure S24: ClueGO biological pathways involving the significant proteins found in A) cerebellum, B) all clusters excluding cerebellum, C) ventricular system, D) cerebral cortex and E) corpus callosum.
D
E

### Slide 24
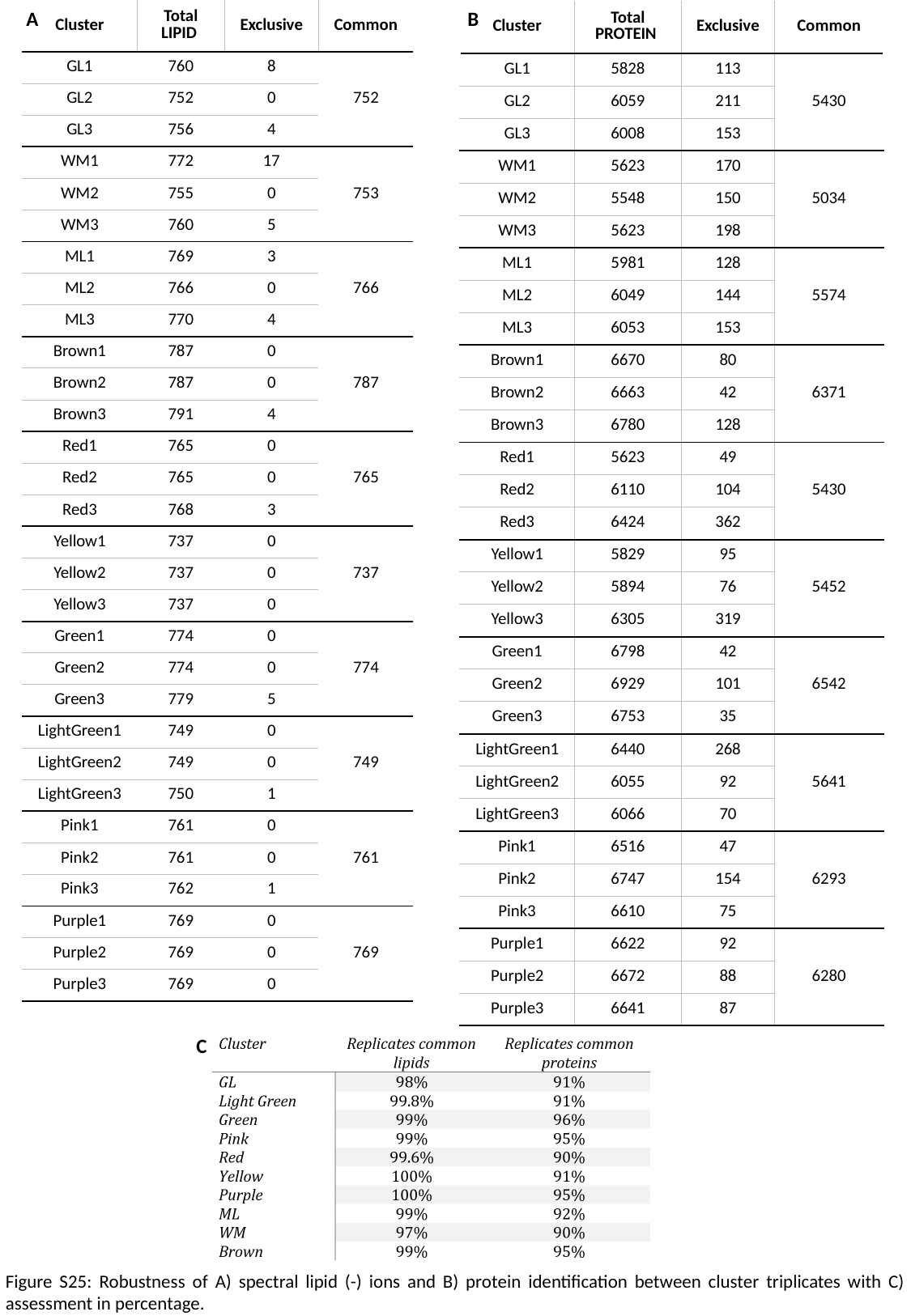

A
| Cluster | Total LIPID | Exclusive | Common |
| --- | --- | --- | --- |
| GL1 | 760 | 8 | 752 |
| GL2 | 752 | 0 | |
| GL3 | 756 | 4 | |
| WM1 | 772 | 17 | 753 |
| WM2 | 755 | 0 | |
| WM3 | 760 | 5 | |
| ML1 | 769 | 3 | 766 |
| ML2 | 766 | 0 | |
| ML3 | 770 | 4 | |
| Brown1 | 787 | 0 | 787 |
| Brown2 | 787 | 0 | |
| Brown3 | 791 | 4 | |
| Red1 | 765 | 0 | 765 |
| Red2 | 765 | 0 | |
| Red3 | 768 | 3 | |
| Yellow1 | 737 | 0 | 737 |
| Yellow2 | 737 | 0 | |
| Yellow3 | 737 | 0 | |
| Green1 | 774 | 0 | 774 |
| Green2 | 774 | 0 | |
| Green3 | 779 | 5 | |
| LightGreen1 | 749 | 0 | 749 |
| LightGreen2 | 749 | 0 | |
| LightGreen3 | 750 | 1 | |
| Pink1 | 761 | 0 | 761 |
| Pink2 | 761 | 0 | |
| Pink3 | 762 | 1 | |
| Purple1 | 769 | 0 | 769 |
| Purple2 | 769 | 0 | |
| Purple3 | 769 | 0 | |
B
| Cluster | Total PROTEIN | Exclusive | Common |
| --- | --- | --- | --- |
| GL1 | 5828 | 113 | 5430 |
| GL2 | 6059 | 211 | |
| GL3 | 6008 | 153 | |
| WM1 | 5623 | 170 | 5034 |
| WM2 | 5548 | 150 | |
| WM3 | 5623 | 198 | |
| ML1 | 5981 | 128 | 5574 |
| ML2 | 6049 | 144 | |
| ML3 | 6053 | 153 | |
| Brown1 | 6670 | 80 | 6371 |
| Brown2 | 6663 | 42 | |
| Brown3 | 6780 | 128 | |
| Red1 | 5623 | 49 | 5430 |
| Red2 | 6110 | 104 | |
| Red3 | 6424 | 362 | |
| Yellow1 | 5829 | 95 | 5452 |
| Yellow2 | 5894 | 76 | |
| Yellow3 | 6305 | 319 | |
| Green1 | 6798 | 42 | 6542 |
| Green2 | 6929 | 101 | |
| Green3 | 6753 | 35 | |
| LightGreen1 | 6440 | 268 | 5641 |
| LightGreen2 | 6055 | 92 | |
| LightGreen3 | 6066 | 70 | |
| Pink1 | 6516 | 47 | 6293 |
| Pink2 | 6747 | 154 | |
| Pink3 | 6610 | 75 | |
| Purple1 | 6622 | 92 | 6280 |
| Purple2 | 6672 | 88 | |
| Purple3 | 6641 | 87 | |
C
Figure S25: Robustness of A) spectral lipid (-) ions and B) protein identification between cluster triplicates with C) assessment in percentage.

### Slide 25
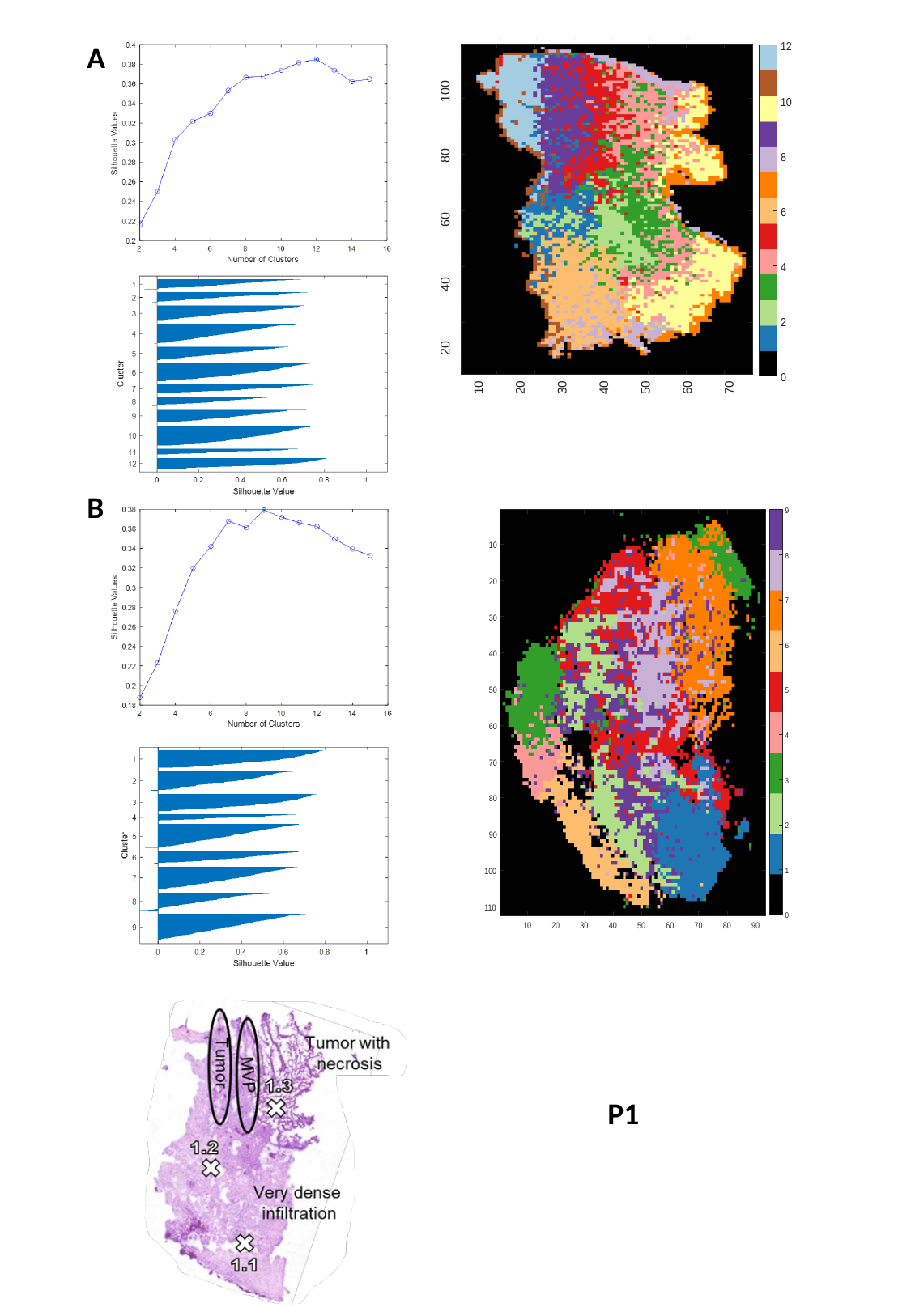

A
B
P1

### Slide 26
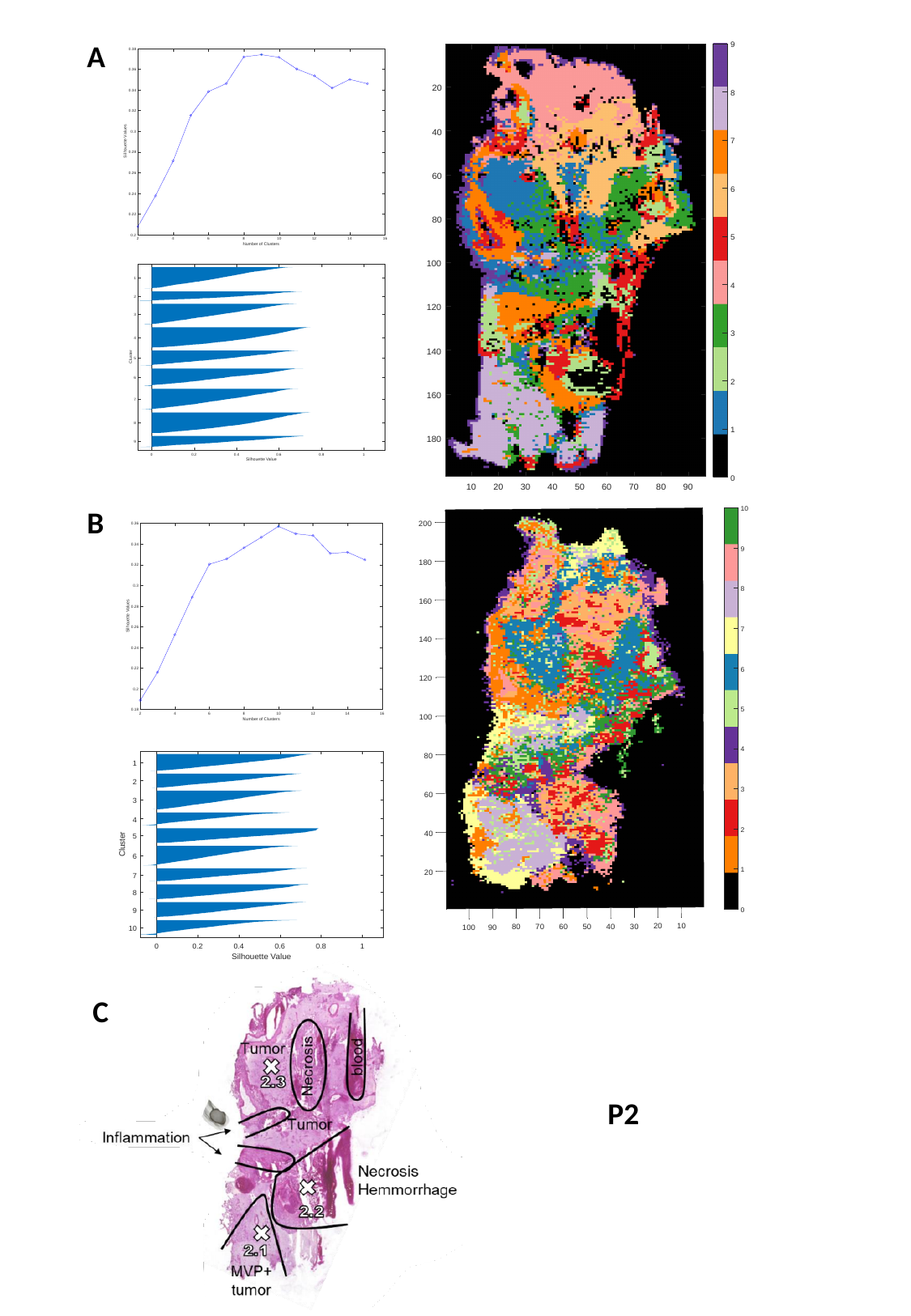

A
B
C
P2

### Slide 27
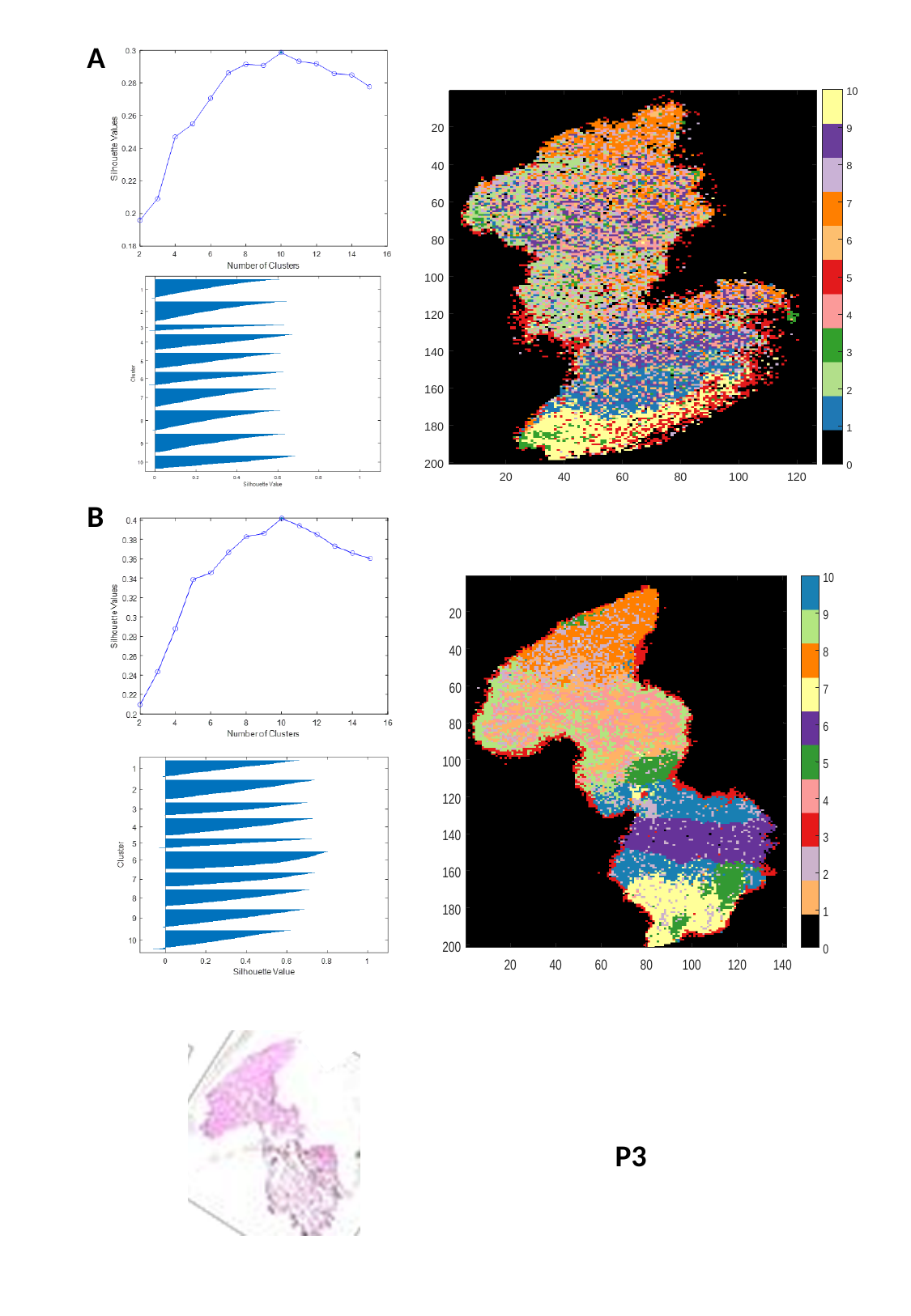

A
B
P3

### Slide 28
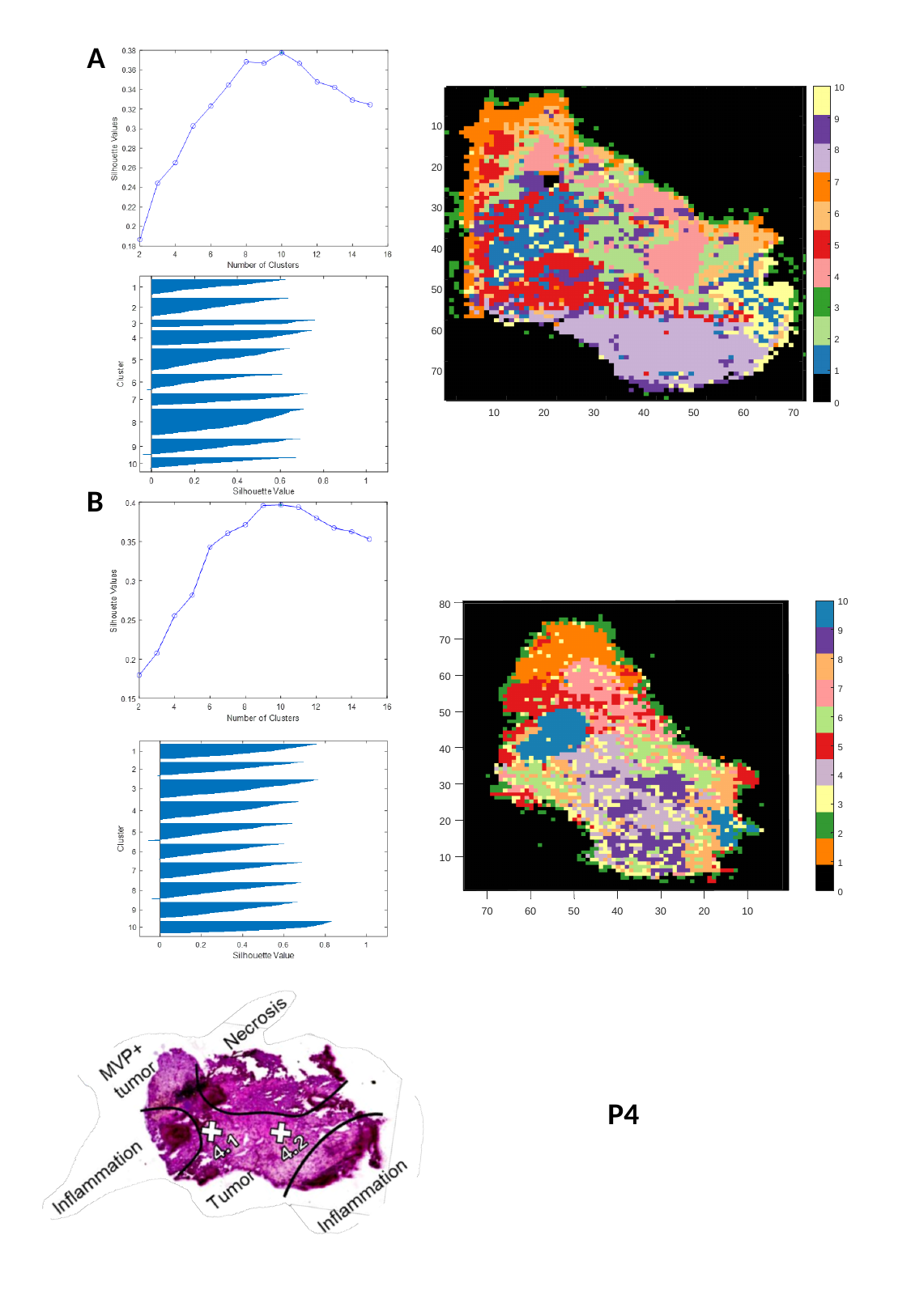

A
B
P4

### Slide 29
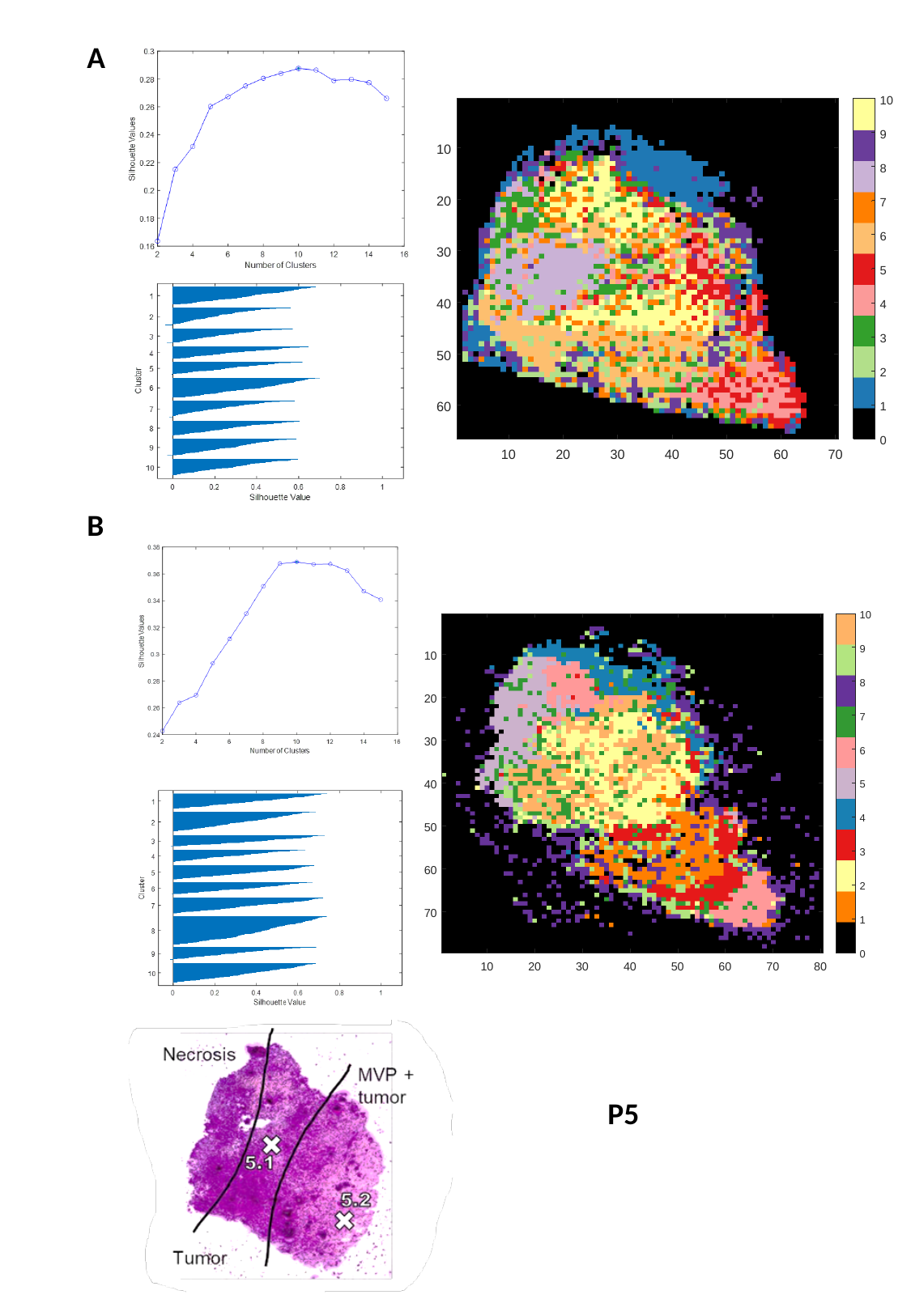

A
B
P5

### Slide 30
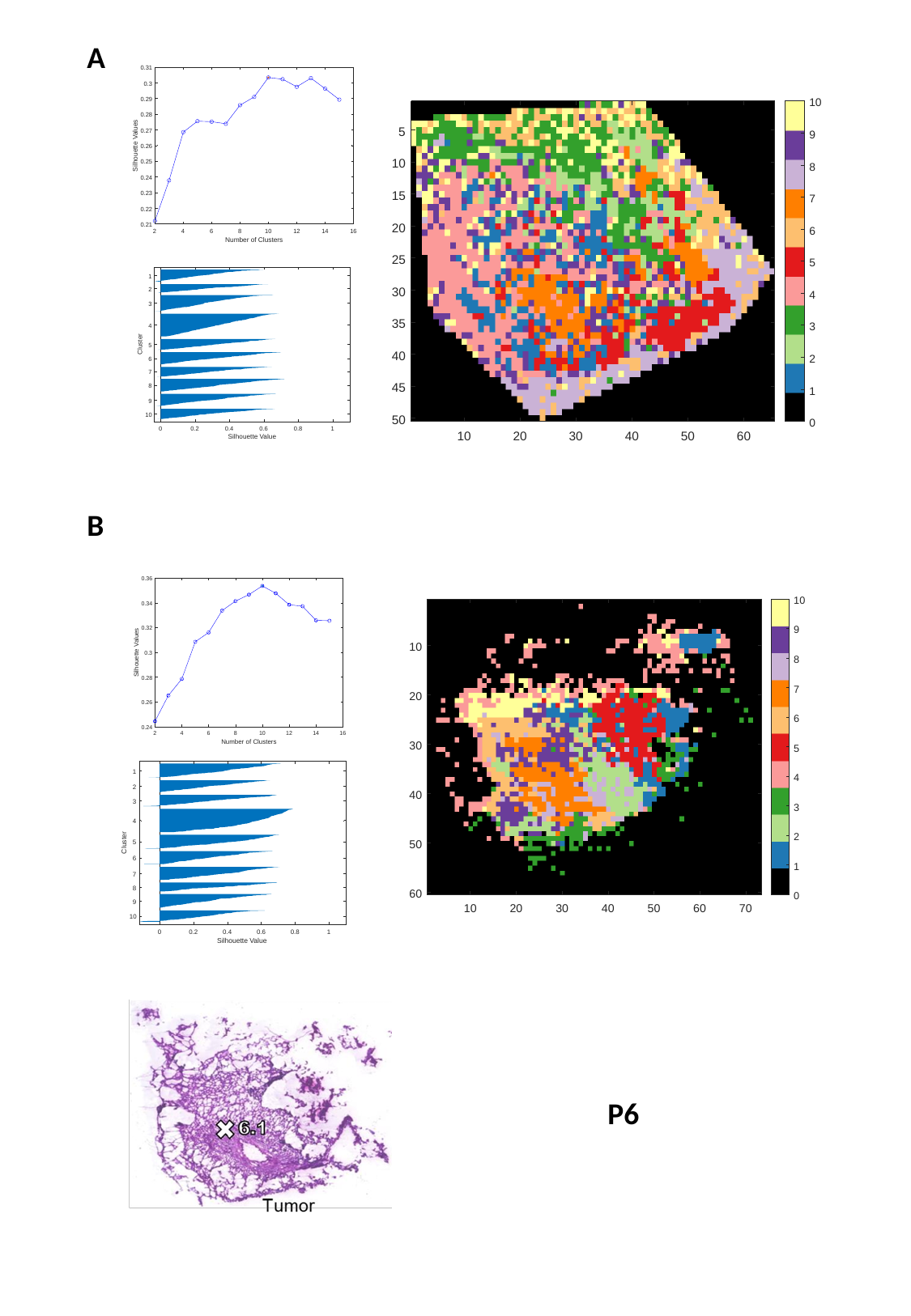

A
B
P6

### Slide 31
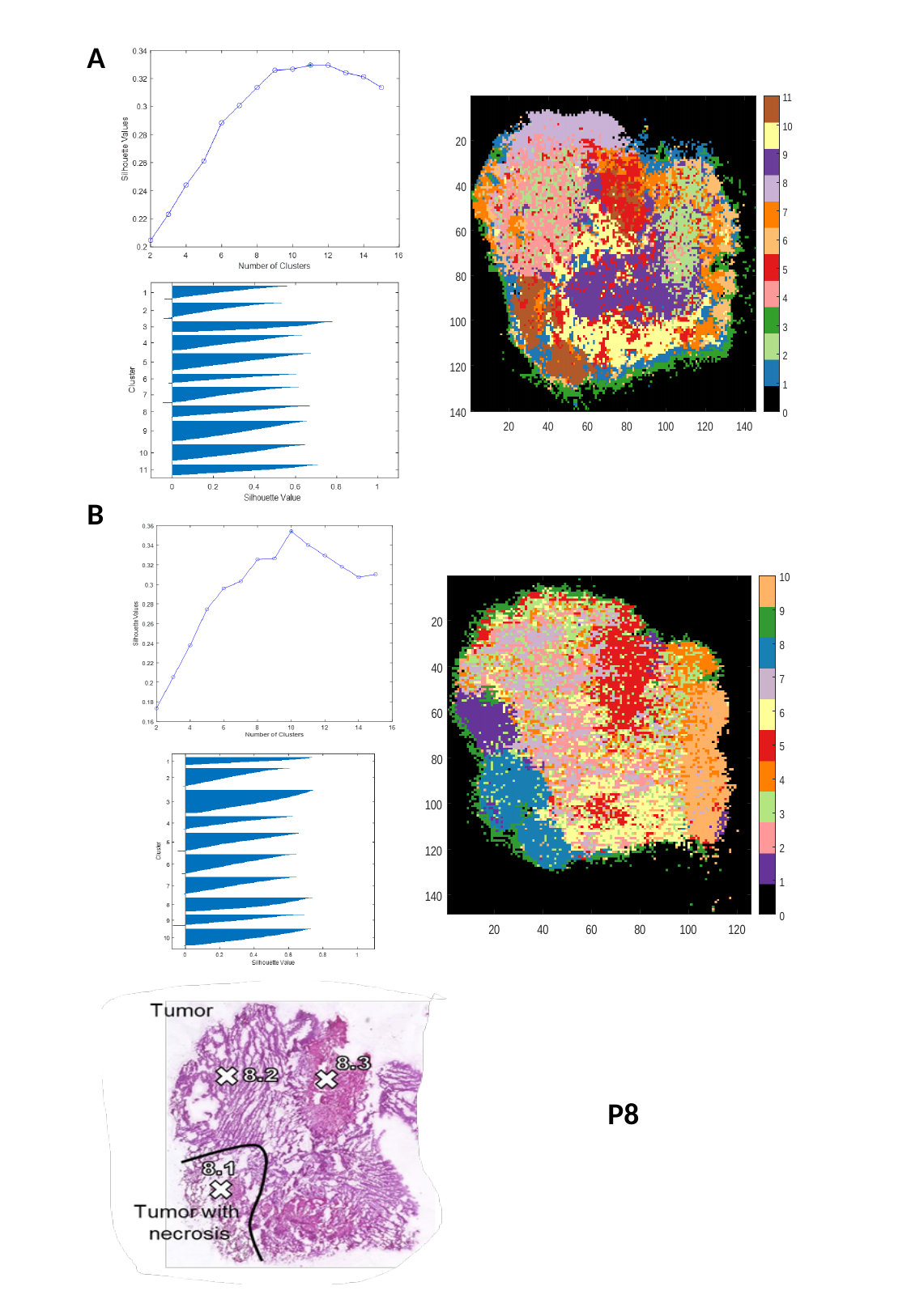

A
B
P8

### Slide 32
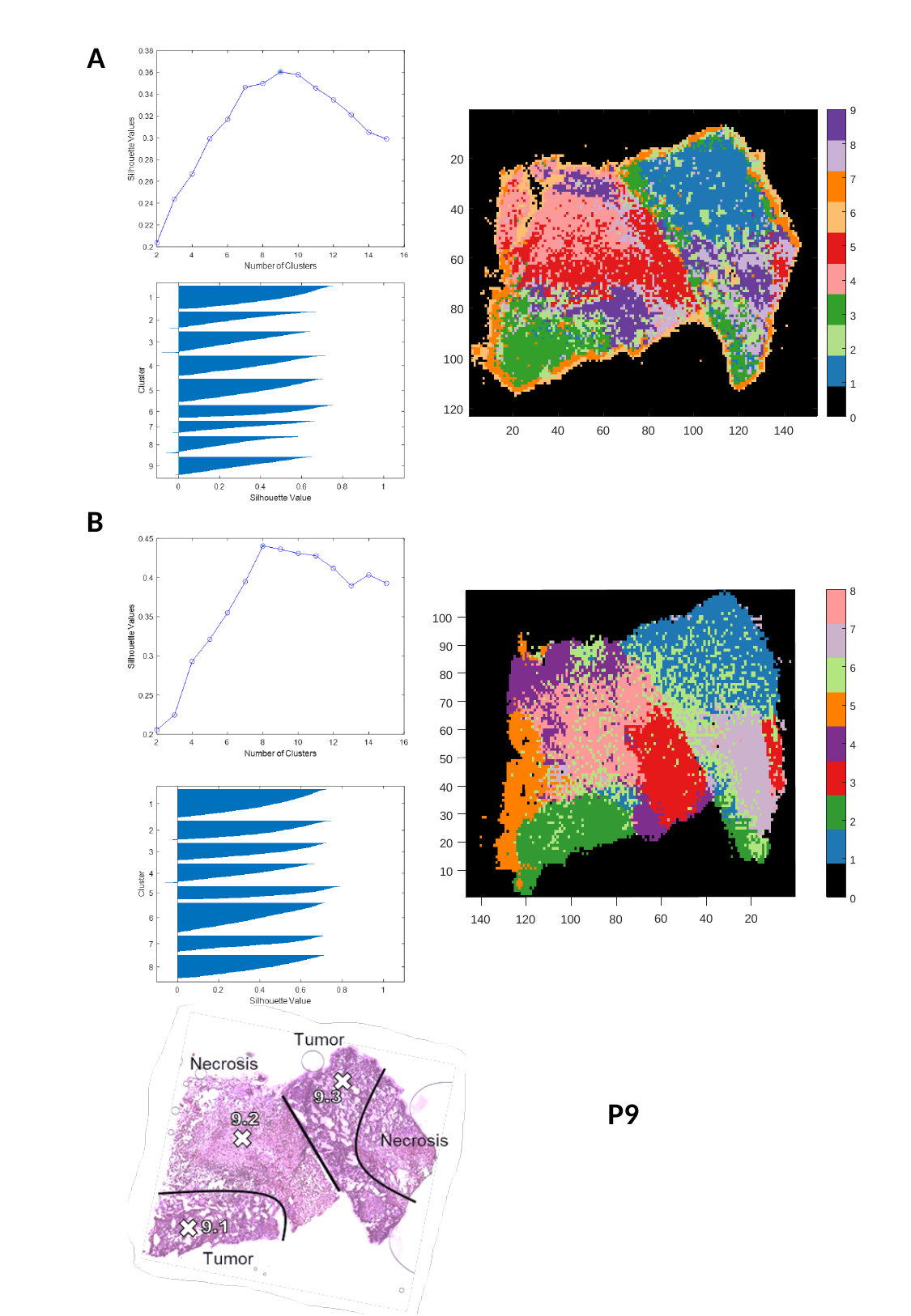

A
B
P9

### Slide 33
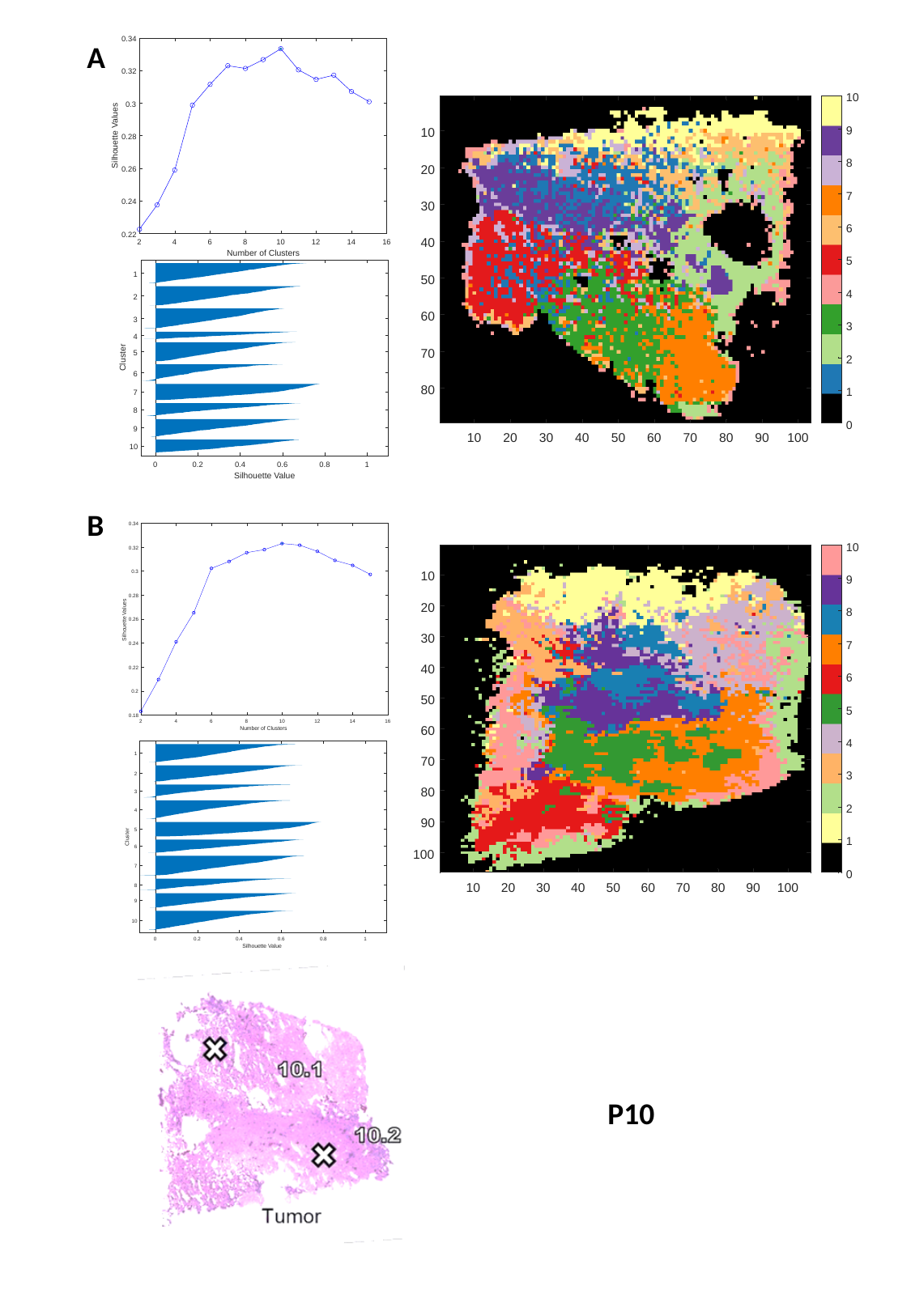

A
B
P10

### Slide 34
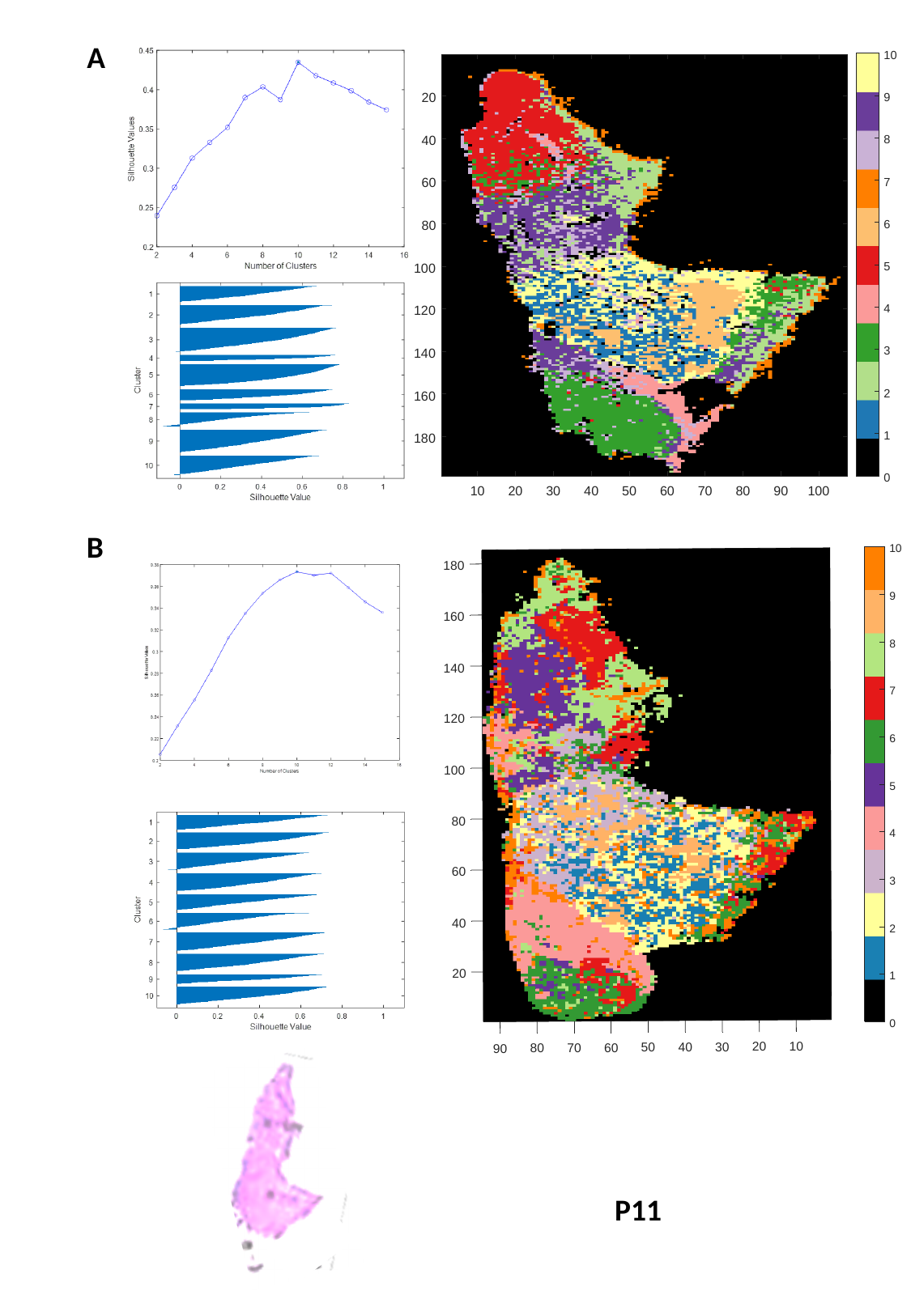

A
B
P11

### Slide 35
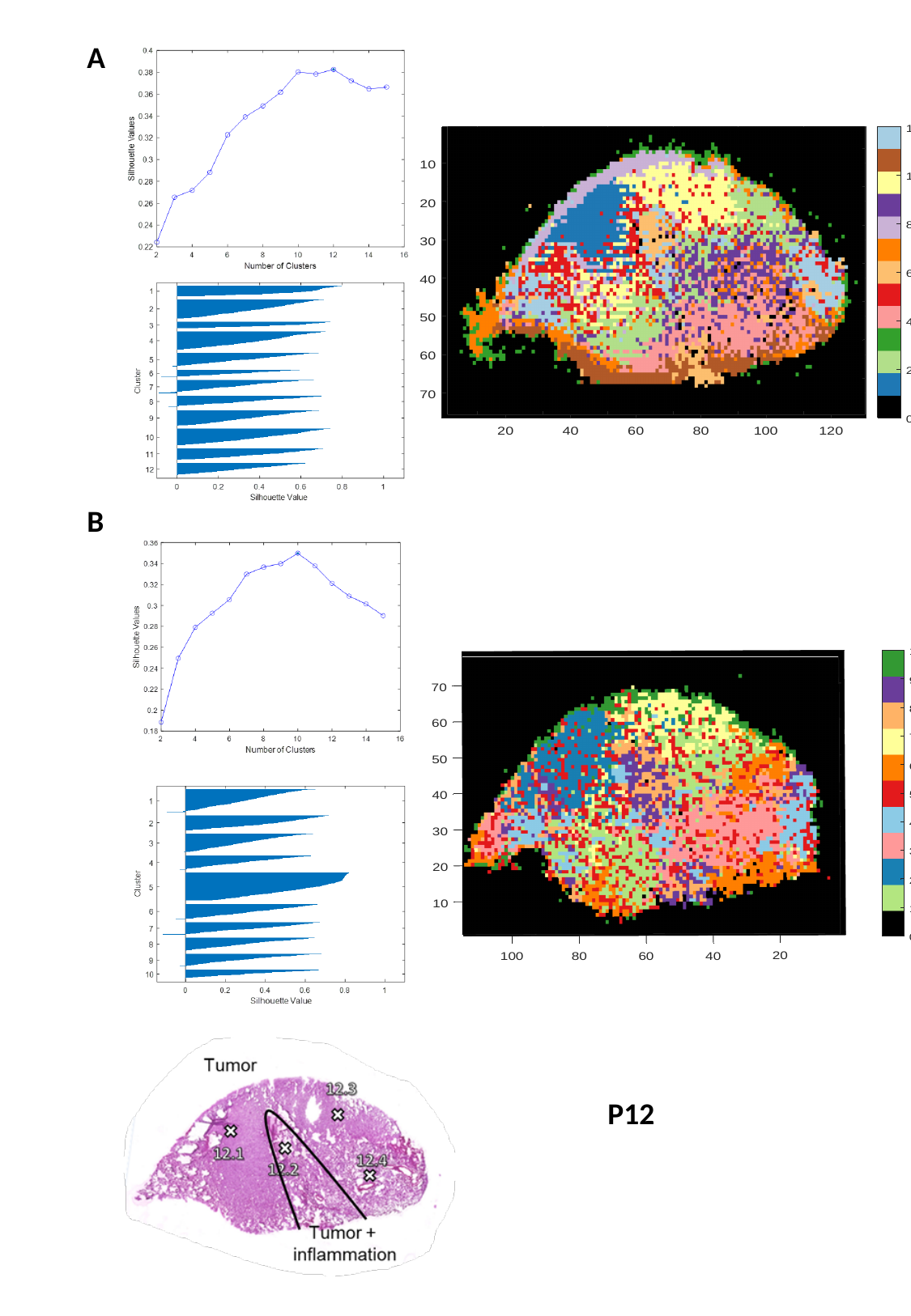

A
B
P12

### Slide 36
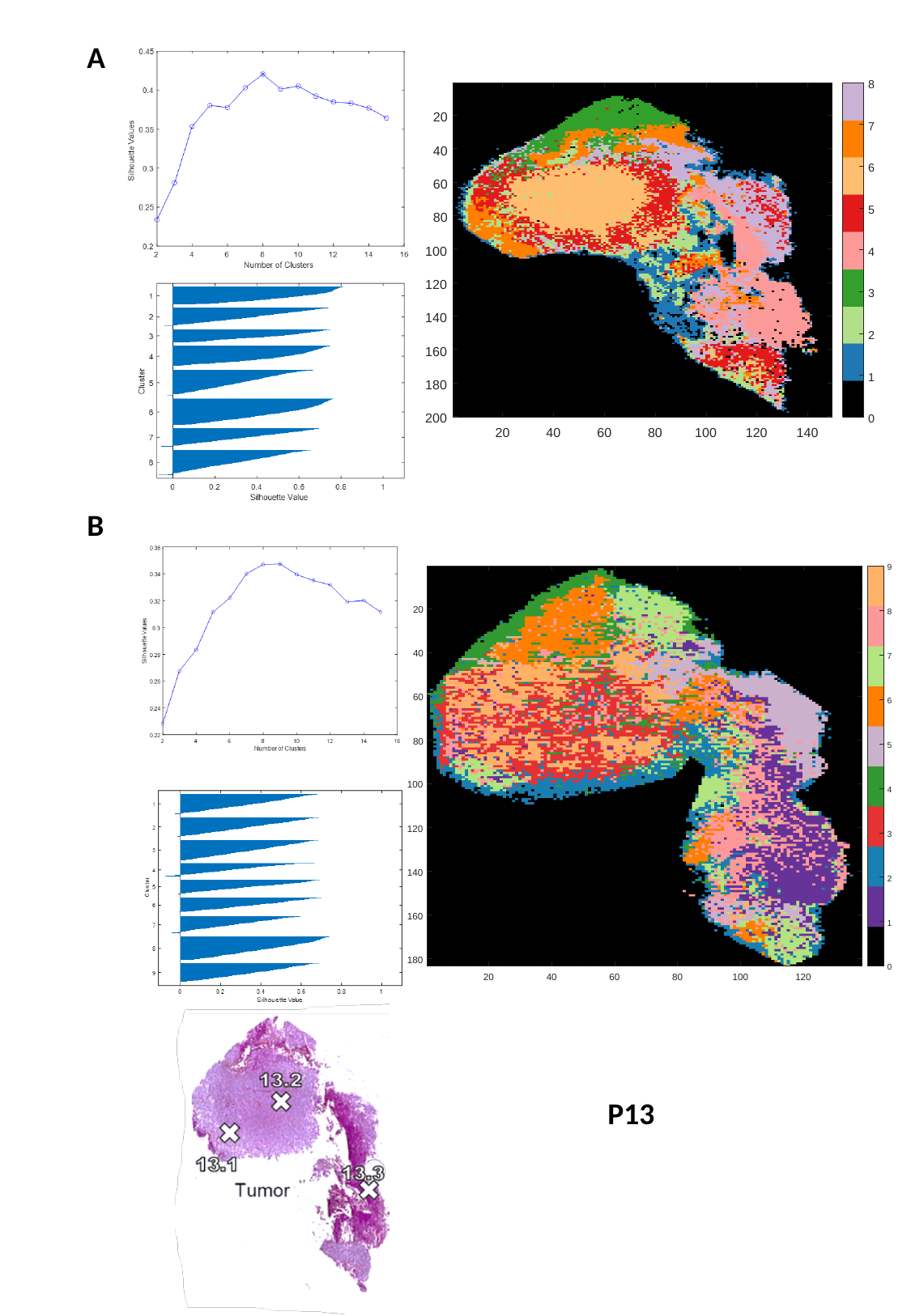

A
B
P13

### Slide 37
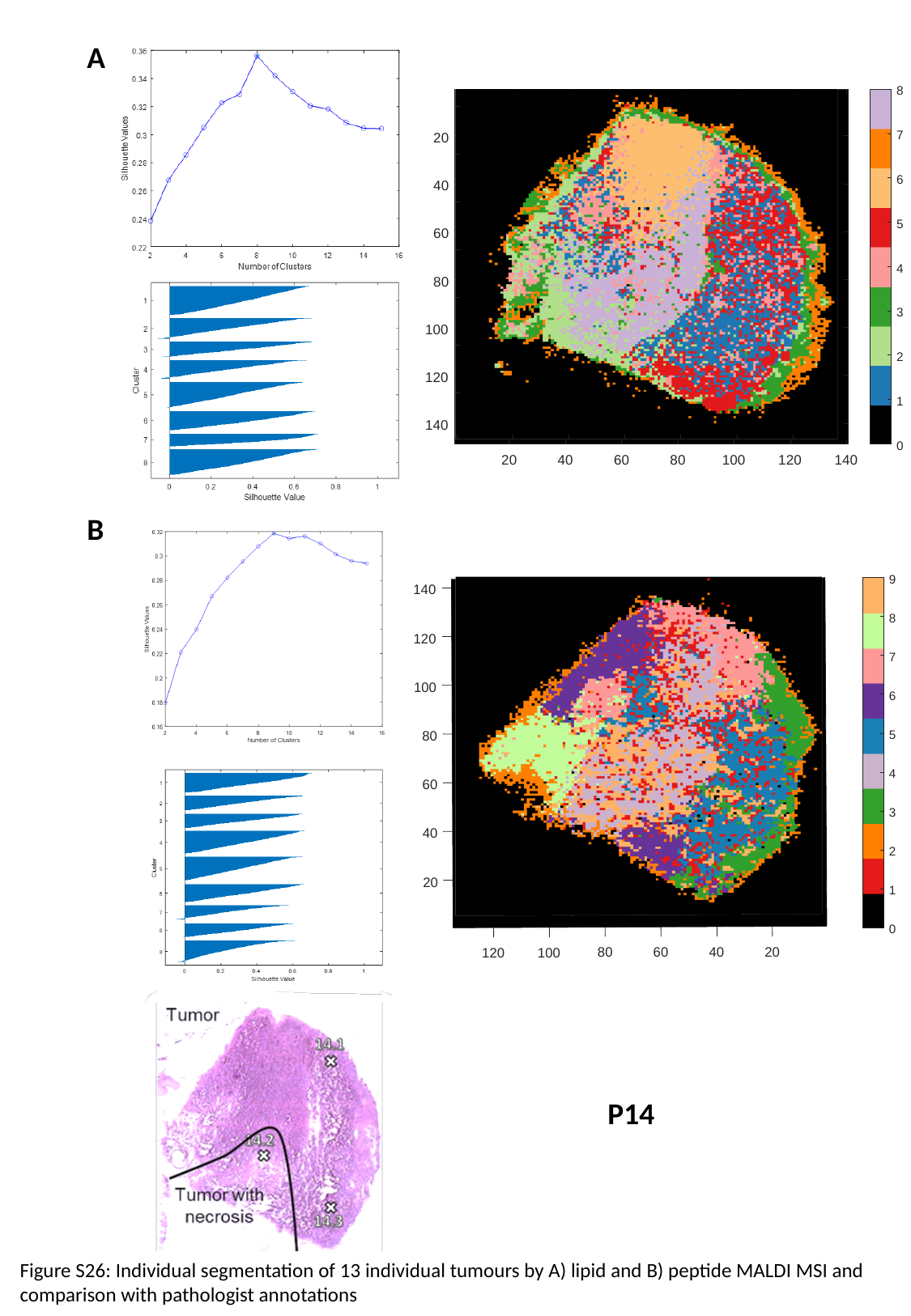

A
B
P14
Figure S26: Individual segmentation of 13 individual tumours by A) lipid and B) peptide MALDI MSI and comparison with pathologist annotations

### Slide 38
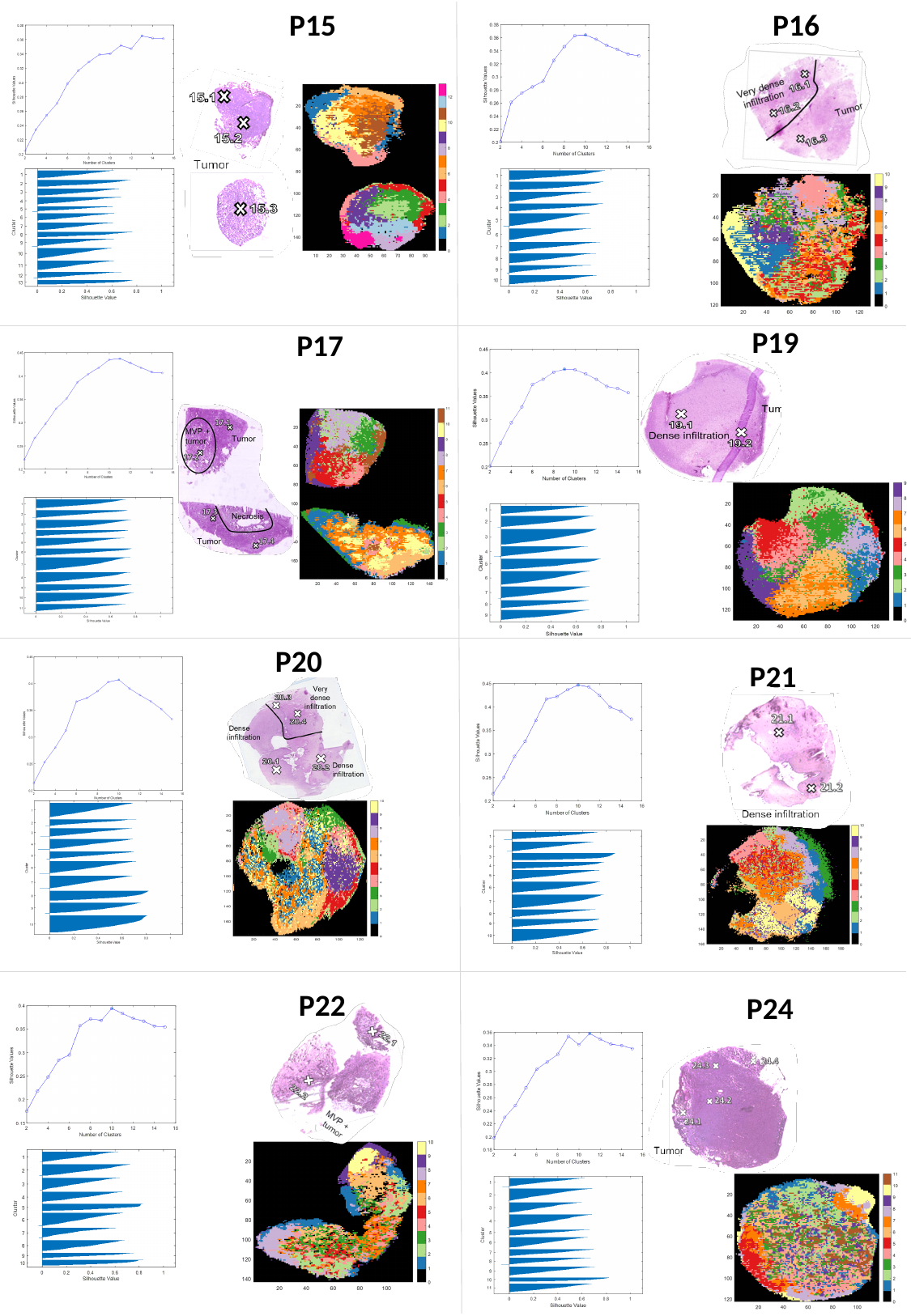

P15
P16
P19
P17
P20
P21
P22
P24

### Slide 39
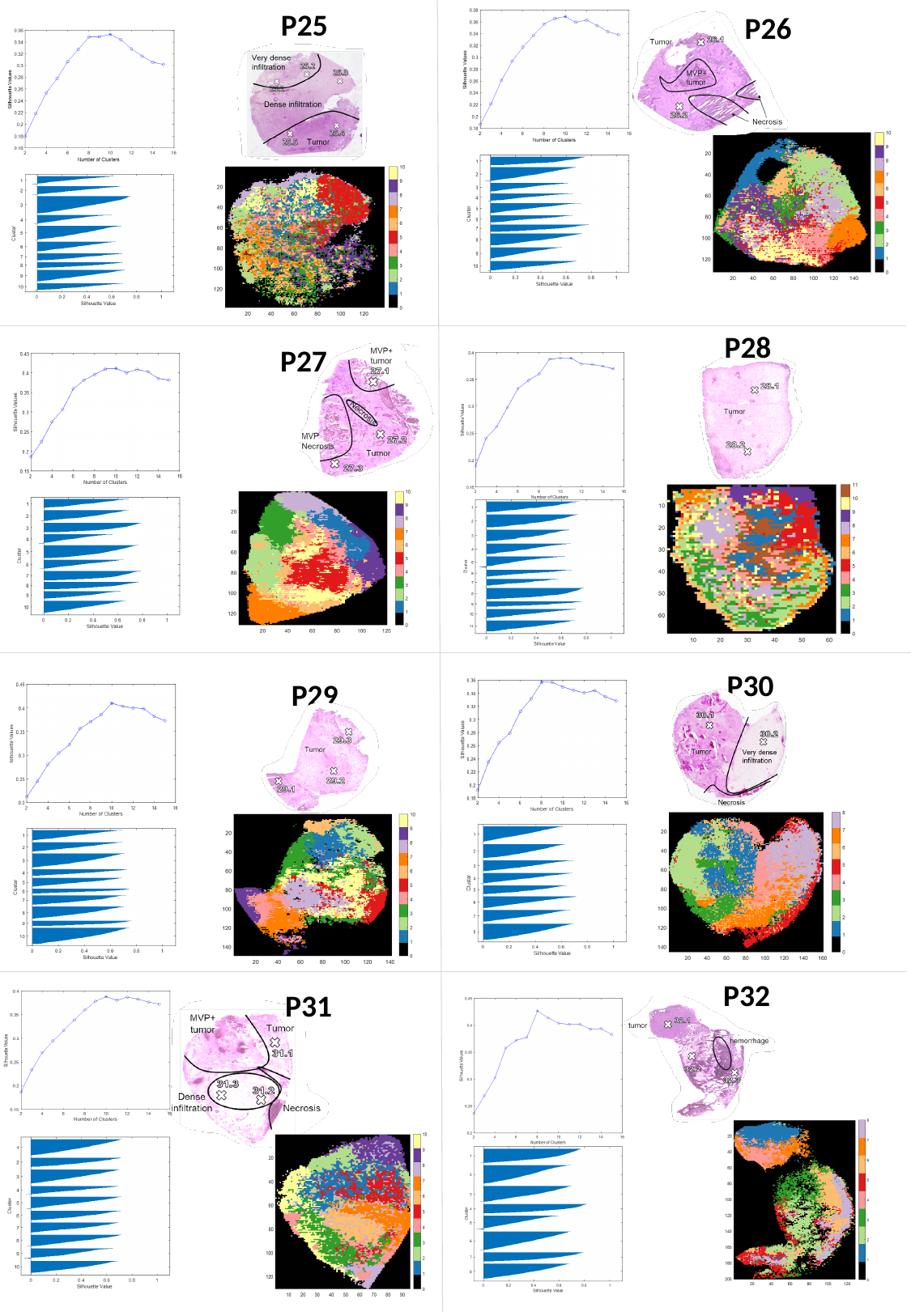

P25
P26
P28
P27
P30
P29
P32
P31

### Slide 40
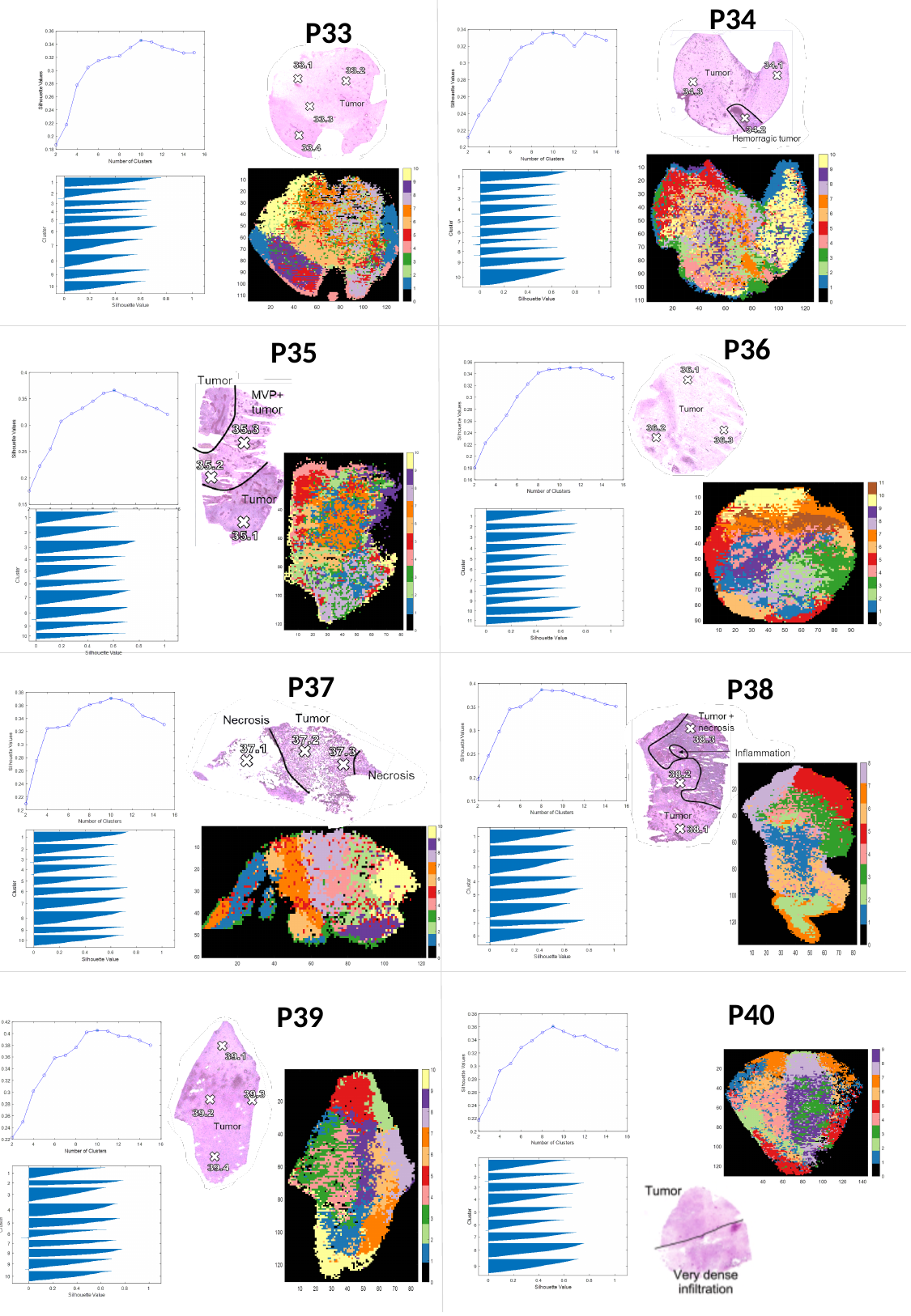

P34
P33
P36
P35
P37
P38
P40
P39

### Slide 41
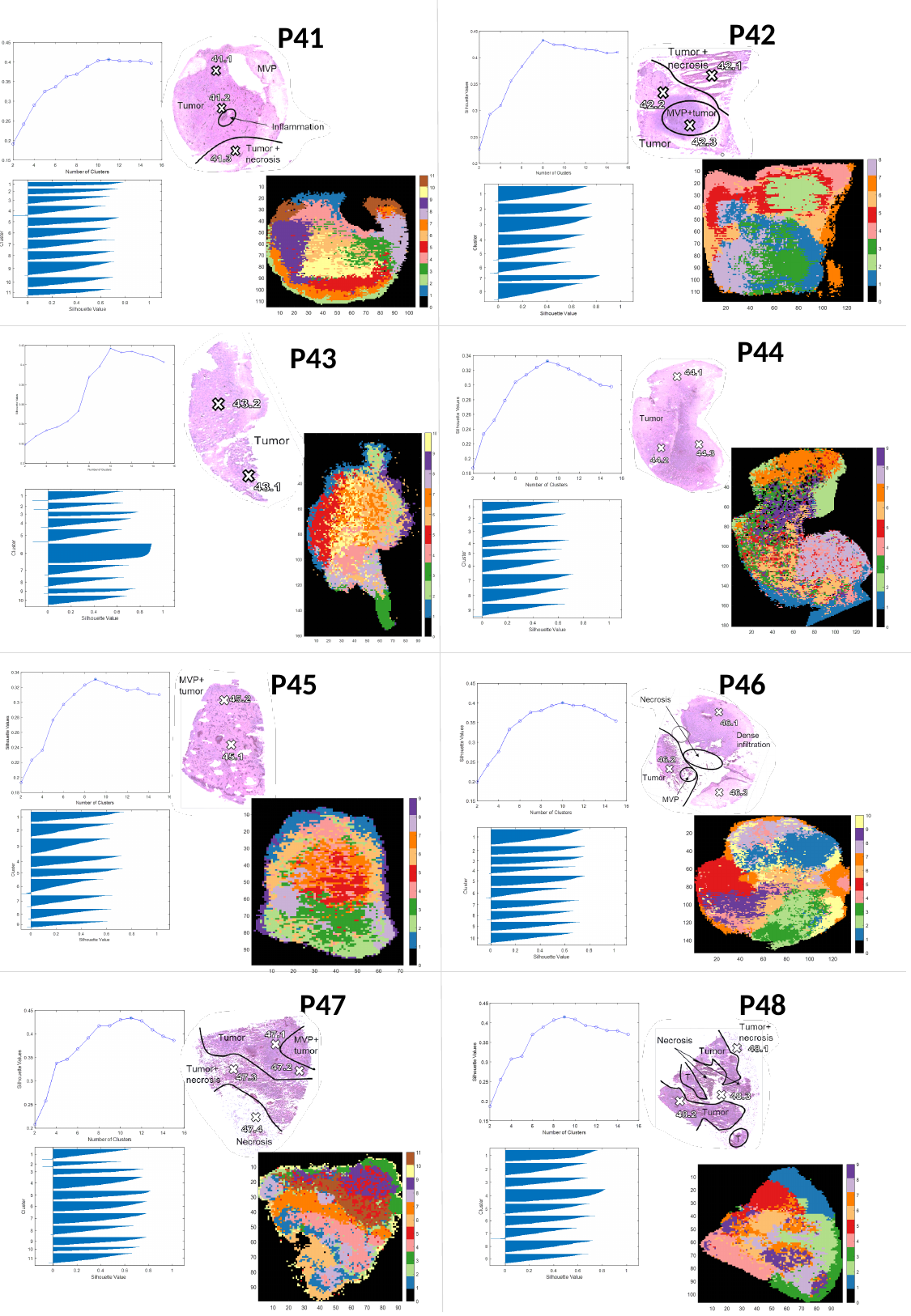

P42
P41
P44
P43
P45
P46
P48
P47
peptide MSI

### Slide 42
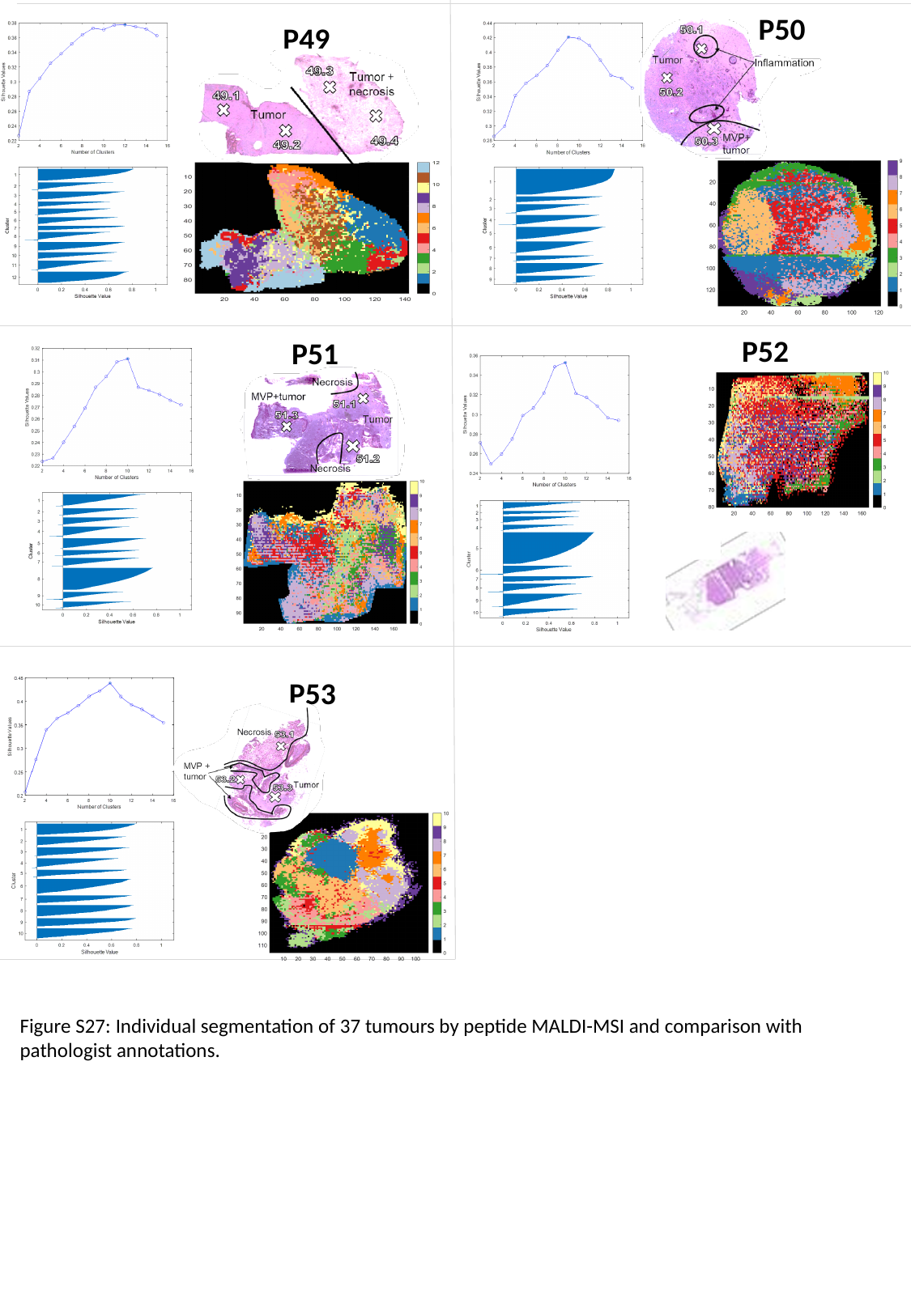

P50
P49
P52
P51
P53
Figure S27: Individual segmentation of 37 tumours by peptide MALDI-MSI and comparison with pathologist annotations.

### Slide 43
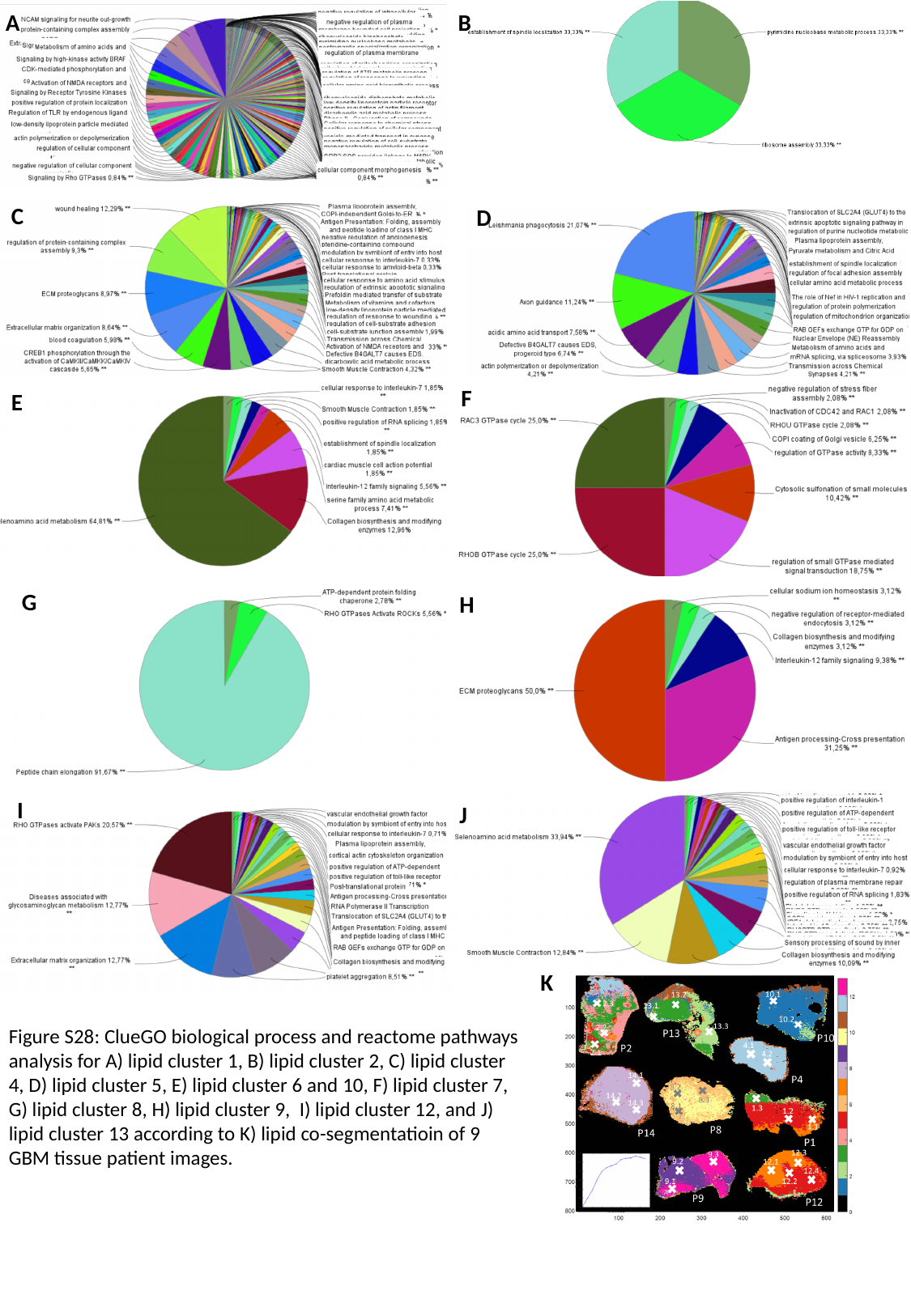

A
B
C
D
F
E
G
H
I
J
K
Figure S28: ClueGO biological process and reactome pathways analysis for A) lipid cluster 1, B) lipid cluster 2, C) lipid cluster 4, D) lipid cluster 5, E) lipid cluster 6 and 10, F) lipid cluster 7, G) lipid cluster 8, H) lipid cluster 9, I) lipid cluster 12, and J) lipid cluster 13 according to K) lipid co-segmentatioin of 9 GBM tissue patient images.

### Slide 44
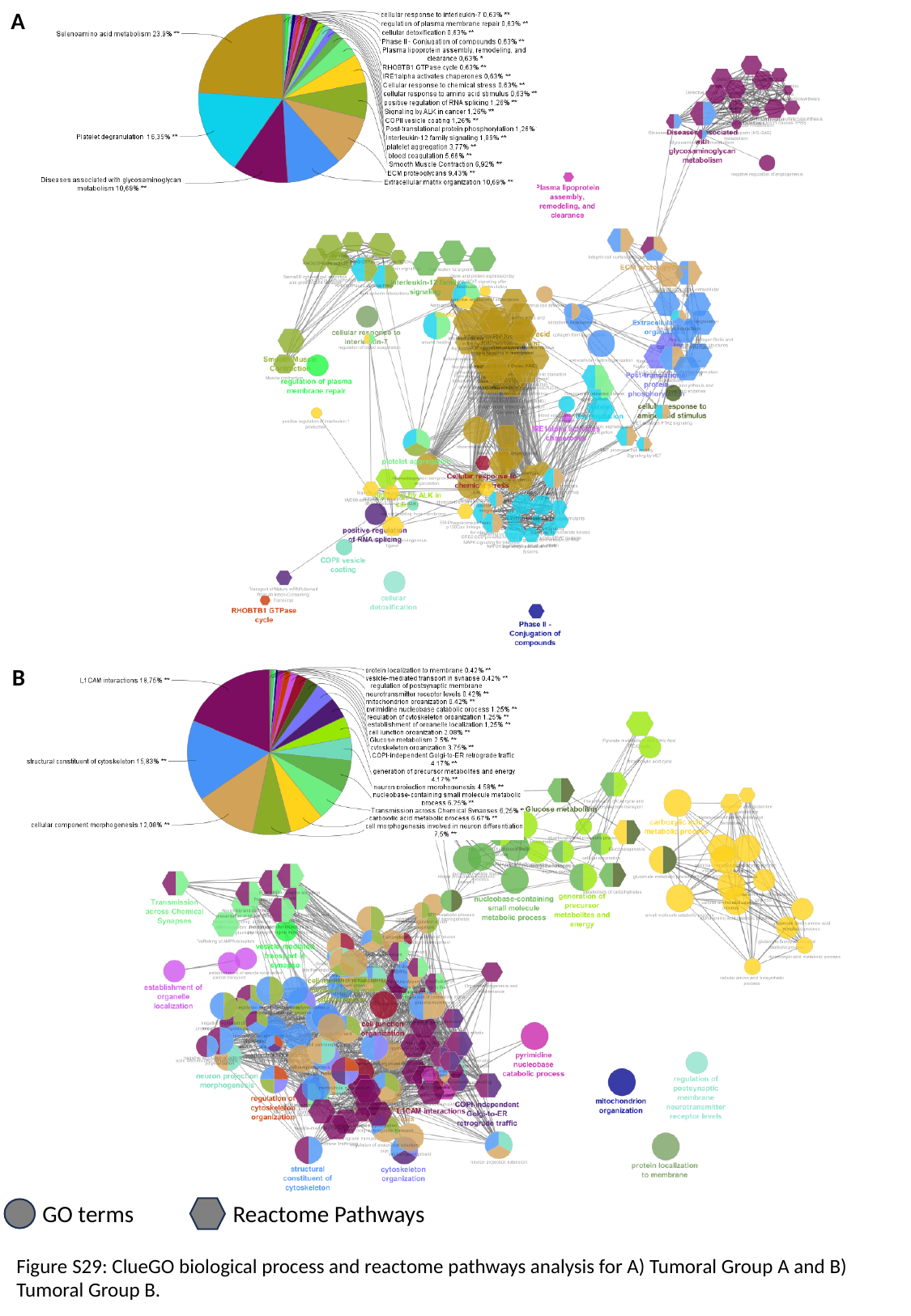

A
B
GO terms
Reactome Pathways
Figure S29: ClueGO biological process and reactome pathways analysis for A) Tumoral Group A and B) Tumoral Group B.

### Slide 45
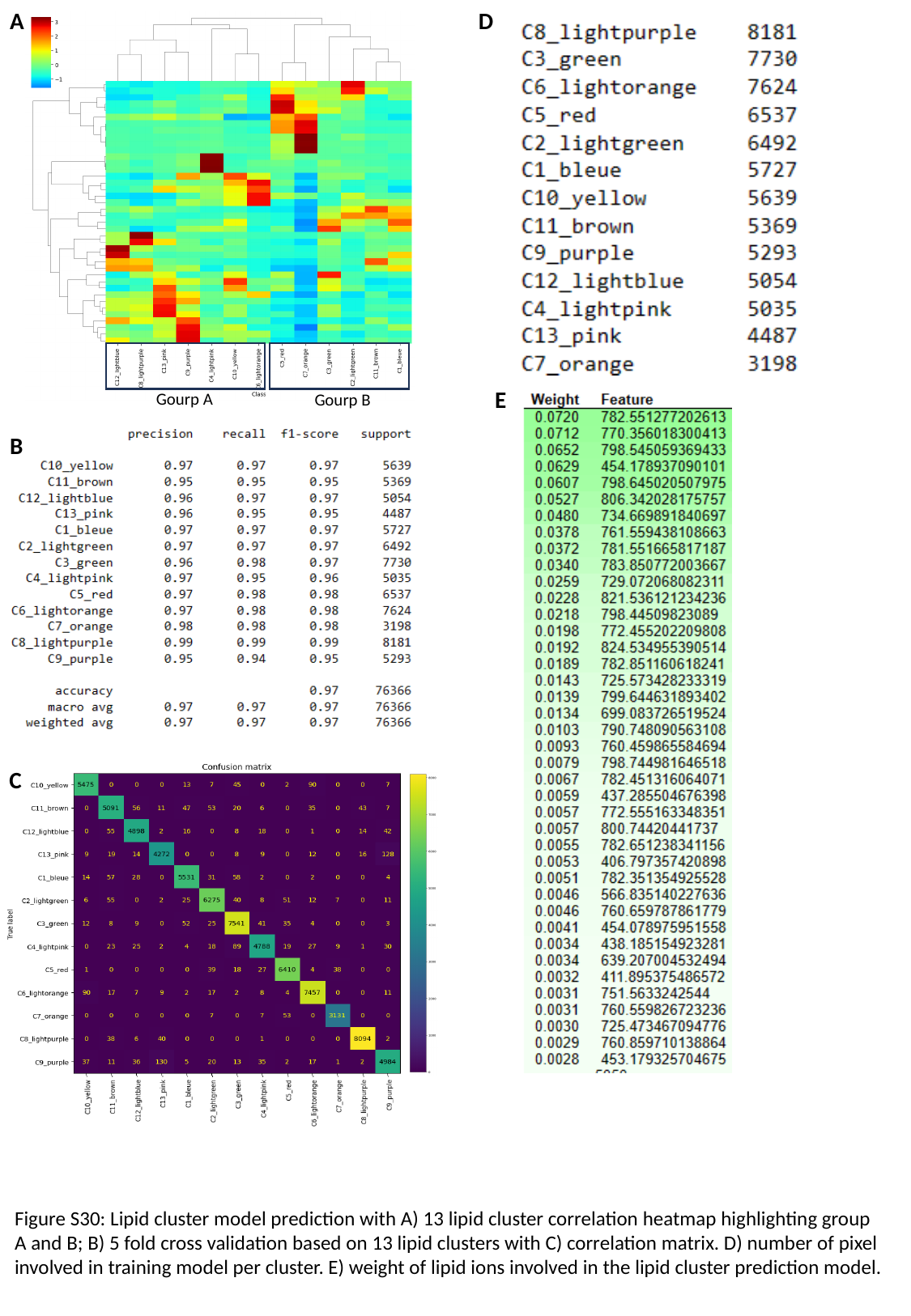

A
Gourp A
Gourp B
D
E
B
C
Figure S30: Lipid cluster model prediction with A) 13 lipid cluster correlation heatmap highlighting group A and B; B) 5 fold cross validation based on 13 lipid clusters with C) correlation matrix. D) number of pixel involved in training model per cluster. E) weight of lipid ions involved in the lipid cluster prediction model.

### Slide 46
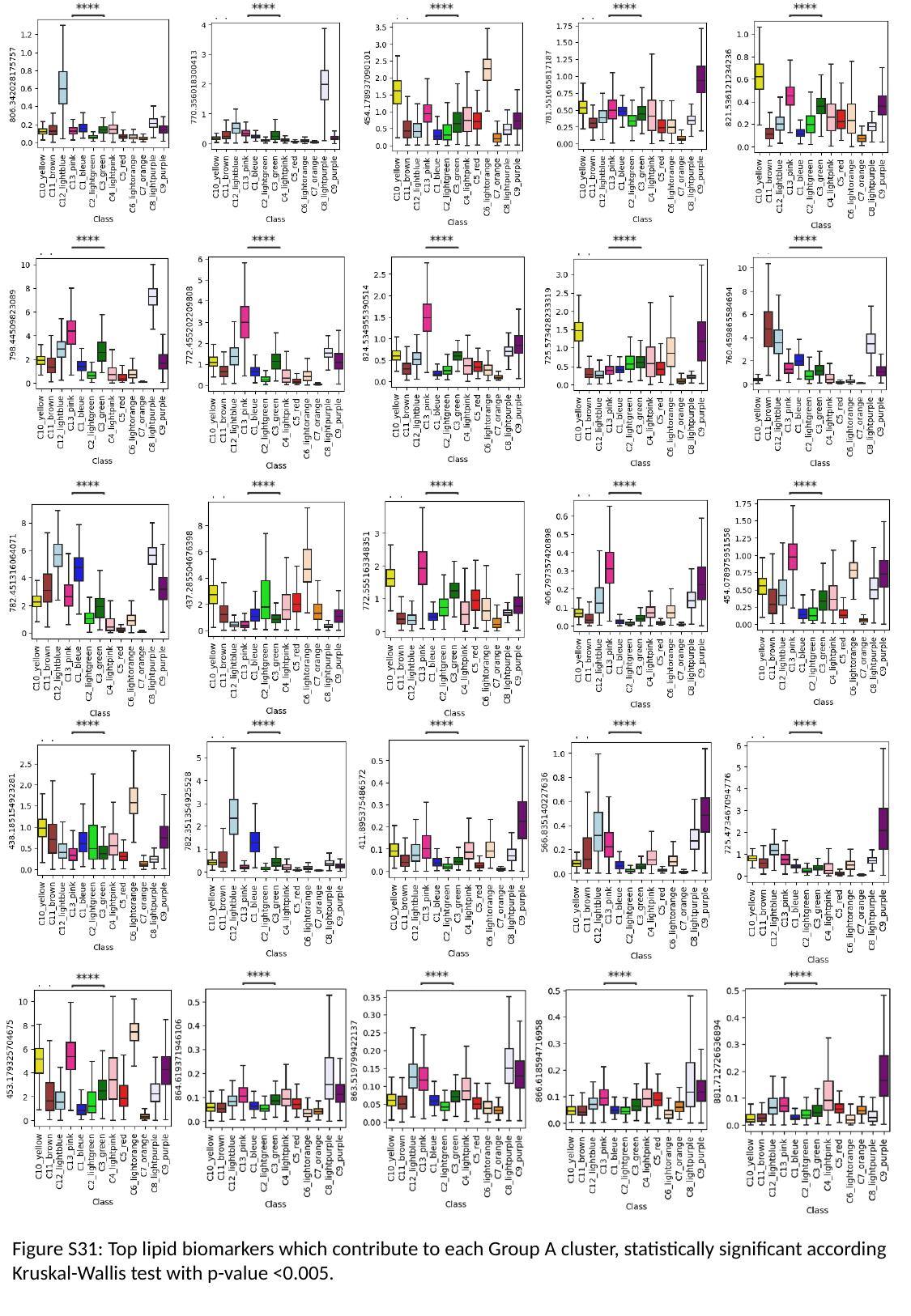

Figure S31: Top lipid biomarkers which contribute to each Group A cluster, statistically significant according Kruskal-Wallis test with p-value <0.005.

### Slide 47
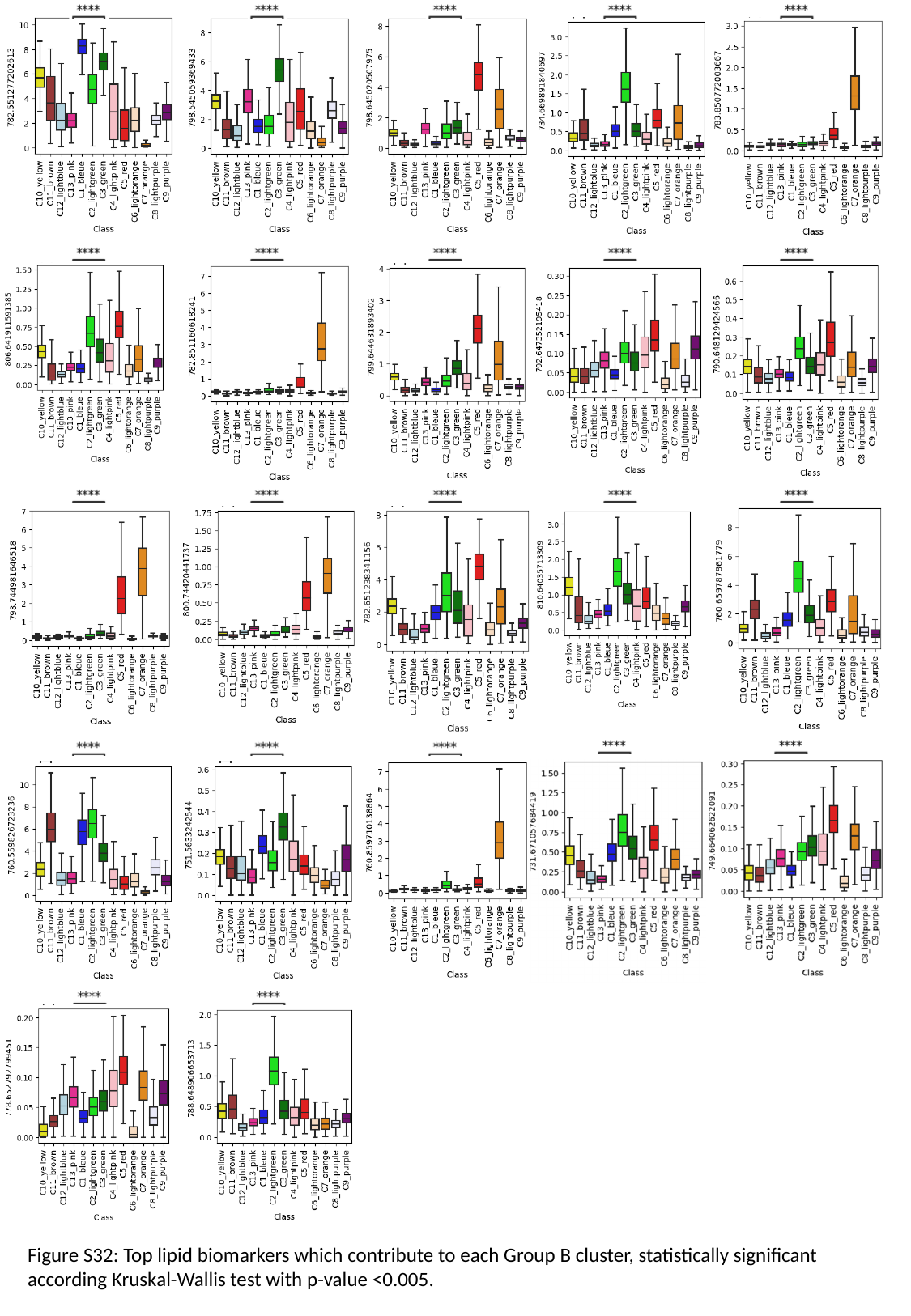

Figure S32: Top lipid biomarkers which contribute to each Group B cluster, statistically significant according Kruskal-Wallis test with p-value <0.005.

### Slide 48
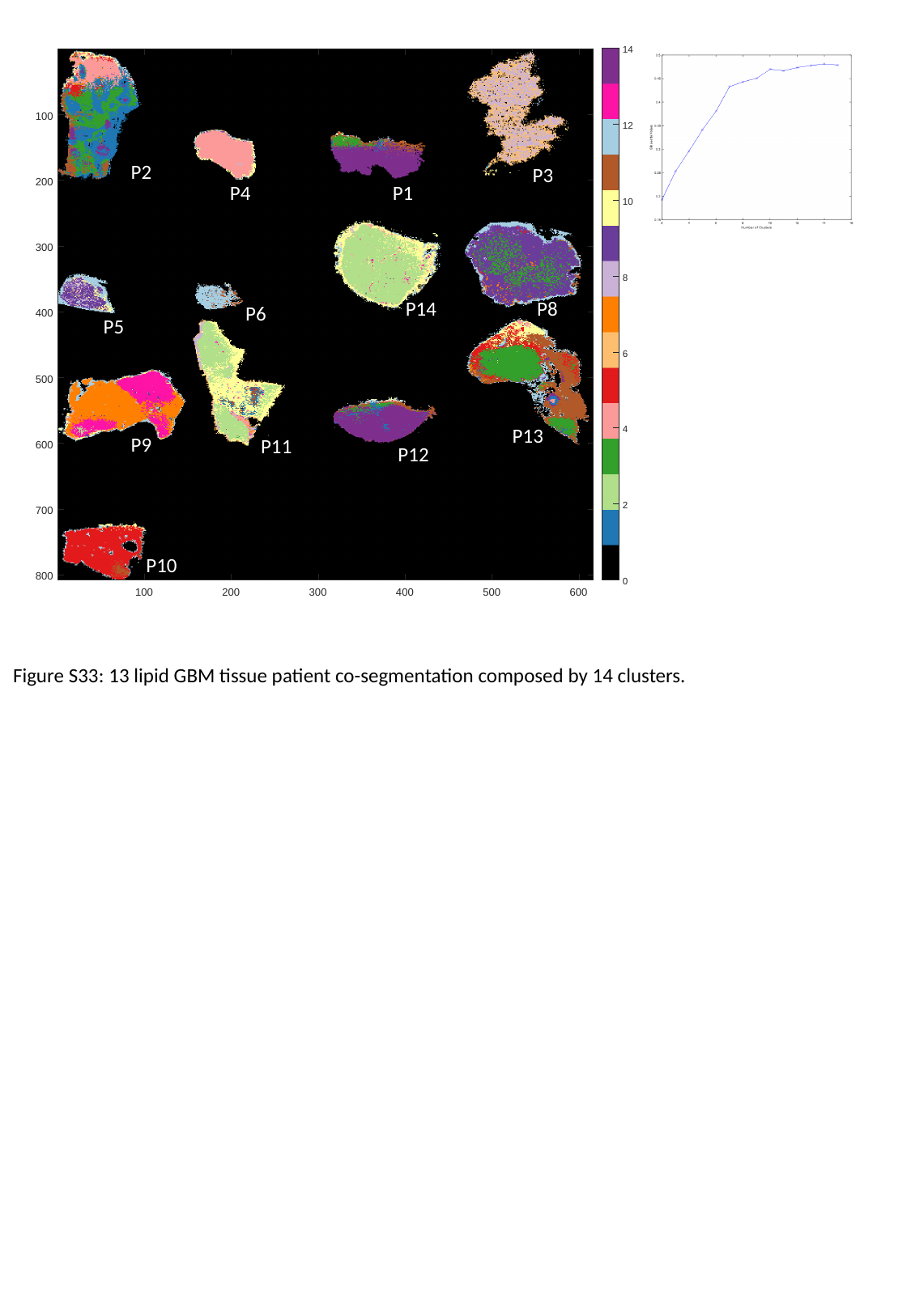

P2
P3
P4
P1
P14
P8
P6
P5
P13
P9
P11
P12
P10
Figure S33: 13 lipid GBM tissue patient co-segmentation composed by 14 clusters.

### Slide 49
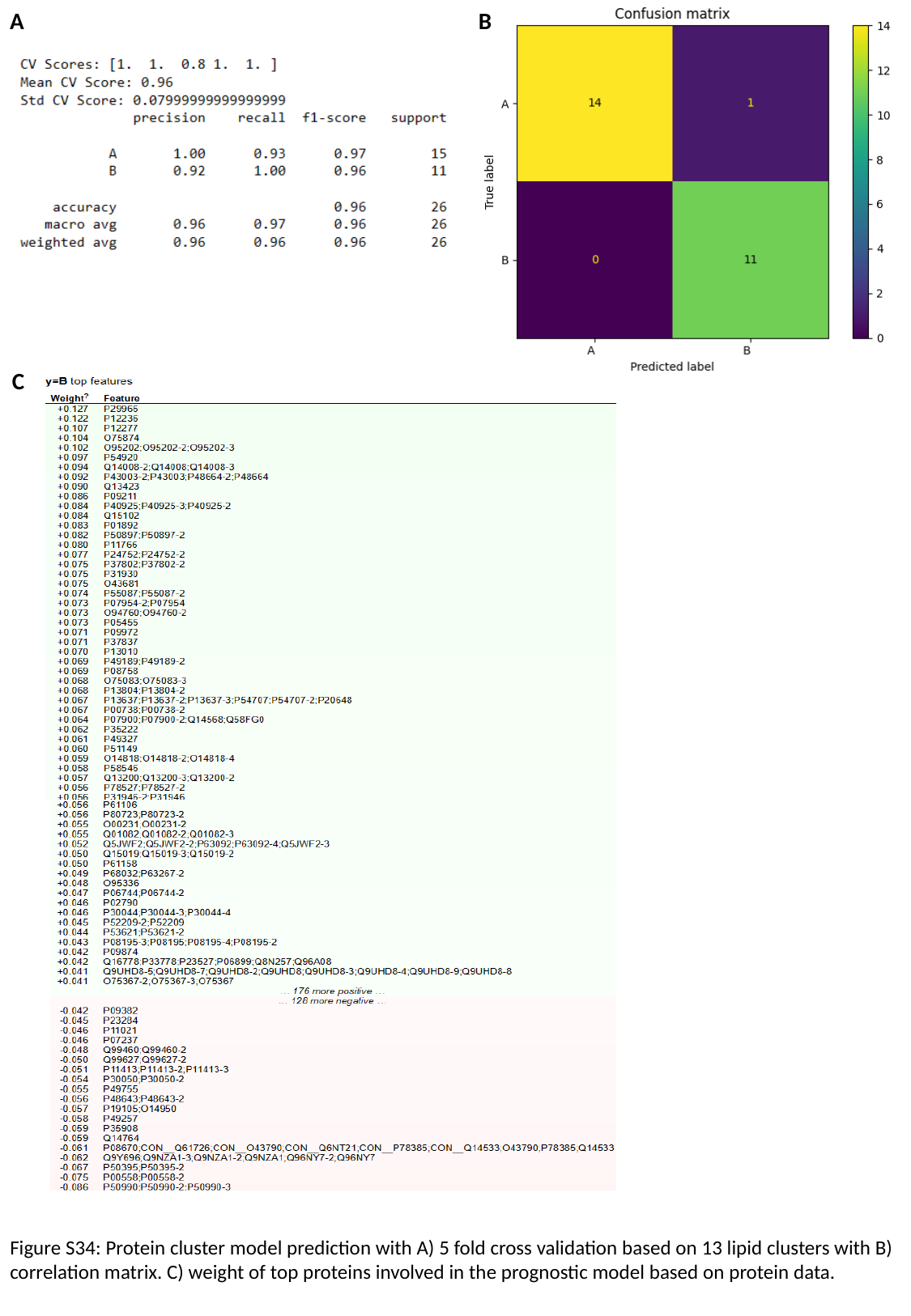

A
B
C
Figure S34: Protein cluster model prediction with A) 5 fold cross validation based on 13 lipid clusters with B) correlation matrix. C) weight of top proteins involved in the prognostic model based on protein data.

### Slide 50
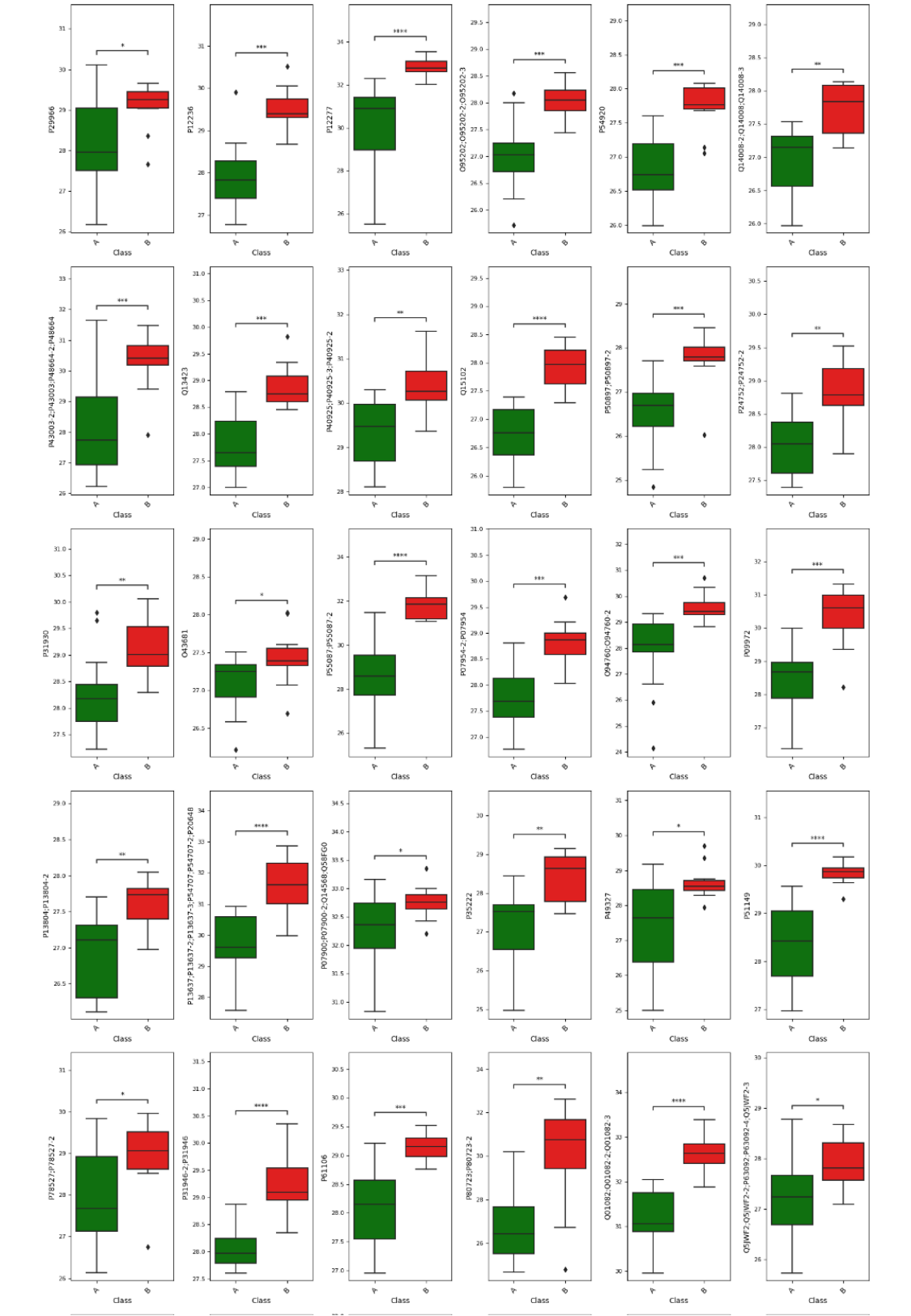

### Slide 51
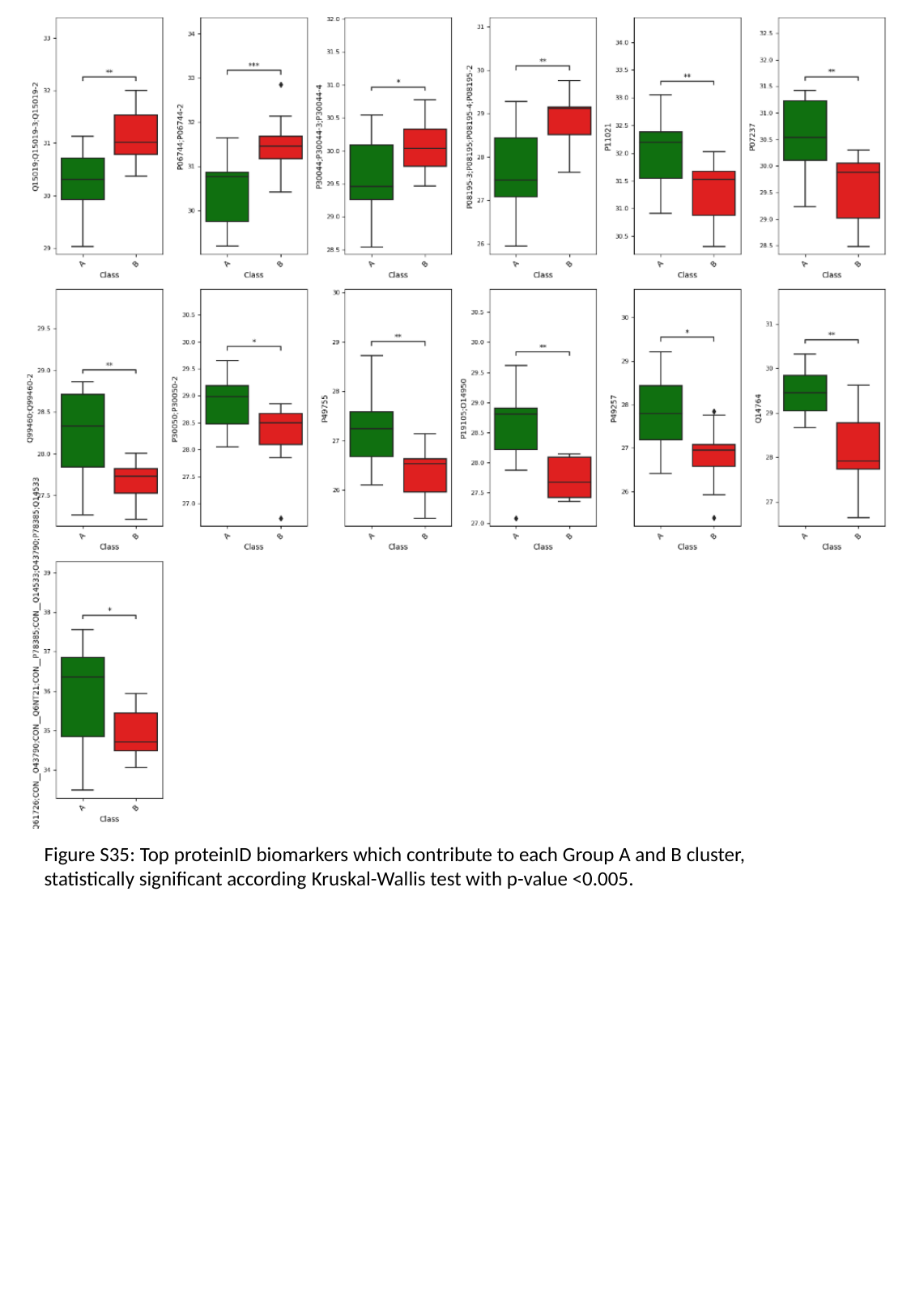

Figure S35: Top proteinID biomarkers which contribute to each Group A and B cluster, statistically significant according Kruskal-Wallis test with p-value <0.005.

### Slide 52
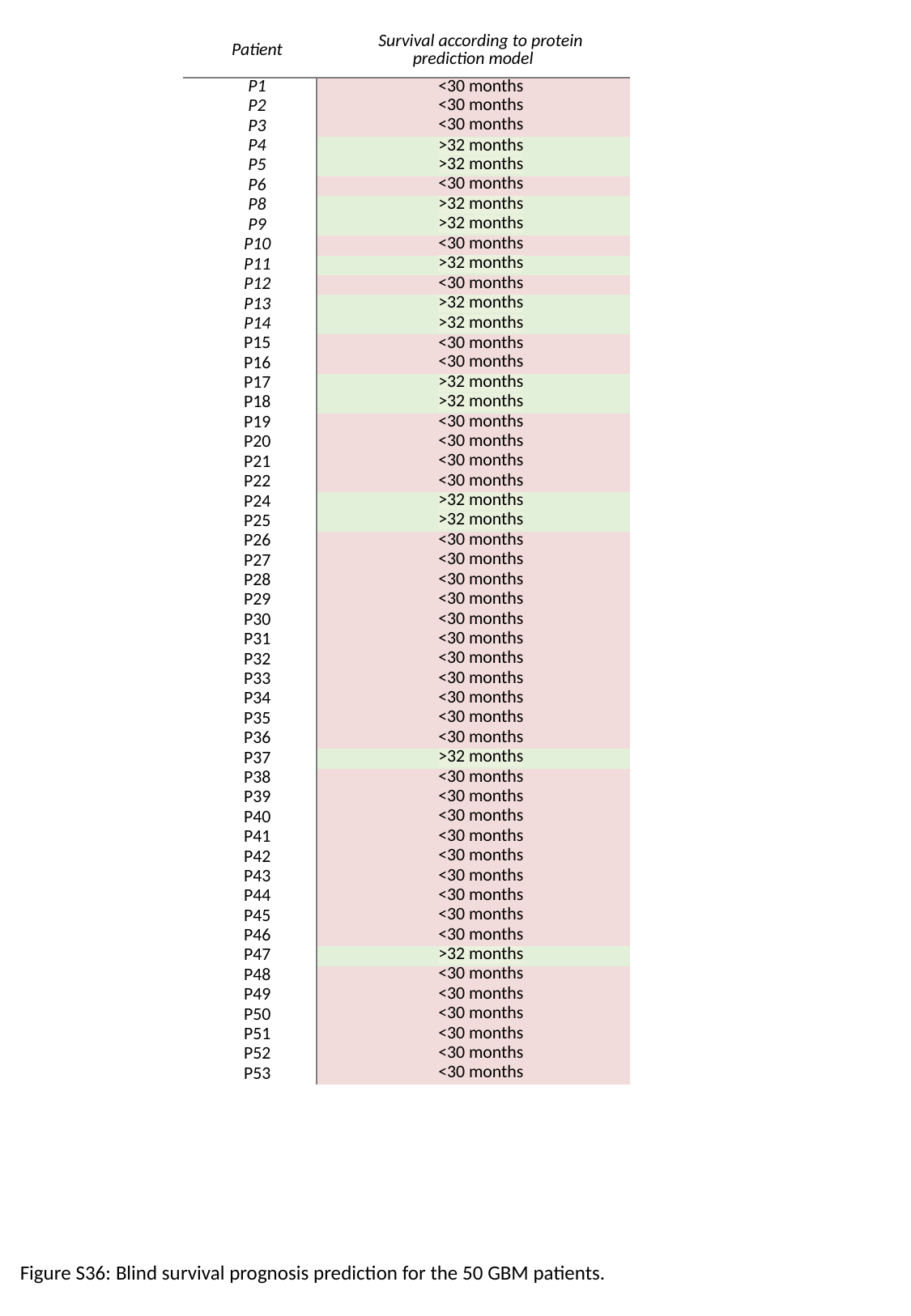

| Patient | Survival according to protein prediction model |
| --- | --- |
| P1 | <30 months |
| P2 | <30 months |
| P3 | <30 months |
| P4 | >32 months |
| P5 | >32 months |
| P6 | <30 months |
| P8 | >32 months |
| P9 | >32 months |
| P10 | <30 months |
| P11 | >32 months |
| P12 | <30 months |
| P13 | >32 months |
| P14 | >32 months |
| P15 | <30 months |
| P16 | <30 months |
| P17 | >32 months |
| P18 | >32 months |
| P19 | <30 months |
| P20 | <30 months |
| P21 | <30 months |
| P22 | <30 months |
| P24 | >32 months |
| P25 | >32 months |
| P26 | <30 months |
| P27 | <30 months |
| P28 | <30 months |
| P29 | <30 months |
| P30 | <30 months |
| P31 | <30 months |
| P32 | <30 months |
| P33 | <30 months |
| P34 | <30 months |
| P35 | <30 months |
| P36 | <30 months |
| P37 | >32 months |
| P38 | <30 months |
| P39 | <30 months |
| P40 | <30 months |
| P41 | <30 months |
| P42 | <30 months |
| P43 | <30 months |
| P44 | <30 months |
| P45 | <30 months |
| P46 | <30 months |
| P47 | >32 months |
| P48 | <30 months |
| P49 | <30 months |
| P50 | <30 months |
| P51 | <30 months |
| P52 | <30 months |
| P53 | <30 months |
Figure S36: Blind survival prognosis prediction for the 50 GBM patients.

### Slide 53
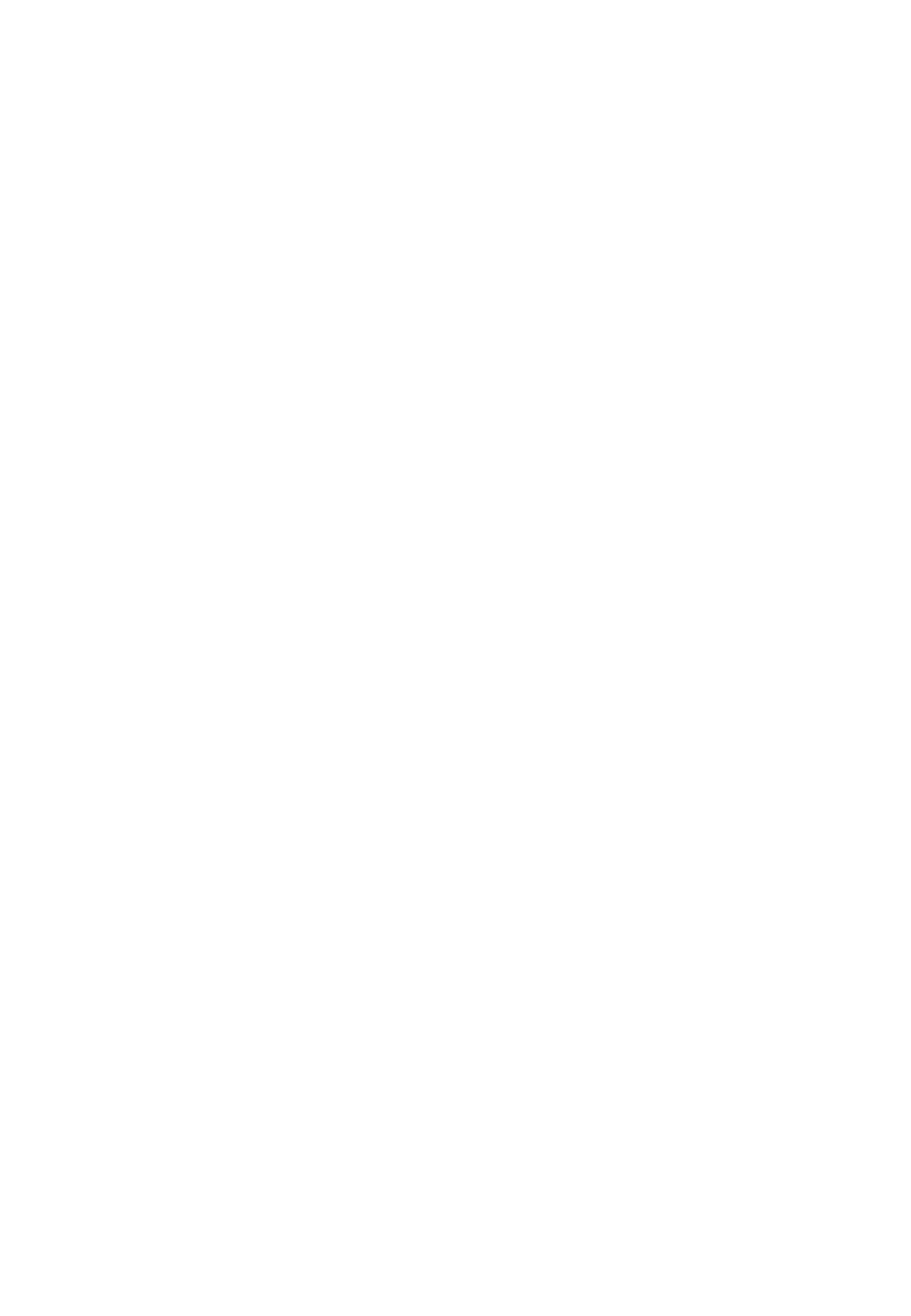
