## Supplementary material for "Heterogeneity Assessment and Protein Pathway Prediction via Spatial Lipidomic and Proteomic Correlation: Advancing Dry Proteomics concept for Human Glioblastoma Prognosis": Supplemntary Spreadsheet Legend

*Co-corresponding

**SUPPLEMENTARY SPREADSHEETS LEGENDS**

**Supplementary Spreadsheets S1:** List of proteins identified in molecular layer, white matter and granular layer rat brain cerebellum according spatial proteomic experiments. Proteins were annotated through their protein group, protein ids, protein name, genes, first protein description from uniport rat data base, as well as number of unique peptides assigned to each protein and protein sequence % coverage.

**Supplementary Spreadsheets S2:** List of exclusive proteins present in molecular layer, white matter and granular layer rat brain cerebellum according spatial proteomic experiments. Proteins were annotated through their protein group, protein ids, protein name, genes, a first protein description from uniport rat data base.

**Supplementary Spreadsheets S3**: Significant proteins involved in molecular layer, white matter and granular layer rat brain cerebellum biological triplicates after ANOVA test (p-value < 0.01) on spatial proteomic data. Proteins were annotated through their protein group, protein ids, protein name, genes, and first protein description from uniport rat data base.

**Supplementary Spreadsheets S4:** List of proteins identified in horizontal rat brain section clusters according spatial proteomic experiments. Proteins were annotated through their protein group, protein ids, protein name, genes, first protein description from uniport rat data base, as well as number of unique peptides assigned to each protein and protein sequence % coverage.

**Supplementary Spreadsheets S5**: ANOVA test (p-value < 0.01) significant proteins involved in rat brain horizontal section clusters from biological triplicate spatial proteomic data. Proteins were annotated through their protein group, protein ids, protein name, genes, and first protein description from uniport rat data base.

**Supplementary Spreadsheets S6:** Common protein identified according current study and Delcourt, V. *et al.*, 2018.in hippocampus and corpus callosum rat brain

**Supplementary Spreadsheets S7:** List of proteins identified in Glioblastoma cohort according spatial proteomic experiments. Proteins were annotated through their protein group, protein ids, protein name, genes, first protein description from uniport rat data base, as well as number of unique peptides assigned to each protein and protein sequence % coverage.

**Supplementary Spreadsheets S8:** Proteins involved in GBM classification group A and group B. Proteins were annotated through their protein group, protein ids, protein name, and genes.
