## Supplementary figures and images for "Heterogeneity Assessment and Protein Pathway Prediction via Spatial Lipidomic and Proteomic Correlation: Advancing Dry Proteomics concept for Human Glioblastoma Prognosis"

### Graphical Abstract

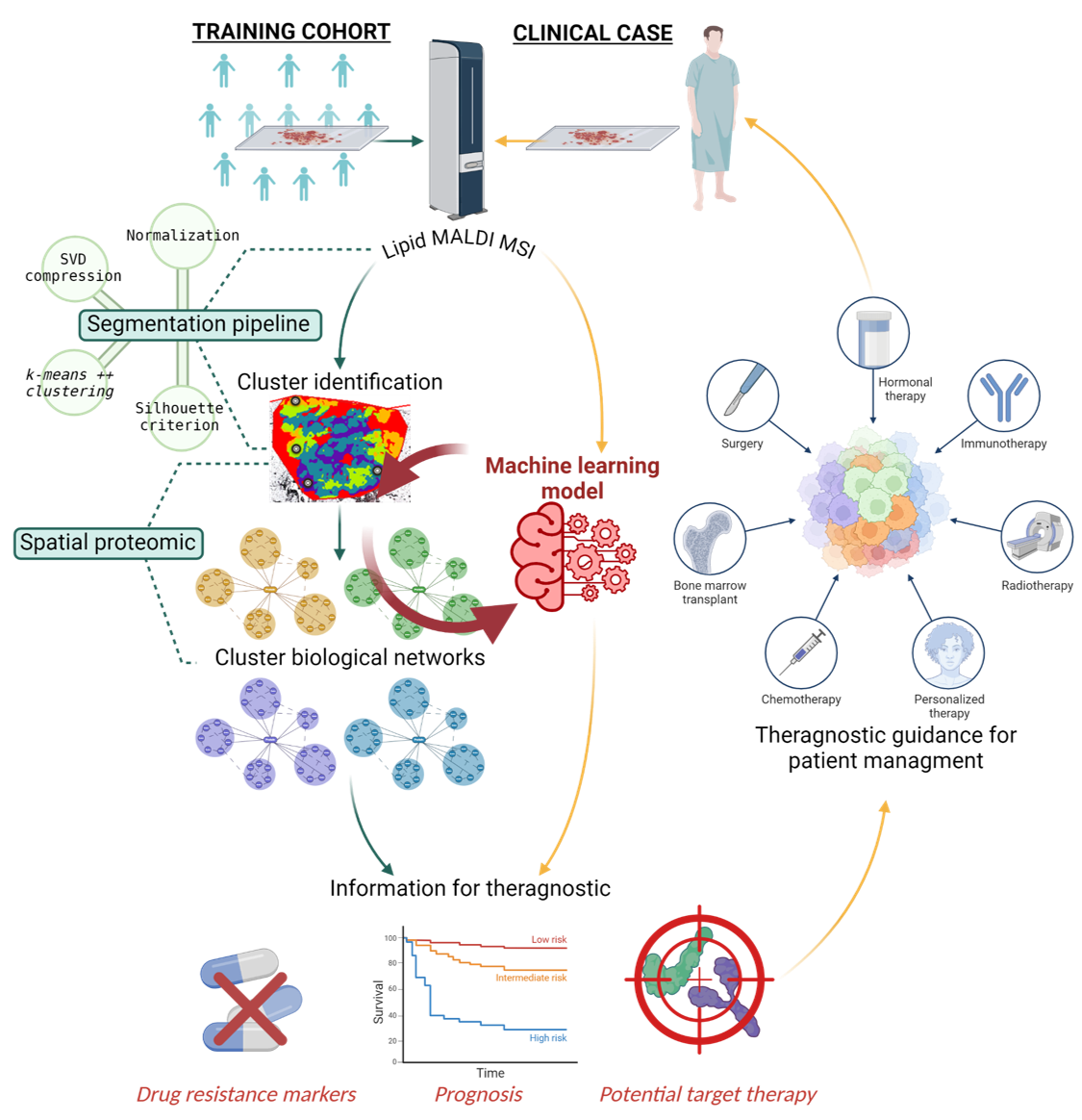
